# Genome-wide mapping of therapeutically-relevant SARS-CoV-2 RNA structures

**DOI:** 10.1101/2020.06.15.151647

**Authors:** Ilaria Manfredonia, Chandran Nithin, Almudena Ponce-Salvatierra, Pritha Ghosh, Tomasz K. Wirecki, Tycho Marinus, Natacha S. Ogando, Eric J. Snider, Martijn J. van Hemert, Janusz M. Bujnicki, Danny Incarnato

**Author notes:** Correspondence: Danny Incarnato, Janusz Bujnicki, Martijn J. van Hemert. These authors contributed equally to this work (Ilaria Manfredonia, Chandran Nithin, Almudena Ponce-Salvatierra, Pritha Ghosh, Tomasz K. Wirecki).

## Abstract

SARS-CoV-2 is a betacoronavirus with a linear single-stranded, positive-sense RNA genome of ∼30 kb, whose outbreak caused the still ongoing COVID-19 pandemic. The ability of coronaviruses to rapidly evolve, adapt, and cross species barriers makes the development of effective and durable therapeutic strategies a challenging and urgent need. As for other RNA viruses, genomic RNA structures are expected to play crucial roles in several steps of the coronavirus replication cycle. Despite this, only a handful of functionally conserved structural elements within coronavirus RNA genomes have been identified to date.

Here, we performed RNA structure probing by SHAPE-MaP to obtain a single-base resolution secondary structure map of the full SARS-CoV-2 coronavirus genome. The SHAPE-MaP probing data recapitulate the previously described coronavirus RNA elements (5′ UTR, ribosomal frameshifting element, and 3′ UTR), and reveal new structures. Secondary structure-restrained 3D modeling of highly-structured regions across the SARS-CoV-2 genome allowed for the identification of several putative druggable pockets. Furthermore, ∼8% of the identified structure elements show significant covariation among SARS-CoV-2 and other coronaviruses, hinting at their functionally-conserved role. In addition, we identify a set of persistently single-stranded regions having high sequence conservation, suitable for the development of antisense oligonucleotide therapeutics.

Collectively, our work lays the foundation for the development of innovative RNA-targeted therapeutic strategies to fight SARS-related infections.

## Introduction

RNA viruses encode the information needed to take control of the host cell on two levels (Boerneke et al., 2019). On one hand, the linear sequence of their RNA genomes encodes for all the proteins needed to take over the host cell machinery and to assemble new viral particles. On the other hand, their single-stranded RNA genome folds back on itself to form intricate secondary and tertiary structures that have been proven essential for viral replication, protein synthesis, packaging, immune system evasion, and more. RNA viruses, that are responsible for numerous deadly diseases (e.g. AIDS, Hepatitis C, SARS, Dengue, and Ebola), are characterized by higher mutation rates compared to DNA viruses, enabling them to rapidly evolve and adapt (Sanjuán et al., 2010). As a consequence of their high mutation rates, RNA viruses can rapidly develop resistance towards drugs and vaccines by slightly altering their core proteins (Irwin et al., 2016). In contrast, certain RNA structures formed in the context of viral RNA genomes are well conserved (Boerneke et al., 2019), in spite of changes in the underlying encoded amino acid sequence, making them valuable therapeutic targets.

Coronaviruses (CoV) are positive-sense, single-stranded RNA viruses, members of the *Coronaviridae* family (Cui et al., 2018). This family consists of four genera, of which two (alpha and beta) can only infect mammals, while the other two (gamma and delta) mostly infect birds, although some of them can also infect mammals. These viruses were not anticipated to be highly pathogenic in humans until the outbreak of the *severe acute respiratory syndrome* coronavirus (SARS-CoV) in 2002 (Wit et al., 2016). Since then, two other major outbreaks of coronaviruses occurred, one by the *Middle East respiratory syndrome* coronavirus (MERS-CoV) in 2012 and recently by the new SARS-related coronavirus SARS-CoV-2 (also known as 2019-nCoV) at the end of 2019 (Zhou et al., 2020). The latter, still ongoing at the time of writing this article, rapidly resulted in a pandemic, with (to date) nearly 8 million people infected and over 400,000 deaths across the world. The ability to evolve inside different reservoirs and to cross species barriers, infecting humans with high morbidity and mortality, makes this genus a recurrent potential threat for worldwide public health. Indeed, certain SARS-CoV-like viruses from bats have been previously shown to be able to infect human cells without the need for any prior adaptation; thus, suggesting that SARS-related outbreaks can potentially re-emerge at any time (Wit et al., 2016).

In light of these considerations, identifying new therapeutically-relevant and durable druggable targets for the treatment of coronavirus infections constitutes a key and highly-timely need. In this perspective, RNA structural elements represent attractive targets for drug discovery. Indeed, inhibition of viral replication by RNA-targeting small molecule drugs has already been proven to be feasible for other RNA viruses, such as the human immunodeficiency virus (HIV), hepatitis C virus (HCV), SARS-CoV, and influenza A virus (IAV; (Hermann, 2016). Additionally, the identification of highly-conserved weakly structured regions within viral RNA genomes might aid the design of oligonucleotide-bases antiviral therapeutics.

Coronaviruses bear the largest genomes among RNA viruses, ranging from approximately 26 to 32 kb. A handful of functional cis-regulatory RNA structural elements have been previously identified by phylogenetic analyses and these include structures in the 5′ and 3′ untranslated regions (UTRs) and the ribosomal frameshifting element (FSE; (Madhugiri et al., 2016, 2018; Yang and Leibowitz, 2015). In the 5′ UTR of most betacoronaviruses (Beta-CoV), several stem-loops (SL1-5) involved in mediating viral replication have been identified. The ORF1a-ORF1b boundary hosts the frameshifting element (FSE; (Brierley et al., 1989). This pseudoknotted structure is pivotal for the programmed ribosomal frameshifting, that allows the translation of the ORF1ab polyprotein. A second putative pseudoknot has been proposed to be located in the 3′ UTR (Goebel et al., 2003; Stammler et al., 2011), that includes an extremely conserved octanucleotide within the hypervariable region (HVR; (Goebel et al., 2006), and the stem-loop-like 2 motif (s2m; (Robertson et al., 2004).

Sharing over 79.6% sequence identity to the genome of SARS-CoV (Zhou et al., 2020), SARS-CoV-2 has been predicted to harbor these typical RNA structure elements (Rangan et al., 2020). However, the structural complexity of SARS-CoV-2 and other Beta-CoV genomes has remained largely unexplored so far, probably because of the challenges connected with the structural analysis of RNA molecules of such a big size. Deep comprehension of the structural architecture of the SARS-CoV-2 RNA genome is crucial to identify new key (un)structured elements for the development of innovative therapeutic approaches.

To address this need, in this work we provide the first experimental characterization of the full-length genome of a coronavirus by SHAPE mutational profiling (SHAPE-MaP) analysis (Siegfried et al., 2014), using the novel SARS-CoV-2 virus as a model. After modeling the secondary structure of the entire RNA genome, we identified a subset of regions showing a low propensity towards folding, ideal for the design of antisense oligonucleotide (ASO) therapeutics, as well as a set of stably folded RNA structures, of which at least ∼8% are under selective pressure and show significant covariation. By coupling secondary structure constraints and coarse-grained 3D modeling, we then infer the 3D structure of these elements and identify the most suitable for accommodating the interaction with small molecule drugs. Collectively, our work provides the cornerstone for the development of RNA-targeted therapeutic strategies to fight SARS-related infections.

## Results and Discussion

To determine the secondary structure landscape of SARS-CoV-2, we devised a multiplex PCR strategy (see Methods) to perform the targeted SHAPE mutational profiling (SHAPE-MaP) analysis of the entire viral genome (∼30 kb). First, we performed *in vitro* refolding of total RNA from Vero E6 cells infected with SARS-CoV-2, followed by probing with 2-methylnicotinic acid imidazolide (NAI; Figure 1A). This approach has a considerable advantage, compared to *in vitro* transcription, of allowing the analysis of high amounts of full-length SARS-CoV-2 RNA, bearing post-transcriptional RNA modifications that could occur *in vivo* during natural infection, and that might affect local RNA folding.

**Figure 1.**
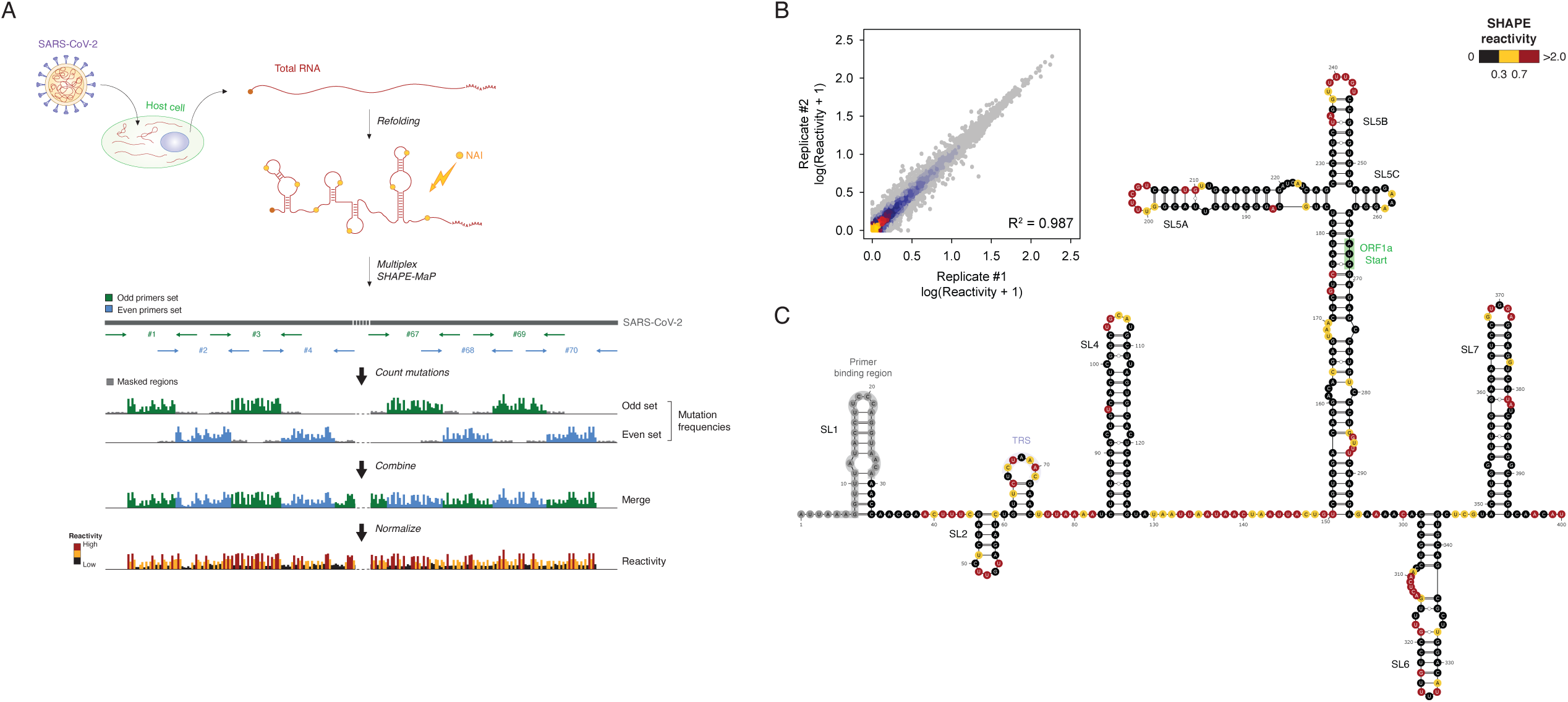
Genome-wide SHAPE-MaP analysis of SARS-CoV-2. (**A**) Schematic of the multiplex targeted SHAPE-MaP approach for querying the SARS-CoV-2 genome from total RNA. (**B**) Heat scatter plot showing the correlation between SHAPE-MaP reactivities for the two replicates (*R*^*2*^ = 0.987). (**C**) Measured SHAPE-MaP reactivities superimposed on the reference Sarbecovirus 5′ UTR structure.

We designed a set of 70 primer pairs, divided into two terminally-overlapping subsets (odd and even), to perform the targeted SHAPE-MaP analysis in ∼500 bp tiles. The terminal overlap between the two sets was introduced to query those bases that would otherwise be blind to SHAPE-MaP analysis because of primer binding. Following the equimolar pooling of amplicons and fragmentation, DNA libraries from two biological replicates of SHAPE-probed RNA were prepared and sequenced on the Illumina platform. The mutational signature for the whole SARS-CoV-2 genome was then obtained by merging the odd and even sets, after computationally masking the respective primer-pairing regions. Correlation of replicates was very high (*p* = 0.97 for both odd and even set on raw mutation frequencies; *p* = 0.99 genome-wide after normalization; Figure 1B and Figure S1). Given this, we combined both replicates for secondary structure modeling.

Overall, we sequenced over 30 million high-quality reads per replicate, of which >99% were successfully aligned to the SARS-CoV-2 genome, resulting in a total median per-base coverage >100,000X, with ∼99.9% of the genome being covered at least 1X. As accurate structure inference requires a reliable mutational signal, which in turn depends on the sequencing depth, we imposed a minimum coverage threshold of 1,000X. This filter retained over 96.5% of the genome. We observed exceptional agreement between known Beta-CoV secondary structure motifs and our SHAPE-MaP reactivities (Figure 1C and Figure S2), with on average >76% of highly reactive bases (≥0.7) being annotated as single-stranded. When including also terminal base-pairs (adjacent to helix termini or bulges/loops), the measured agreement rose up to ∼90%.

Next, we used SHAPE-derived constraints to model the secondary structure of the SARS-CoV-2 genome (see Methods). This approach has been extensively validated and previously applied by us and other groups to successfully map viral RNA structures (Siegfried et al., 2014; Simon et al., 2019). To convert NAI reactivities into pseudo-free energy contributions, a complementary SHAPE-MaP dataset was produced by probing *E. coli* rRNAs with NAI under *ex vivo* deproteinized conditions and determined optimal slope and intercept to be 0.8 and -0.2 by grid search, respectively (Figure S3). By incorporating these parameters into our RNA Framework algorithm for SHAPE-directed structure prediction (Incarnato et al., 2018), we have obtained a high-quality single-base resolution secondary structure map of the SARS-CoV-2 RNA genome (Figure 2).

**Figure 2.**
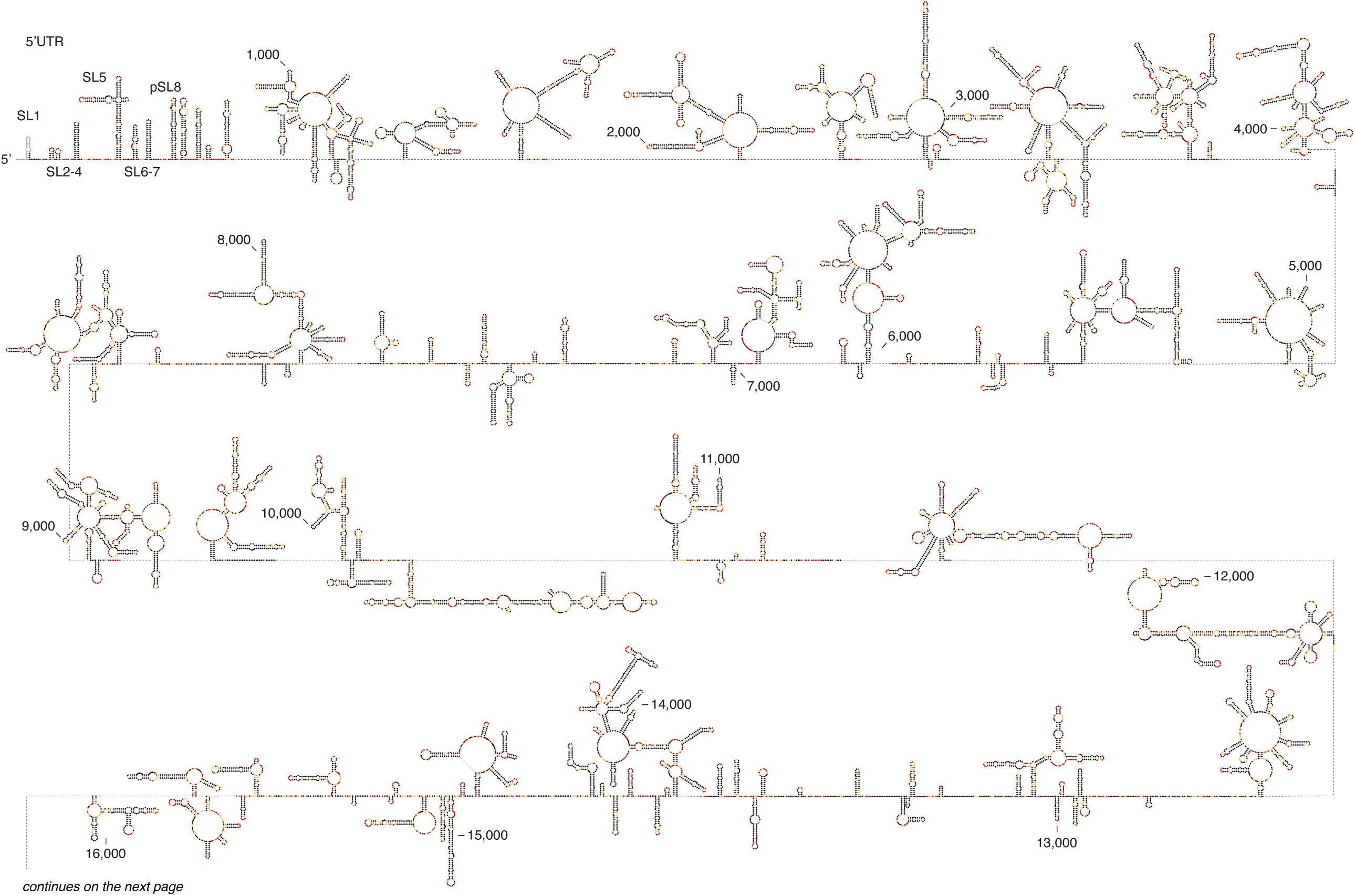
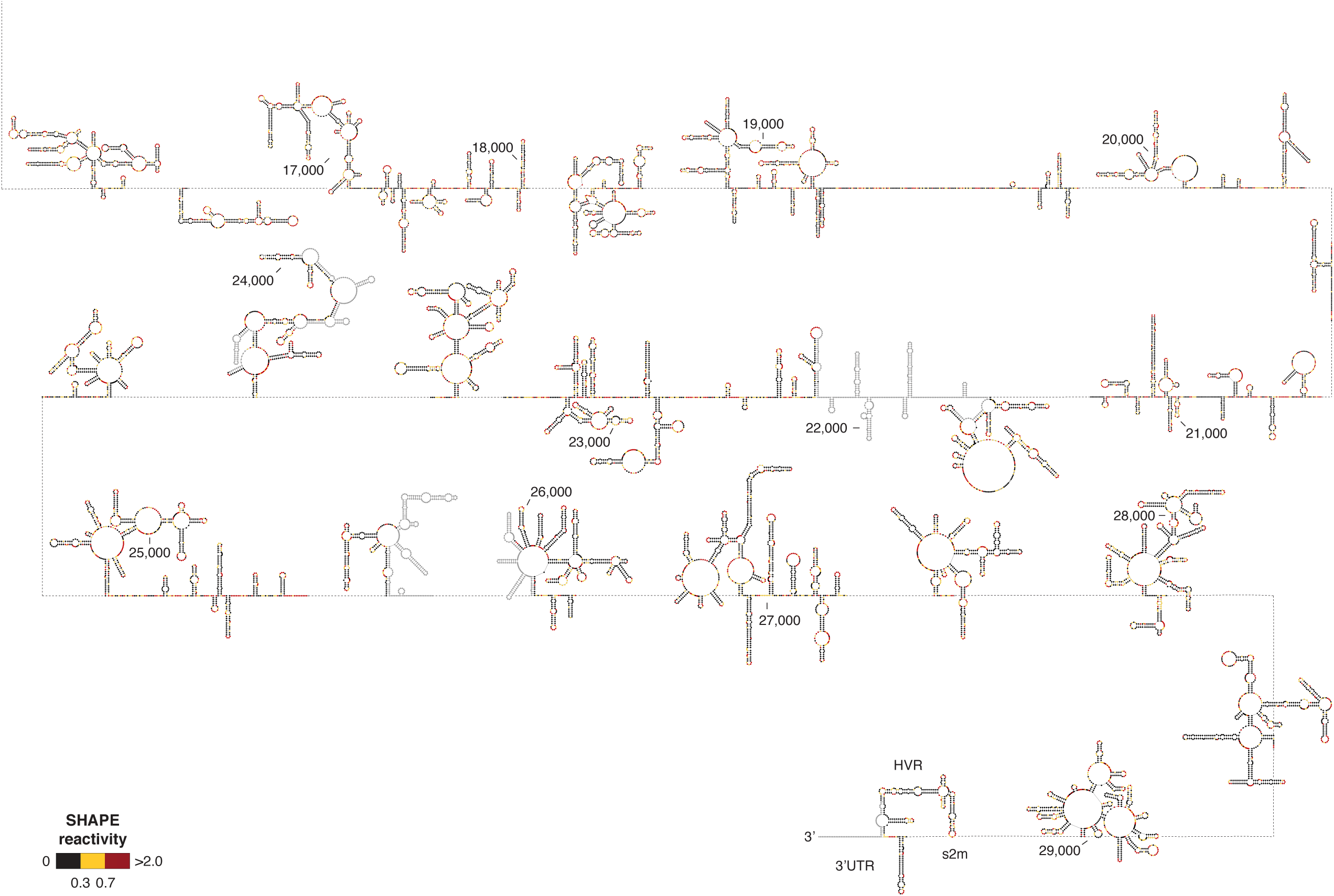
Structure map of the full SARS-CoV-2 genome. Maximum expected accuracy (MEA) structure representation of the SARS-CoV-2 genome. Bases are color-coded according to SHAPE-MaP reactivities. Bases with coverage lower than 1000X are marked in grey. The approximate genomic position is indicated every 1000 nt. Structure was plotted with VARNA (Darty et al., 2009).

From the first inspection of our model, it appeared that roughly one half (∼55.5%) of the SARS-CoV-2 genome is structured, in agreement with the observed fraction of low reactive residues as determined by SHAPE probing (∼57.8% of reactivities ≤0.3). Comparison of the distribution of base reactivities for unpaired, internally-paired, and terminally-paired residues showed a pattern comparable to that of E. coli ribosomal RNAs (Figure S4), hence supporting the quality of our SARS-CoV-2 RNA structure model. Previously described RNA structure elements typical of Sarbecoviruses and other Beta-CoV, such as 5′ UTR helices S1 to S7, the ribosomal frameshifting element (FSE) and the 3′ UTR stem-loop II-like motif (s2m) were all successfully modeled in the context of the full genome, without the need to impose any hard constraints, further supporting the high accuracy and reliability of the SARS-CoV-2 genome structure model proposed here. Interestingly, the recently proposed stem-loops SL8 to SL10 (positions: ∼400-450 nt; Rangan et al., 2020) are not supported by our data. Instead, the region spanning these three short hairpins appears to be involved in the formation of a large stem-loop-like structure, spanning nucleotides 407 to 478 (putative stem-loop 8, pSL8).

We next analyzed the SARS-CoV-2 genome structure using a previously defined metric, the Shannon entropy (Siegfried et al., 2014). The Shannon entropy provides an estimate of the likelihood of a given RNA region to fold into multiple conformations. Therefore, we determined the Shannon entropy across the whole SARS-CoV-2 genome and combined this measure with SHAPE data to identify two sets of regions: regions of persistent single-strandedness (low Shannon - high SHAPE) and regions stably folding into well-defined conformations (low Shannon - low SHAPE).

Regions of persistent single-strandedness are unlikely to form any structure (or to be involved in intramolecular base-pairing, in general), thus representing preferred targets for the design of antisense oligonucleotide (ASO) therapeutics. Our analysis (see Methods) identified 70 such regions, at least 25 nucleotides long, spanning a total of 2,910 bases (∼9.7% of the SARS-CoV-2 genome) matching these criteria (Figure S5 and Table S2). We next evaluated the sequence conservation of these regions, by multiple sequencing alignment with different sets of coronavirus sequences (Figure S6), specifically: 1) SARS-CoV; 2) MERS-CoV; 3) other Beta-CoV; 4) all other CoV (alpha, gamma, and delta). Notably, ∼23.9% of these regions, accounting for roughly 711 bases (∼2.4% of the SARS-CoV-2 genome), showed over 70% conservation in SARS-CoV (average ∼81.3%) and over 50% conservation in the other three datasets (average ∼63.5% for MERS-CoV, ∼60.4% for other Beta-CoV, and ∼53% for all other CoV). Particularly, five regions (accounting for a total of 240 bases) showed exceptionally high conservation in all the analyzed alignments (average SARS-CoV: 86.4%; MERS-CoV: 78.6%; other Beta-CoV: 74.2%; all other CoV: 68.3%), suggesting that some essential regulatory RNA element might reside here and be under strong selection.

Next, we focused on low Shannon - low SHAPE regions (Figure 3). In total, we identified 149 structure elements having both low SHAPE reactivities, reflecting low nucleotide dynamics (possibly due to base-pairing), and low Shannon entropies, reflecting high structural stability, thus the presence of a predominant RNA structural conformation. Overall, these regions encompassed ∼38.6% of the SARS-CoV-2 genome (11,547 bases) and included the 5′ and 3′ UTRs. These regions are likely to be involved in the mechanistic and structural aspects of the viral RNA function. Previous computational analyses have shown that complex structured RNA molecules often exhibit pockets with physical properties suitable for small molecule binding, which can be identified in RNA 3D structure models (Hewitt et al., 2019). Thus, the discovery of multiple structured regions in SARS-CoV-2 genomic RNA offered an opportunity to test their potential to be targeted by small molecules.

**Figure 3.**
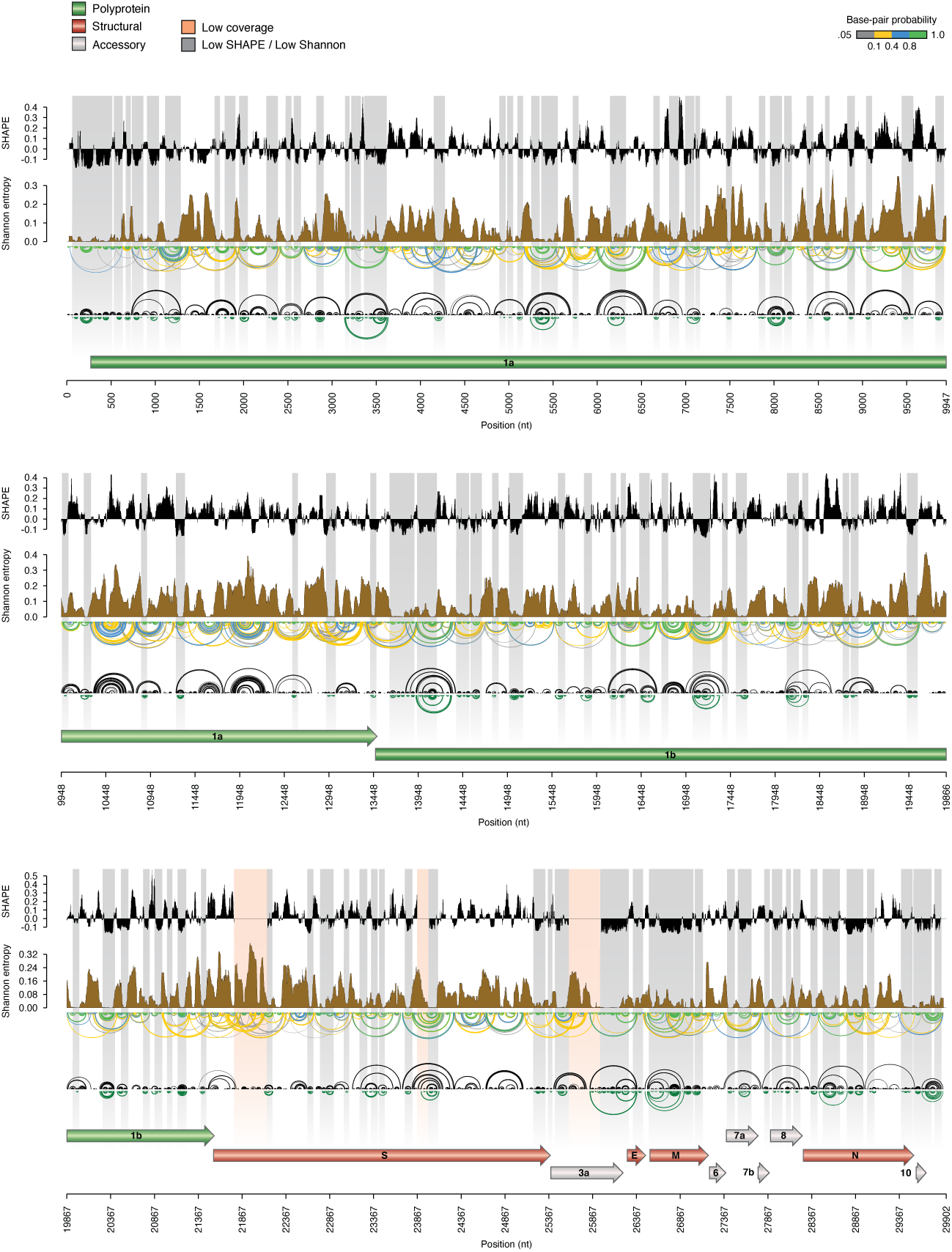
Identification of RNA regions with well-defined foldings. Map of the SARS-CoV-2 genome depicting (top to bottom): median SHAPE reactivity (in 50 nt centered widows, with respect to the median reactivity across the whole genome), Shannon entropy, base-pairing probabilities, maximum expected accuracy (MEA) structure. Regions with both low Shannon and low SHAPE, likely identifying RNA regions with well-defined foldings, are marked in grey and depicted as green arcs below the MEA structure. Three regions with sequencing coverage below 1,000X are marked in red.

Starting from the set of low Shannon - low SHAPE structures, shorter contiguous helices, that were likely to interact with each other’s, were combined to yield a total of 70 RNA segments. Then, by using the SimRNA method (Boniecki et al., 2015), we modeled their 3D folding. We used secondary structure information derived from the SHAPE-MaP experiments to restrain the folding trajectories that sampled the 3D conformational space in search of local and global energy minima. For each 3D-folded segment, we identified clusters of the most commonly occurring 3D conformations, and we generated 1000 lowest-energy models to represent the most stable 3D conformations. In general, while the secondary structures agreed very well with that inferred from experimental probing, the 3D structures exhibited significant overall conformational variability, which was expected given the elongated character of secondary structures inferred from experimental data. Nonetheless, the 3D modeling revealed a number of persistent complex spatial motifs, often comprising junctions and structured bulges and loops.

Next, we used the Fpocket software (Guilloux et al., 2009) to score the putative druggable sites in 1000 lowest-energy models for each RNA segment, based on the criteria we established on a set of experimentally determined 3D structures of small viral RNAs complexed with small molecule ligands (Figure S7; see Methods). We mapped the predicted druggable pockets on the RNA sequence to identify clusters of residues that formed potential ligand-binding sites in the largest fraction of best-energy models (Figures S8-S57). The majority of targetable pockets are found in high-information-content structures such as multiway junctions (Figures 4 and Figures S58-S59; Warner et al., 2018), although they are also very often found in bulges. On the other hand, terminal loops and symmetrical bulges were rarely identified as druggable sites in SARS-CoV-2 RNA segments.

**Figure 4.**
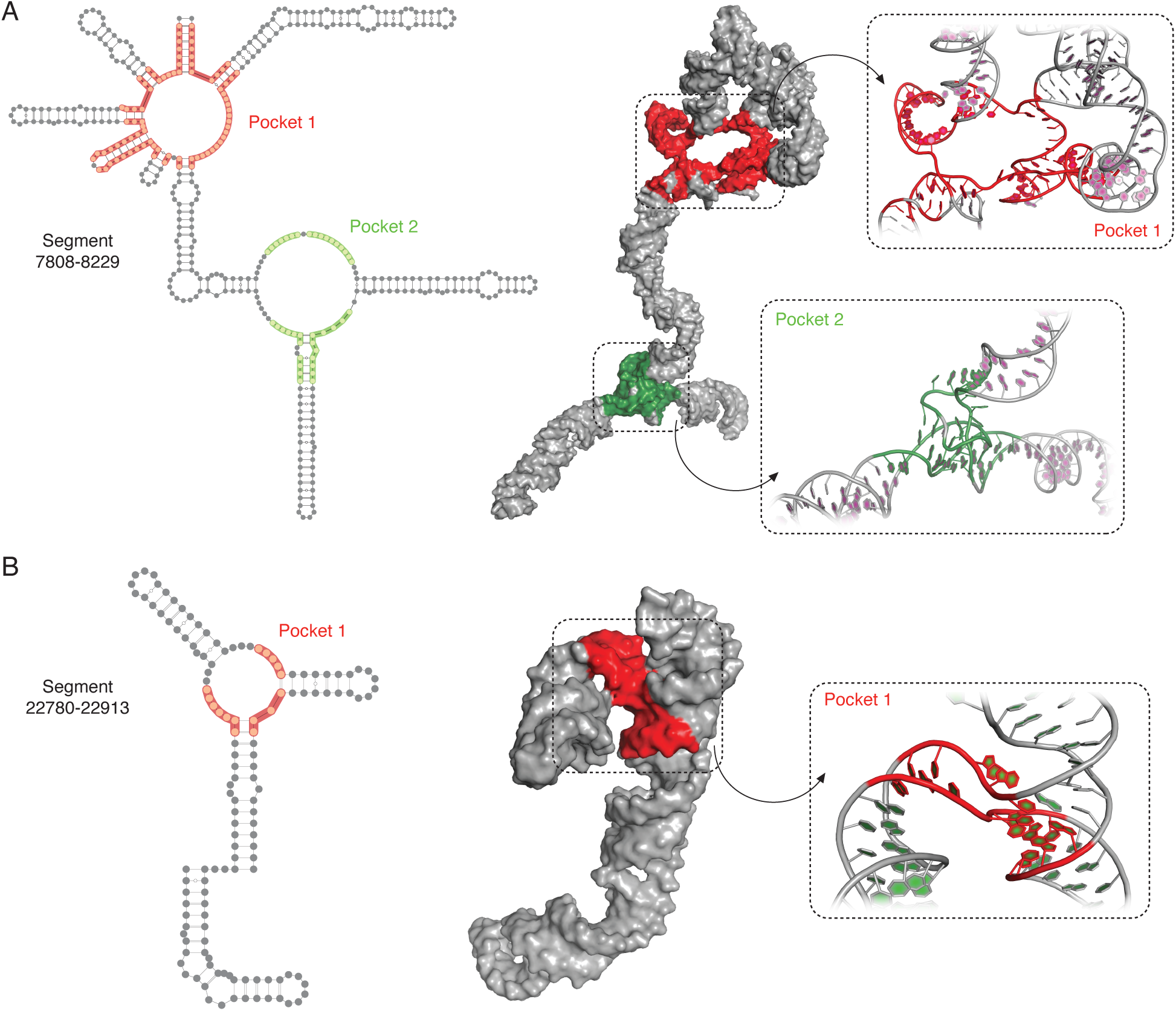
3D modeling of SARS-CoV-2 RNA structured segments and identification of druggable pockets. Consensus secondary structure for segment 7808-8229 (**A**), and segment 22780-22913 (**B**), as derived from the 1000 lowest energy 3D structure structures modeled by SimRNA (using the MEA secondary structure inferred from SHAPE-MaP as a restraint). Residues composing the identified druggable pockets are shaded. Different shading colors mark distinct pockets. The 3D model of the medoid structure from the most abundant cluster is also shown, with pockets colored as for the secondary structure. The insets show the atomic representation of the identified pockets.

We identified a druggable pocket at the base of the 3′ stem-loop II-like motif (s2m, found within the segment 29548-29870; Figure S57). The s2m is a highly conserved motif found at the 3′-UTRs of astrovirus, coronavirus, and equine rhinovirus genomes (Jonassen et al., 1998) and has been proposed to affect host translation by either interacting with ribosomal proteins or by getting processed into a mature microRNA (Robertson et al., 2004; Tengs et al., 2013). Our analyses also recovered a potentially druggable site within the FSE (found within the segment 13367-13546; Figure S27), which is crucial for viral fitness since it allows for the synthesis of functional and structural viral proteins (Brierley et al., 1989). The latter pocket encompasses a previously reported one, that could be targeted with a small molecule proven to successfully inhibit the frameshifting activity of SARS-CoV (Park et al., 2011).

To further prioritize the structured segments of SARS-CoV-2 RNA more likely to represent functional elements, we devised an automatic strategy for the identification of sequence segments showing significant base-pair covariation across coronaviruses (see Methods). By building a first covariance model (CM) using Infernal (Nawrocki and Eddy, 2013) and the sole SARS-CoV-2 sequence (together with the secondary structure inferred from SHAPE-MaP data), we searched (with a relaxed E-value cutoff) for homologous sequences in a non-redundant CoV database, discarding those matches in which less than 55% of the canonical base-pairs from the SARS-CoV-2 structure were supported. Furthermore, by taking advantage of the extremely conserved architecture of CoV genomes (Lauber et al., 2013), we further discarded those matches whose relative position in the genome changed by more than 3.5% with respect to that of SARS-CoV-2. The resulting alignment was then used to refine the CM and the whole procedure was repeated three times.

This analysis identified a total of 12 RNA elements (out of the originally selected 149 low Shannon - low SHAPE structures, ∼8%) for which at least 5% of the canonical base-pairs showed significant covariation (on average, 15.1% at E-value < 0.05, 18.6% at E-value < 0.1), as determined by the R-scape software (Rivas et al., 2016). Among these, our pipeline successfully recovered both the SL5 element of Sarbecovirus 5′ UTR and the 3′ UTR (Figure S60). Furthermore, we recovered a number of high-information-content structures for which the Fpocket analysis identified high-scoring, putative druggable pockets (Figure 5). These RNA structures show variable degrees of conservation across different CoV genera, with 6 out of 12 (50%) being conserved in both Alpha-CoV and Gamma-CoV, and 4 of them being conserved in Delta-CoV as well (Figure S61). It is however worth pointing out that, while the presence of covariation can support the presence of functional RNA structures, the lack of covariation due to high sequence conservation does not provide any statistical support for such conclusions. Therefore, although we did not find sufficient significant covariation support for the remaining structures (possibly also due to the stringency of our filtering criteria), many of them can anyway represent either recently evolved RNA structures or RNA cis-regulatory elements residing within regions whose sequence is constrained by the underlying amino-acid sequence.

**Figure 5.**
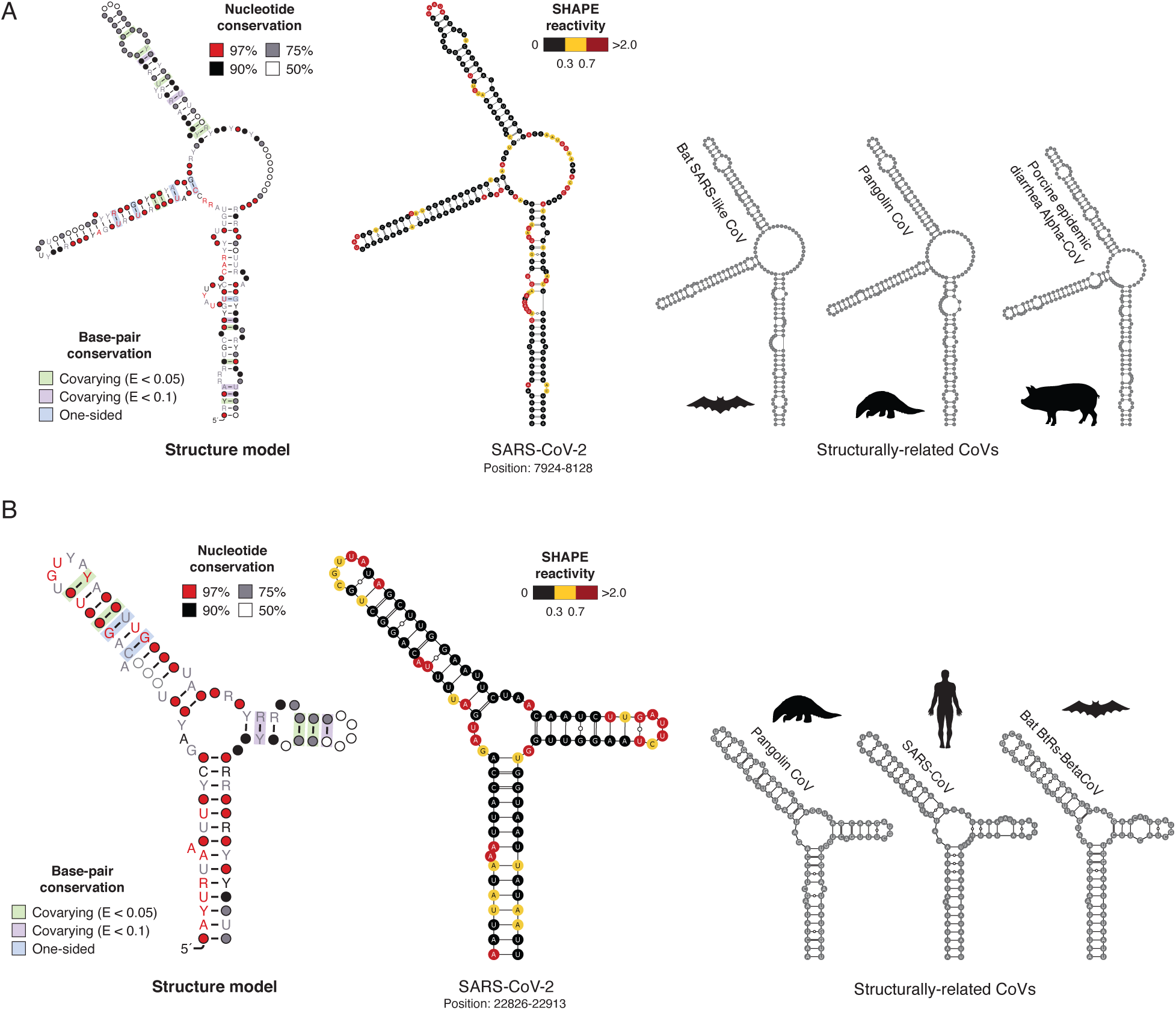
Structure of conserved elements in SARS-CoV-2 RNA. Structure models for segments 7924-8128 (**A**), and 22826-22913 (**B**). Both segments consist of conserved three-way junction structures (see Figure 4). Structure models have been generated using the R2R software. Base-pairs showing significant covariation (as determined by R-scape) are boxed in green (E-value < 0.05) and violet (E-value < 0.1) respectively. Alongside, the structure of each segment in SARS-CoV-2 with superimposed SHAPE-MaP reactivities is shown, together with the putative conserved structure in other CoV as derived from the alignment and manually optimized with the ViennaRNA package.

In summary, in this study, we described the RNA structural landscape of the SARS-CoV-2 virus. Starting with the experimentally-informed inference of the secondary structure of the SARS-CoV-2 genome, we produced 3D models for sequence segments likely to fold into stable 3D structures and we identified putative druggable pockets in these structures. Many of them presented high-information-content geometries, such as multiway junctions, that represent attractive RNA 3D targets for further analyses. Importantly, some of these structures also showed significant covariation, supporting their functional importance for SARS-CoV-2. Overall, our analyses reveal structures of the SARS-CoV-2 virus RNA that may turn out to be its weak spots. Not only our data will provide a fundamental resource for the development of innovative RNA-targeted therapeutic strategies, but also it will help elucidating still unknown aspects of the life cycle of coronaviruses once the functional role of the structured elements in the coronavirus RNA genome here identified will be characterized.

## STAR★Methods

### Resource Availability

#### Lead Contacts

Further information and requests for resources and reagents should be directed to Danny Incarnato, Janusz M. Bujnicki, or Martijn J. van Hemert.

#### Materials Availability

This study did not generate new unique reagents.

#### Data and Code Availability

SHAPE-MaP data has been deposited to the Gene Expression Omnibus (GEO) database, under the accession GSE151327. RNA Count (RC) files for use with RNA Framework, as well as XML files with normalized SHAPE-MaP reactivities are provided, together with the full secondary structure in dot-bracket notation and the selected low Shannon - low SHAPE structures. Stockholm alignment files and the derived covariance models (CMs) for the structure elements, showing significant covariation support, are also provided. For each structured segment in the SARS-CoV-2 genome, we further provide the representative 3D structure models for up to 10 of the largest clusters, illustrating the most typical conformational classes, as well as the 1000 best-energy models used to predict ligand-binding modes. In all models, the B-factor field presents the relative ligand-binding score obtained from the Fpocket analysis. These additional processed files are available at http://www.incarnatolab.com/datasets/SARS_Manfredonia_2020.php (RNA Framework RC and XML files, dot-bracket structures, CMs, Stockholm alignments) and http://dx.doi.org/10.17632/8gj97c4kgv.1 (3D models of SARS-CoV-2 segments and druggable pockets). All the scripts and tools used for data analysis are detailed in the Key Resources Table.

## Method Details

### Synthesis of NAI

NAI was synthesized as previously described (Spitale et al., 2013). Briefly, 137.14 mg of 2-methylpyridine-3-carboxylic acid (Sigma Aldrich, cat. 325228) were resuspended in 500 μl DMSO anhydrous (Sigma Aldrich, cat. 276855). ∼162.15 mg of 1,1’-Carbonyldiimidazole (Sigma Aldrich, cat. 115533) were resuspended in 500 μl DMSO anhydrous and added dropwise to the 2-methylpyridine-3-carboxylic acid solution while constantly stirring, over a period of 5 minutes. The reaction mixture was then incubated at room temperature with constant stirring for 2 hours. This mixture (assumed to represent a 1 M stock) was aliquoted in 50 μl aliquots and stored at -80°C.

### Cell culture and SARS-CoV-2 infection

Vero E6 cells were cultured in T-175 flasks in Dulbecco’s modified Eagle’s medium (DMEM; Lonza, cat. 12-604F), supplemented with 8% fetal calf serum (FCS; Bodinco), 2 mM L-glutamine, 100 U/mL of penicillin and 100 µg/mL of streptomycin (Sigma Aldrich, cat. P4333-20ML) at 37°C in an atmosphere of 5% CO2 and 95%–99% humidity. Cells were infected at a MOI of 1.5 with SARS-CoV-2/Leiden-0002 (GenBank accession: MT510999), a clinical isolate obtained from a nasopharyngeal sample at LUMC, which was passaged twice in Vero E6 cells before use. Infections were performed in Eagle’s minimal essential medium (EMEM; Lonza, cat. 12-611F) supplemented with 25 mM HEPES, 2% FCS, 2 mM L-glutamine, and antibiotics. At 16 h post-infection, infected cells were harvested by trypsinization, followed by resuspension in EMEM supplemented with 2% FCS, and then washed with 50 mL 1X PBS.

All experiments with infectious SARS-CoV-2 were performed in a biosafety level 3 facility at the LUMC.

### Total RNA extraction and *in vitro* folding

Approximately 5×10^6^ of the harvested infected cells were resuspended in 1 mL of TriPure Isolation Reagent (Sigma Aldrich, cat. 11667157001) and 200 μl of chloroform were added. The sample was vigorously vortexed for 15 sec and then incubated for 2 min at room temperature, after which it was centrifuged for 15 min at 12,500 x g (4°C). The upper aqueous phase was collected in a clean 2 mL tube, supplemented with 1 mL (∼2 volumes) of 100% ethanol, and then loaded on an RNA Clean & Concentrator-25 column (Zymo Research, cat. R1017). For *in vitro* folding, ∼2 μg of total RNA in a volume of 39 μl were denatured at 95°C for 2 min, then transferred immediately to ice and incubated for 1 min. 10 μl of ice-cold 5X RNA Folding Buffer [500 mM HEPES pH 7.9; 500 mM NaCl] supplemented with 20 U of SUPERase•In™ RNase Inhibitor (ThermoFisher Scientific, cat. AM2696) were added. RNA was then incubated for 10 min at 37°C to allow secondary structure formation. Subsequently, 1 μl of 500 mM MgCl_2_ (pre-warmed at 37°C) was added and RNA was further incubated for 20 min at 37°C to allow tertiary structure formation.

### Probing of RNA

For probing of RNA, NAI was added to a final concentration of 50 mM and samples were incubated at 37°C for 10 min. A negative control reaction was also prepared, by adding an equal amount of DMSO. Reactions were then quenched by the addition of 1 volume DTT 1 M and then purified on an RNA Clean & Concentrator-5 column (Zymo Research, cat. R1013).

### Extraction of native deproteinized *E. coli* rRNAs

Deproteinized *E. coli* RNA was prepared essentially as previously described (Simon et al., 2019), with minor changes. Briefly, a single colony of *Escherichia coli* K-12 DH10B was picked and inoculated in LB medium without antibiotics, then grown to exponential phase (OD_600_ ∼ 0.5). 2 mL aliquots were collected by centrifugation at 1000 x g (4°C) for 5 min. Cell pellets were resuspended in 1 mL of Resuspension Buffer [15 mM Tris-HCl pH 8.0; 450 mM Sucrose; 8 mM EDTA], and lysozyme was added to a final concentration of 100 μg/mL. After incubation at 22°C for 5 min and on ice for 10 min, protoplasts were collected by centrifugation at 5000 x g (4°C) for 5 min. Pellets were resuspended in 120 μl Protoplast Lysis Buffer [50 mM HEPES pH 8.0; 200 mM NaCl; 5 mM MgCl_2_; 1.5% SDS], supplemented with 0.2 μg/μl Proteinase K. Samples were incubated for 5 min at 22°C and for 5 min on ice. SDS was removed by addition of 30 μl SDS Precipitation Buffer [50 mM HEPES pH 8.0; 1 M Potassium Acetate; 5 mM MgCl_2_], followed by centrifugation at 17,000 x g (4°C) for 5 min. Supernatant was then extracted 2 times with phenol:chloroform:isoamyl alcohol (25:24:1, pre-equilibrated 3 times with a buffer containing [50 mM HEPES pH 8.0; 200 mM NaCl; 5 mM MgCl_2_]), and once with chloroform. 20 U SUPERase•In™ RNase Inhibitor were then added and RNA was equilibrated at 37°C for 20 min prior to probing.

### *Ex vivo* probing of *E. coli* rRNAs

180 μl of deproteinized *E. coli* rRNAs, pre-equilibrated at 37°C for 20 min, were mixed with 20 μl of NAI. RNA was then allowed to react at 37°C for 15 minutes, with moderate shaking, after which 200 μl of 1 M DTT were added, to quench the reaction. Samples were then vortexed briefly and 1 mL of ice-cold QIAzol added to each sample, followed by extensive vortexing. RNA extraction was carried out as already described above for SARS-CoV-2-infected cells.

### Multiplex SHAPE-MaP of SARS-CoV-2 RNA

For multiplex SHAPE-MaP, 70 oligonucleotide pairs, tiling the entire length of the SARS-CoV-2 genome (29,903 nt), were automatically designed using Primer3 (Untergasser et al., 2012) and the following parameters: amplicon size between 480 and 520 bp, maximum poly(N) length between 2 and 3, minimum/optimal/maximum oligonucleotide size of 20/25/30, minimum/optimal/maximum Tm of 56/60/62 degrees, minimum/optimal/maximum GC content of 30/50/65 %. Also, pairs were designed in such a way that the target amplicon would include the target region of the reverse primer from the previous set and of the forward primer of the following set. Primers were then searched against the GENCODE v33 human transcriptome, keeping only those with less than 60% predicted base-pairing or more than 60% predicted base-pairing and more than 2 mismatched bases at the 3′ end. For reverse transcription, all reverse primers were equimolarly pooled to yield a 100 μM RT primer mix. An additional anchored oligo-dT primer (TTTTTTTTTTTTTTTTTTTTVN) was added to the mix. For each sample, two 20 μl reverse transcription reactions were carried out. 3 μg of total RNA were mixed with 0.5 μl RT primer mix and 1 μl dNTPs (10 mM each), incubated at 70°C for 5 min and then transferred immediately to ice for 1 min. Reactions were then supplemented with 4 μl 5X RT Buffer [250 mM Tris-HCl pH 8.3; 375 mM KCl], 2 μl DTT 0.1 M, 1 μl MnCl_2_ 120 mM, 20 U SUPERase•In™ RNase Inhibitor and 200 U SuperScript II RT (ThermoFisher Scientific, cat. 18064014). Reactions were incubated at 42°C for 2 h and then RT was heat-inactivated at 75°C for 15 min. RNA was then degraded by the addition of 5 U RNase H (New England Biolabs, cat. M0297S) and cDNA from every 2 reactions was purified on a single RNA Clean & Concentrator-5 column and eluted in 36 μl NF H_2_O.

For targeted amplification, primer pairs were split into 2 non-overlapping sets, odd and even. For each of these sets, pairs were pooled in smaller sets of 3-4 pairs, as follows: 1-19-37-55, 2-20-38-56, 3-21-39-57, 4-22-40-58, 5-23-41-59, 6-24-42-60, 7-25-43-61, 8-26-44-62, 9-27-45-63, 10-28-46-64, 11-29-47-65, 12-30-48-66, 13-31-49-67, 14-32-50-68, 15-33-51-69, 16-34-52-70, 17-35-53, 18-36-54. The complete primer list can be found in Table S1. PCR reactions were carried out in 50 μl, using 2 μl of eluted cDNA, 0.3 mM final each dNTP, 0.1 μM final each primer in the set and 2.5 U TaKaRa Taq™ DNA Polymerase (TaKaRa, cat. R001A). Cycling was performed using a touch-down approach. Briefly, for the first 10 cycles, the annealing temperature was lowered by 0.5°C per cycle, starting at 58°C. Then, 20 additional cycles were performed at the lowest temperature. PCR products were purified using 1 volume of NucleoMag NGS Clean-up and Size Select beads (Macherey Nagel, cat. 744970), checked on a 2% agarose gel. Each primer set gave a single specific band of the expected size. PCR products were equimolarly pooled to yield the final odd and even sets. 500 ng of each set were fragmented in a final volume of 20 μl, using 2 μl of NEBNext dsDNA Fragmentase (New England Biolabs, cat. M0348), supplemented with 1 μl 200 mM MgCl_2_, by incubating at 37°C for 25 min. This yielded fragments in the range of ∼50-200 bp. Reactions were purified using 1 volume of NucleoMag NGS Clean-up and Size Select beads. 5 ng of fragmented DNA were then used as input for the NEBNext® Ultra™ II DNA Library Prep Kit for Illumina® (New England Biolabs, cat. E7645L), as per manufacturer instructions.

### SHAPE-MaP of *E. coli* rRNAs

Total *E. coli* RNA was first fragmented to a median size of 150 nt by incubation at 94°C for 8 min in RNA Fragmentation Buffer [65 mM Tris-HCl pH 8.0; 95 mM KCl; 4 mM MgCl_2_], then purified with NucleoMag NGS Clean-up and Size Select beads (Macherey Nagel, cat. 744970), supplemented with 10 U SUPERase•In™ RNase Inhibitor, and eluted in 2 μl NF H_2_O. Eluted RNA was supplemented with 0.5 μl 20 μM random hexamers and 0.25 μl dNTPs (10 mM each), then incubated at 70°C for 5 min and immediately transferred to ice for 1 min. Reverse transcription reactions were conducted in a final volume of 5 μl. Reactions were supplemented with 1 μl 5X RT Buffer [250 mM Tris-HCl pH 8.3; 375 mM KCl], 0.5 μl DTT 0.1 M, 0.25 μl MnCl_2_ 120 mM, 5 U SUPERase•In™ RNase Inhibitor and 50 U SuperScript II RT. Reactions were incubated at 25°C for 10 min to allow partial primer extension, followed by 2 h at 42°C. SSII was heat-inactivated by incubating at 75°C for 15 min. 6 mM final EDTA was added to chelate Mn^2+^ ions, followed by 5 min incubation at room temperature and addition of 6 mM final MgCl_2_. Reverse transcription reactions were then used as input for the NEBNext® Ultra II Non-Directional RNA Second Strand Synthesis Module (New England Biolabs, cat. E6111L). Second strand synthesis was performed by incubating 1 h at 16°C, as per manufacturer instructions. DsDNA was purified using NucleoMag NGS Clean-up and Size Select beads, and used as input for the NEBNext® Ultra™ II DNA Library Prep Kit for Illumina, following manufacturer instructions.

### SHAPE-MaP data analysis

Analysis of SHAPE-MaP data has been conducted using RNA Framework v2.6.9 (https://github.com/dincarnato/RNAFramework; Incarnato et al., 2018). Reads pre-processing and mapping was performed using the *rf-map* module (parameters: *-ctn -cmn 0 -cqo -cq5 20 -b2 -mp “--very-sensitive-local”*). Reads were trimmed of terminal Ns and low-quality bases (Phred < 20). Reads with internal Ns were discarded. Mapping was performed using the “very-sensitive-local” preset of Bowtie2 (Langmead and Salzberg, 2012). The mutational signal was then derived using the *rf-count* module (parameters: *-m -rd*), enabling the right re-alignment of deletions. When processing SARS-CoV-2 data, a mask file containing the sequences of primer pairing regions was passed along (through the *-mf* parameter). Generated RC files, from both the even and odd sets, containing per base mutation counts and coverage, were then combined in a single RC file using the *rf-rctools* module (mode: *merge*). Data normalization was performed using the *rf-norm* module (parameters: *-sm 3 -nm 3 -n 1000 -mm 1*), by setting the minimum base coverage to 1,000X and the maximum mutation frequency to 1, and by using the Siegfried *et al*., 2014 scoring scheme (Siegfried et al., 2014) and box-plot normalization of SHAPE reactivities.

### SHAPE-informed SARS-CoV-2 RNA secondary structure modeling

To model the secondary structure of the SARS-CoV-2 genome, we first sought to determine optimal slope/intercept parameters by grid search, using the *rf-jackknife* module (parameters: *-rp “-md 600 -nlp” -x*) and ViennaRNA package v2.4.14 (Lorenz et al., 2011), with *E. coli* SHAPE-MaP data and reference *E. coli* 16S/23S rRNA structures (Cannone et al., 2002). Isolated base-pairs were disallowed, the maximum base-pairing distance was set to 600 nt and the relaxed structure comparison mode was enabled (a base pair *i/j* was considered to be correctly predicted if any of the following pairs exist in the reference structure: *i/j*; *i-1/j*; *i+1/j*; *i/j-1*; *i/j+1*; Deigan et al., 2009). Optimal slope and intercept parameters were respectively determined to be -0.8 and 0.2 (extremely close to previously determined parameters -1.1 and 0; Huber et al., 2019).

As folding the full SARS-CoV-2 genome as a single entity would be an extremely challenging, and currently unfeasible, computational task, folding was performed using the *rf-fold* module and a windowed approach previously used by us and other groups (Siegfried et al., 2014; Simon et al., 2019) to model viral RNA genomes (parameters: *-sl 0.8 -in -0.2 -w -fw 3000 -fo 300 -wt 200 -pw 1500 -po 250 -dp -sh -nlp -md 600*). At all stages, SHAPE data was incorporated in the form of soft constraints. Briefly, in the first stage partition function folding was performed by sliding a window of 1,500 nt along the genome, with an offset of 250 nt. For each window, the first and last 100 nt were ignored, to avoid terminal biases. Two additional foldings were computed at both the 5′ and 3′ ends to increase structure sampling (window sizes: 1,400 and 1,450 nt). Base-pairing probabilities from all windows were then averaged and base-pairs with a probability ≥0.99 were kept and used as a constraint for the next stage. At this stage, base-pairing probabilities were also used to calculate per-base Shannon entropies. In the second stage, the maximum expected accuracy (MEA) structure was predicted by sliding a window of 3,000 nt, with an offset of 300 nt along the genome. Four additional foldings were computed at both the 5′ and 3′ ends to increase structure sampling (window sizes: 2,900, 2,950, 3,050, and 3,100 nt). Base-pairs appearing in >50% of the windows were then retained to yield the final secondary structure pseudoknot-free model. Pseudoknots were instead introduced during the modeling of the 3D structure (see paragraph “Selection of structured segments for RNA 3D structure prediction” below).

### Identification of candidate low Shannon – low/high SHAPE regions

For the identification of low Shannon – low SHAPE regions, median Shannon entropies and SHAPE reactivities were first calculated in sliding, centered 50 nt windows. Then, a window of 50 nt was slid along the genome. Windows in which >75% of the bases were below both the global Shannon and SHAPE median (calculated on the full SARS-CoV-2 genome) were picked and windows residing less than 10 nt apart were merged. Only structure elements having >50% of the base-pairs falling in a low Shannon – low SHAPE region were kept for downstream analyses. For the identification of low Shannon – high SHAPE regions, instead, a window of 25 nt was slid along the genome. Windows in which >75% of the bases were below the global Shannon median and above the global SHAPE median (calculated on the full SARS-CoV-2 genome) and >50% of the bases were predicted to be single-stranded in the MEA structure, were picked and windows residing less than 10 nt apart were merged.

### Identification of conserved RNA structure elements

To identify conserved low Shannon – low SHAPE RNA structure elements, we implemented an automated pipeline (*cm-builder*; https://github.com/dincarnato/labtools) built on top of Infernal 1.1.3 (Nawrocki and Eddy, 2013). Briefly, we first built a covariance model (CM) from a Stockholm file containing only the SARS-CoV-2 sequence and the structure of the selected elements, using the *cmbuild* module. After calibrating the CM using the *cmcalibrate* module, it was used to search for RNA homologs in a database composed of all the non-redundant coronavirus complete genome sequences from the ViPR database (https://www.viprbrc.org/brc/home.spg?decorator=corona; Pickett et al., 2011), as well as a set of representative coronavirus genomes from NCBI database, using the *cmsearch* module. Only matches from the sense strand were kept and a very relaxed E-value threshold of 10 was used at this stage to select potential homologs. Three additional filtering criteria were used. First, we took advantage of the extremely conserved architecture of coronavirus genomes (Lauber et al., 2013) and restricted the selection to matches falling at the same relative position within their genome, with a tolerance of 3.5% (roughly corresponding to a maximum allowed shift of 1050 nt in a 30 kb genome). Through this more “conservative” selection, we only kept matches likely to represent true structural homologs, although at the cost of probably losing some true matches. Second, we filtered out matches retaining less than 55% of the canonical base-pairs from the original structure elements. Third, truncated hits covering <50% of the structure were discarded. The resulting set of homologs was then aligned to the original CM using the *cmalign* module and the resulting alignment was used to build a new CM. The whole process was repeated for a total of 3 times. The alignment was then refactored, removing gap-only positions and including only bases spanning the first to the last base-paired residue. The alignment file was then analyzed using R-scape 1.4.0 (Rivas et al., 2016) and APC-corrected G-test statistics to identify motifs showing significantly covarying base-pairs. To increase the likelihood of selecting only bona fide conserved RNA structural elements, we discarded any structure with less than 5% covarying pairs at E-value < 0.05 and less than 10% covarying pairs at E-value < 0.1. Stockholm files for these structures were then used to build the final CMs and to re-search the entire database with the default stringent E-value cutoff of 0.01.

### Determination of low Shannon – high SHAPE regions’ conservation

To assess the sequence conservation of the identified low Shannon – high SHAPE regions, we computed 4 multiple sequence alignments using MAFFT v7.429 (parameters: *--maxiterate 100 --auto*; Katoh and Standley, 2013), the reference SARS-CoV-2 sequence and one of the following datasets: 1) SARS-CoV (243 sequences); 2) MERS-CoV (281 sequences); 3) other Beta-CoV (excluding SARS-CoV/SARS-CoV-2/MERS-CoV, 681 sequences); 4) other CoV (excluding Beta-CoV, 1657 sequences). Sequences were obtained from the ViPR database and 100% identical sequences were collapsed. Region conservation was calculated as the average of the conservation of each nucleotide in that region (using SARS-CoV-2 as the reference).

### Selection of structured segments for RNA 3D structure prediction

Starting from the chosen low Shannon - low SHAPE regions, sequence segments for RNA 3D structure prediction were selected based on their predicted structural complexity, mutual proximity, as well as the proximity of other predicted structural elements. Briefly, neighboring segments were combined with each other. Very short (< 40 nt) low Shannon - low SHAPE regions predicted to fold into simple hairpins were not analyzed further, unless they could be combined with each other or incorporated into neighboring larger fragments. When predicted secondary structure elements were present in the immediate proximity of low Shannon – low SHAPE regions, suggesting the possibility of additional structural interactions, the sequence segments were expanded to include them. For example, the modeled segment for the frameshifting element comprises a short low Shannon - low SHAPE 70 nt region that was extended in both the 5′ and 3′ directions in order to encompass the neighboring regions predicted to be structured yielding a 180 nt-long segment.

For the final set of 70 segments, we carried out an additional secondary structure prediction step with a customized version of RNAProbe (Wirecki et al., 2020), using ShapeKnots (incorporating SHAPE-MaP reactivities) to evaluate the presence of potential pseudoknots (Hajdin et al., 2013). Finally, the base-pairing patterns common to both the original pseudoknot-free MEA prediction and the ShapeKnots prediction were used to restrain the 3D folding simulations.

### RNA 3D structure modeling

3D structures were predicted for the aforementioned segments using SimRNA, a method for simulations of RNA folding, which uses a coarse-grained representation, relies on the Monte Carlo method for sampling the conformational space, and employs a statistical potential to approximate the energy of the studied RNA molecule (Boniecki et al., 2015; Magnus et al., 2016). The previously derived consensus secondary structure was used as a restraint. Briefly, for segments >90 nt, which exhibited complex structures, we carried out eight independent simulations, each comprising 10 replicas, with 16 million Monte Carlo steps in each run). For segments <90 nt, that were mostly expected to adopt hairpin-like structures, we carried out only one simulation (10 replicas, 16 million Monte Carlo steps). Each simulation resulted in a trajectory, comprising up to 10,000 conformations, broadly representing local energy minima in the whole conformational space sampled by SimRNA for a given molecule.

For each segment, the 5% top-scored conformations were clustered at the RMSD threshold equal to 0.1 times the length of the sequence, yielding clusters of similar conformations (up to 10 clusters for each sequence segment). A single representative of each cluster was retained to represent the most probable class of conformations. Further, 1000 best-scored conformations for each segment, which together represent the low-energy regions of the conformational landscape and approximate the distribution of most common conformations existing in solution, were used to identify potential binding sites for small molecules. For each of the 1000 models, the secondary structure was extracted by running ClaRNA (Waleń et al., 2014) and the set of 1000 secondary structures was used to generate the consensus structure for each segment.

### Prediction of potential ligand binding sites in RNA 3D structures (druggable pockets)

To identify potential ligand-binding pockets in RNA 3D structures, we used Fpocket (Le Guilloux, Schmidtke, and Tuffery 2009), originally developed for the identification of ligand-binding pockets in proteins. We conducted a benchmarking study on a small subset of known viral RNA structures bound to their small molecule ligands (PDB codes: 1LVJ [HIV-1 TAR with PMZ], 1UTS [HIV-1 TAR with P13], 2KMJ [HIV-2 TAR with a pyrimidinylpeptide], 2KTZ [HCV IRES with ISH], and 2L94 [HIV-1 FSE with the frameshifting inhibitor DB213]). These RNAs were folded using SimRNA (one simulation with 10 replicas and 16 million Monte Carlo steps). For each RNA, 1000 decoys with the lowest SimRNA energy were identified and analyzed with Fpocket. In each analyzed model, a ribonucleotide residue with at least two atoms within the pocket was considered as a potential ligand-contacting residue. For each residue in each RNA, a score of 0 to 100 was assigned based on the total number of contacts predicted in the 1000 low-energy conformations. The residues with a score of 40 or above successfully recapitulated the regions in the structures to which ligands were binding, as experimentally determined in 4 out of 5 structures. The sole instance in which a binding pocket could not be retrieved by Fpocket was the ligand-binding site without a specific pocket in the 2L94 structure, in which the ligand binds in the major groove of the RNA helix.

To retrieve the potentially druggable pockets in complex 3D motifs from the SARS-CoV-2 models, we applied a more stringent criterion. The residues with an Fpocket score of 40 or above were selected and extended by one residue on either side to include the neighbors at the sequence level. After the extension, the stretches with six nucleotides or more were retained. The shorter stretches were added, if any of their constituent residues were paired with the longer ones, based on the consensus secondary structure derived from 1000 lowest-energy decoys. Additional residues forming Watson-Crick base pairs with the nucleotides in these stretches were also included in further analyses. For each of the 1000 decoys, pairwise atom-atom distances were calculated to identify groups of residues present as neighbors in 3D-space. The residues consistently present within a 12 Å cut-off distance were assigned to one pocket. The identified pockets were clustered and representatives (most prevalent conformation) were selected for up to the five biggest clusters.

## Acknowledgements

D.I. was supported by the Dutch Research Council (NWO) as part of the research programme NWO Open Competitie ENW - XS with project number OCENW.XS3.044 and by the Groningen Biomolecular Sciences and Biotechnology Institute (GBB, University of Groningen). I.M. and T.M. were supported by personal funding from the GBB and University of Groningen to D.I.. J.M.B. and C.N. were supported by the National Science Centre, Poland (grant nr 2017/26/A/NZ1/01083 to J.M.B.). P.G. was supported by the Foundation for Polish Science (grant nr TEAM/2016-3/18 to J.M.B.). T.K.W. was supported by the National Science Centre, Poland (grant nr 2017/25/B/NZ2/01294 to J.M.B.). A.P.S. was supported by the National Science Centre, Poland (grant nr. 2018/31/D/NZ2/01883 to A.P.S.). Computational resources for calculations done by the Bujnicki group were provided by the Poznań Supercomputing and Networking Center at the Institute of Bioorganic Chemistry, Polish Academy of Sciences through the Polish Grid Infrastructure (grants: rnpmd, simcryox, and plgnithinaneesh2019a) and the Interdisciplinary Centre for Mathematical and Computational Modelling at the University of Warsaw (grants: G73-4, GB76-30, and G81-5). N.O. was supported by the Horizon 2020 research and innovation program under the Marie Skłodowska-Curie grant agreement nr 642434 (ITN Antivirals). Part of this research was supported by the Leiden University Fund (LUF), the Bontius Foundation and donations from the crowdfunding initiative “wake up to corona”.

We would like to further acknowledge Dr. Diana Spierings, Dr. Nancy Halsema, and all the staff at the sequencing facility of ERIBA (European Research Institute for the Biology of Ageing, University Medical Center Groningen).

## Author Contributions

Project conceptualization: D.I.; SARS-CoV-2 isolation and characterization: N.S.O. and E.J.S.; Wet-lab: I.M., T.M., and M.J.H.; Bioinformatics, structure modeling and data analysis: I.M., C.N., T.K.W., P.G., A.P.S., J.M.B., and D.I.; Writing: all the authors.

