## Supplemental Information for "Genome-wide mapping of therapeutically-relevant SARS-CoV-2 RNA structures"

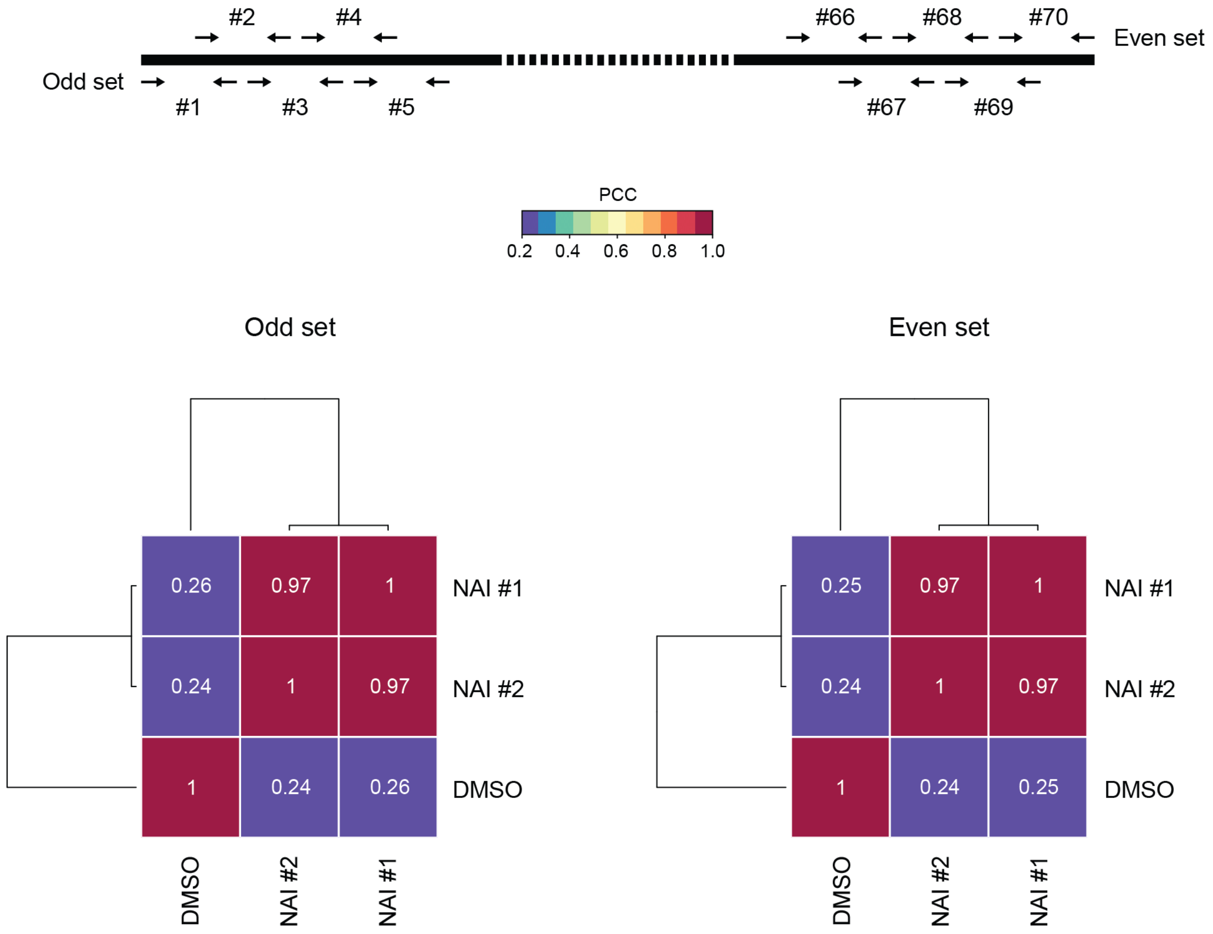


**Figure S1.** Correlation calculated on SHAPE-MaP raw reactivities (mutation frequencies) across the odd and even sets.

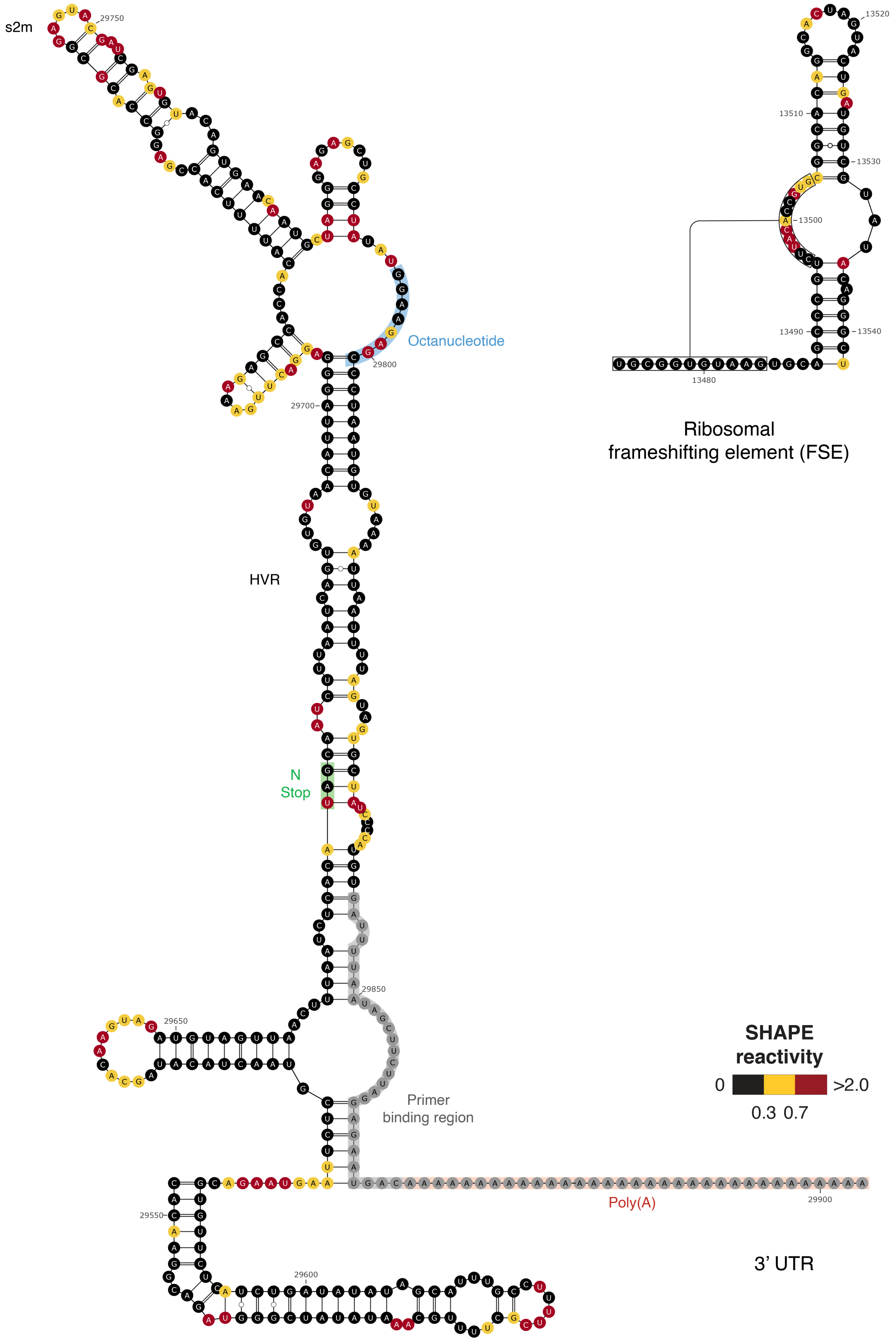


**Figure S2.** Measured SHAPE-MaP reactivities superimposed on the reference Sarbecovirus 3’ UTR and ribosomal frameshifting element (FSE) structures.

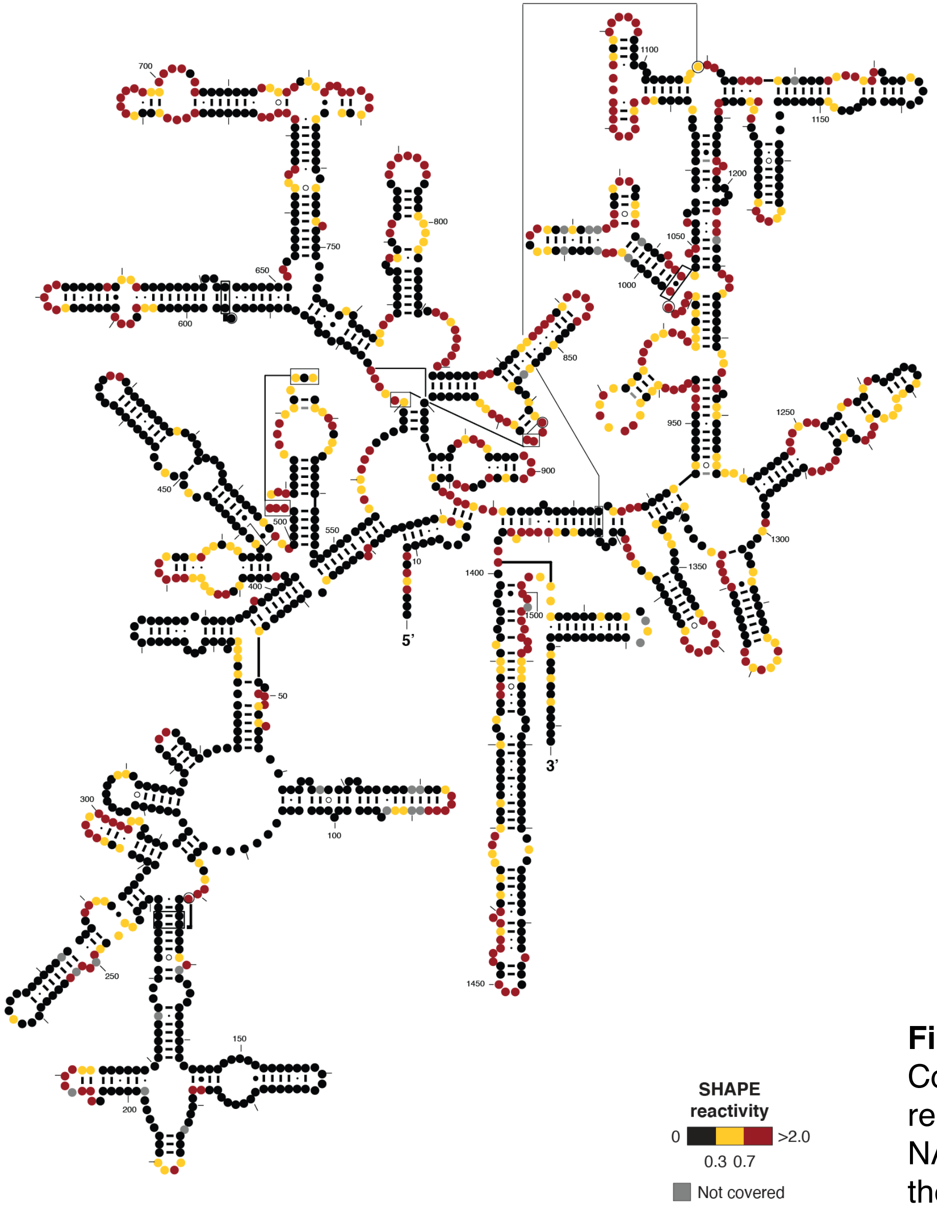


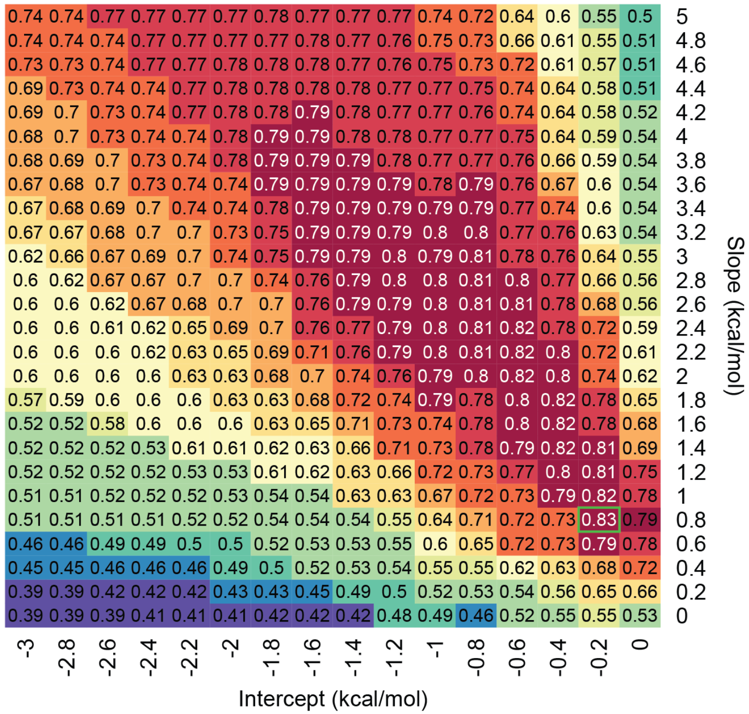


**Figure S3.** (*left*) Phylogenetic structure of *E. coli* 16S rRNA (from the Comparative RNA Web, CRW) with superimposed SHAPE-MaP reactivities. (*right*) Grid search of the optimal slope/intercept pair for NAI. Values represent the geometric mean of sensitivity and PPV for the *E. coli* 16S/23S rRNA secondary structures predicted using each slope/intercept value pair. The chosen value pair is boxed in green.

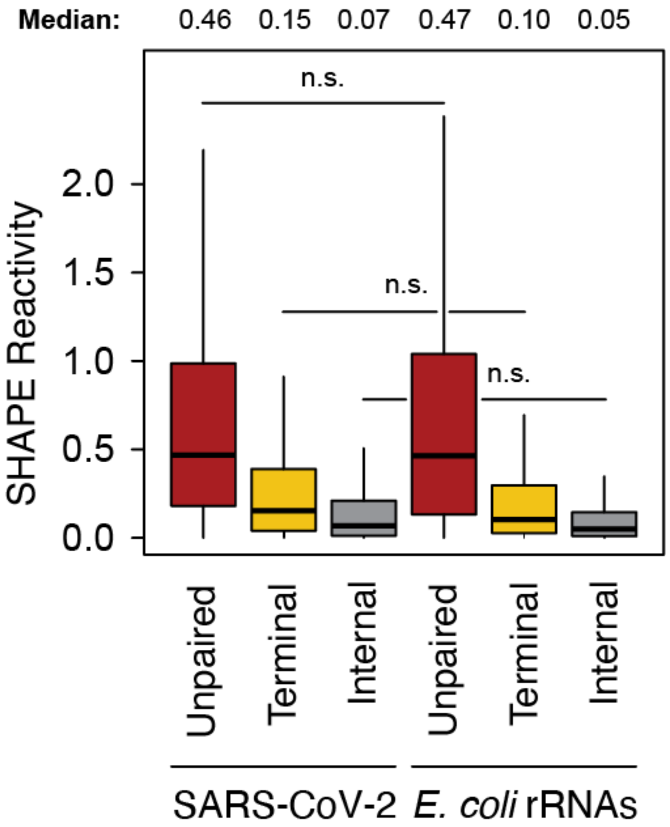


**Figure S4.** Box-plot illustrating the distribution of SHAPE reactivities across unpaired, terminally-paired, and internally-paired residue for both the proposed SARS-CoV-2 RNA model and the reference *E. coli* 16S/23S rRNA structures. No significant difference is observed between the distributions of corresponding classes across the 2 RNAs. All the other comparisons are highly significant (One-way ANOVA, $P < 2.2e\text{-}16$).

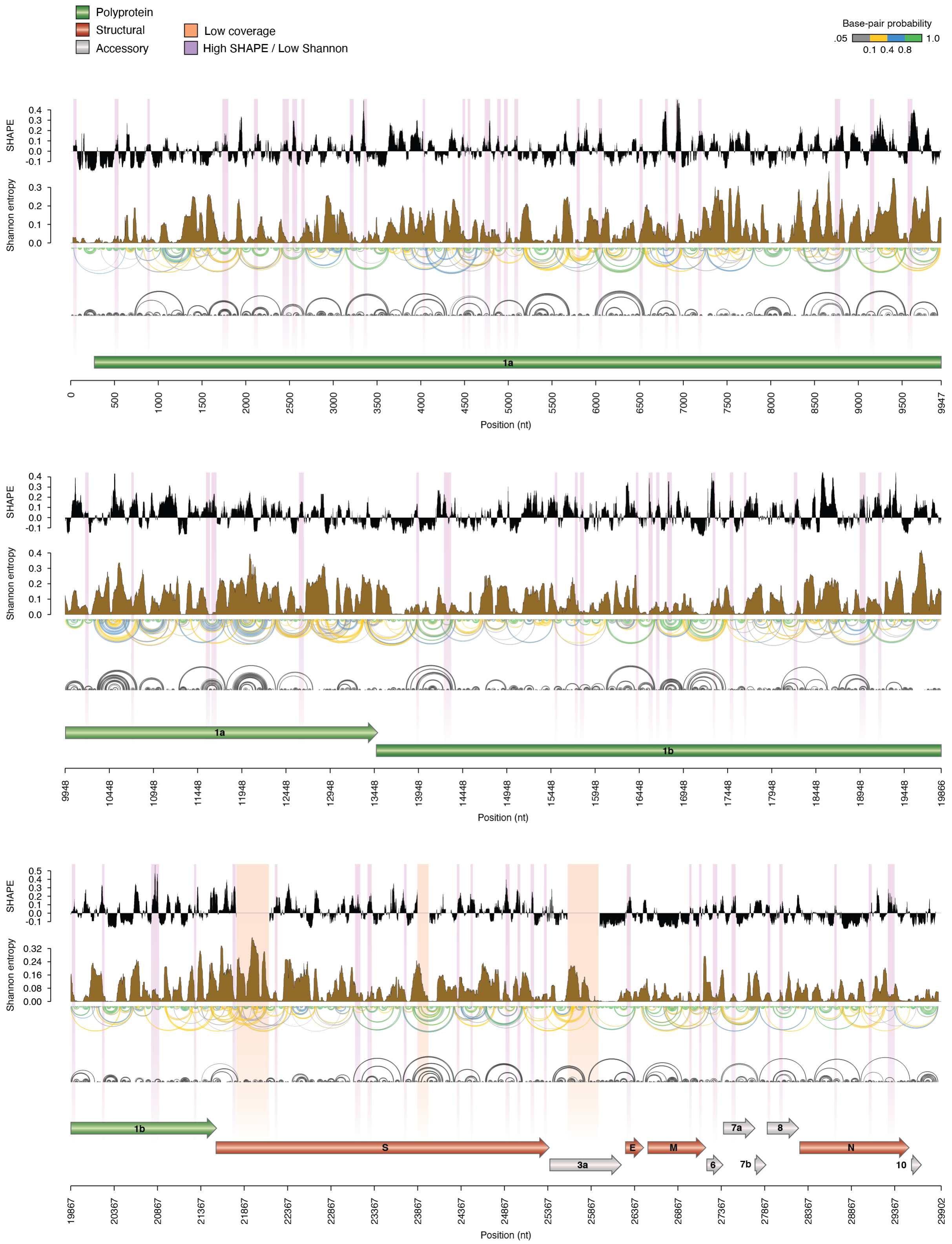


**Figure S5.** Map of the SARS-CoV-2 genome depicting (top to bottom): median SHAPE reactivity (in 50 nt centered widows, with respect to the median reactivity across the whole genome), Shannon entropy, base-pairing probabilities, maximum expected accuracy (MEA) structure. Regions with both low Shannon and high SHAPE, likely identifying RNA regions of persistent single-strandedness, are marked in violet.

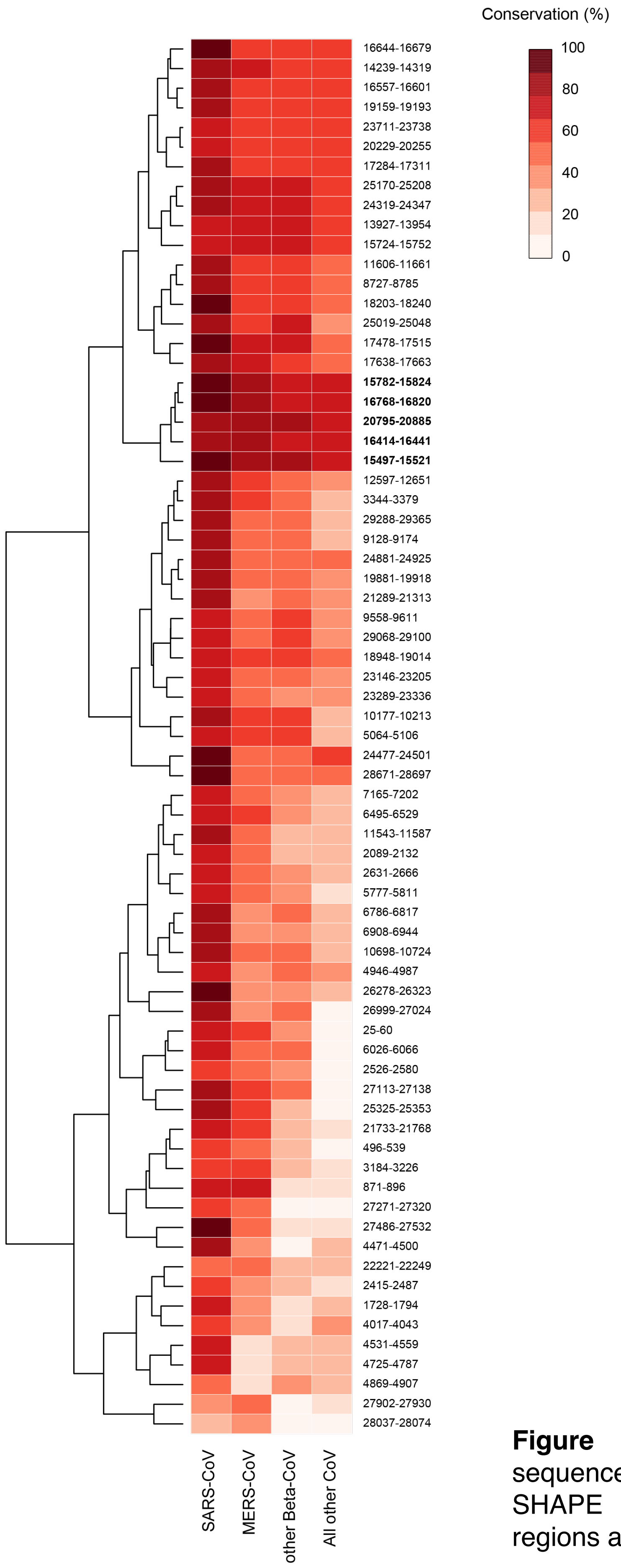


**Figure S6.** Heatmap illustrating the % sequence conservation of low Shannon – high SHAPE regions. The five top-conserved regions are marked in bold.

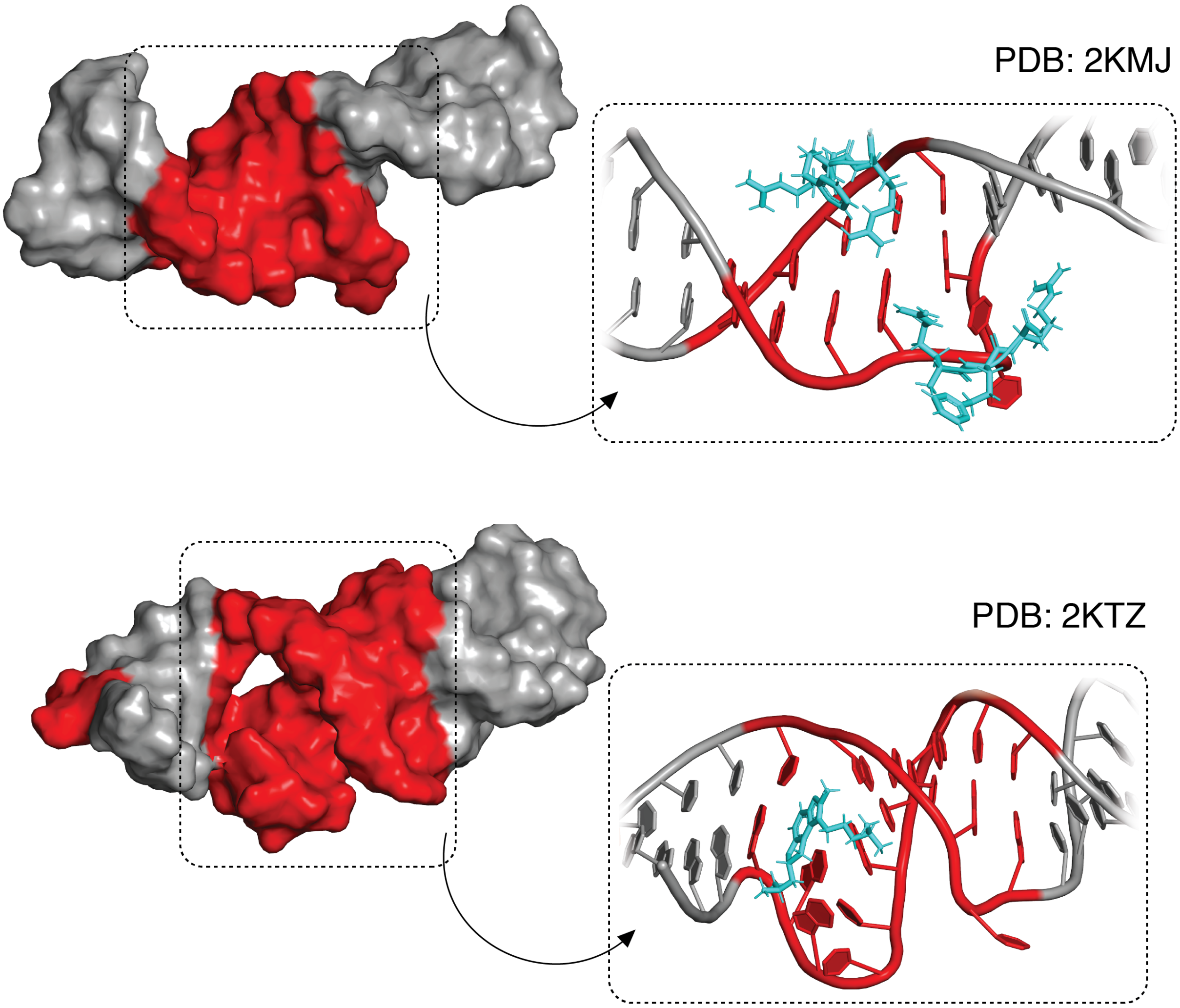


**Figure S7.** Example of two experimentally determined 3D structures of small viral RNAs complexed with small molecule ligands (HIV-2 TAR [2KMJ] and HCV IRES Domain IIa [2KTZ]). The druggable pockets identified by Fpocket are indicated in red.

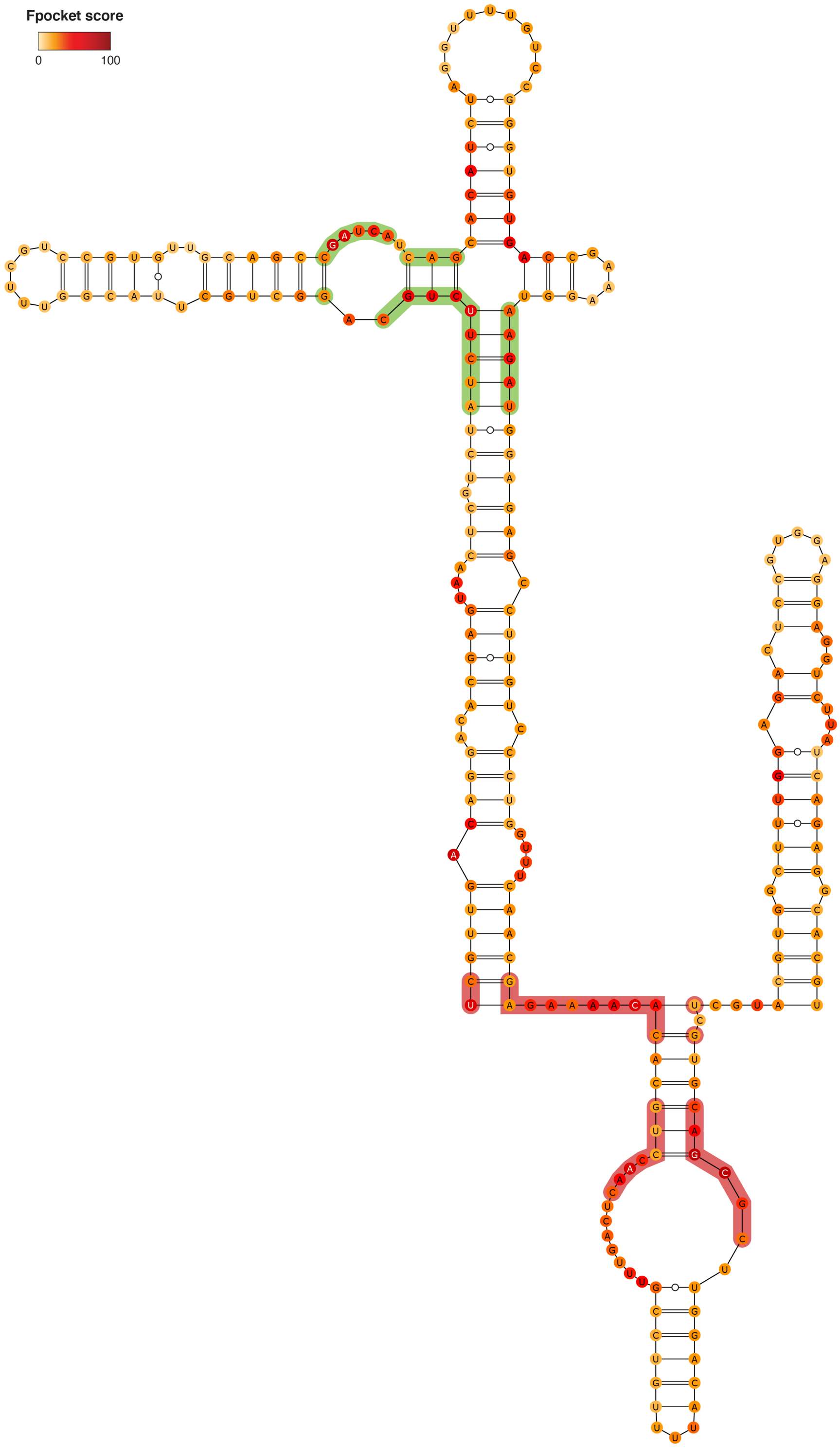


**Figure S8.** Consensus structure for segment 150-394, as derived from the 1000 lowest energy 3D structure structures modeled by SimRNA (using the MEA secondary structure inferred from SHAPE-MaP as a restraint). Bases are color-coded according to the Fpocket normalized score (0-100; see Methods). Residues composing the identified druggable pockets are shaded. Different shading colors mark distinct pockets.

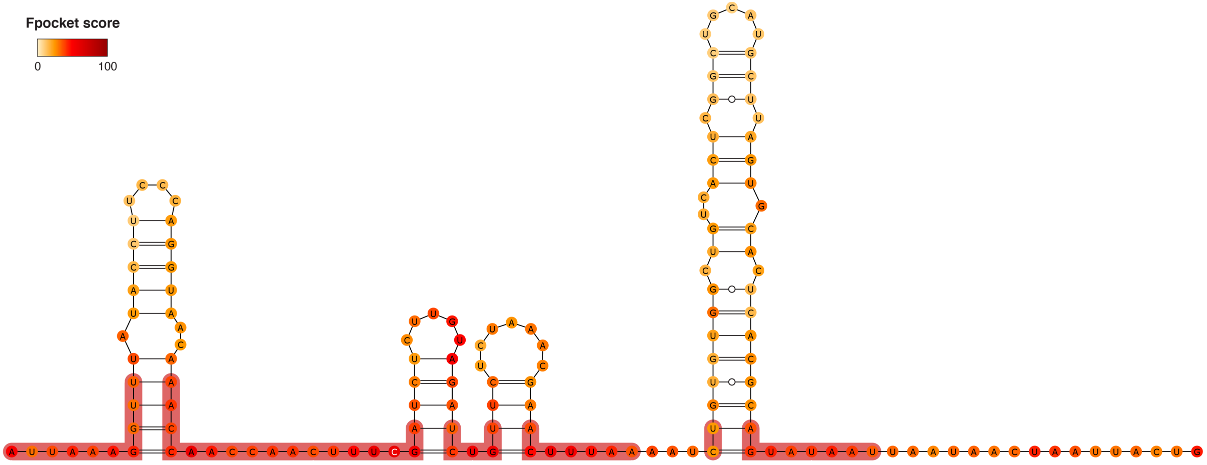


**Figure S9.** Consensus structure for segment 1-149, as derived from the 1000 lowest energy 3D structure structures modeled by SimRNA (using the MEA secondary structure inferred from SHAPE-MaP as a restraint). Bases are color-coded according to the Fpocket normalized score (0-100; see Methods). Residues composing the identified druggable pocket are shaded.

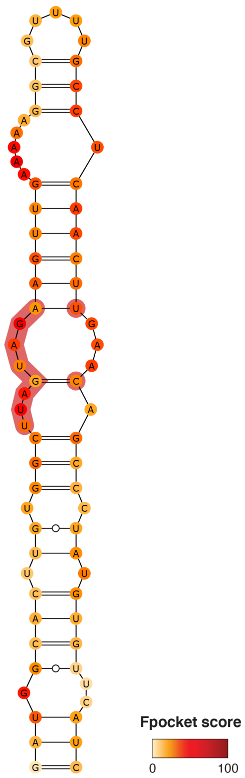


**Figure S10.** Consensus structure for segment 407-478, as derived from the 1000 lowest energy 3D structure structures modeled by SimRNA (using the MEA secondary structure inferred from SHAPE-MaP as a restraint). Bases are color-coded according to the Fpocket normalized score (0-100; see Methods). Residues composing the identified druggable pocket are shaded.

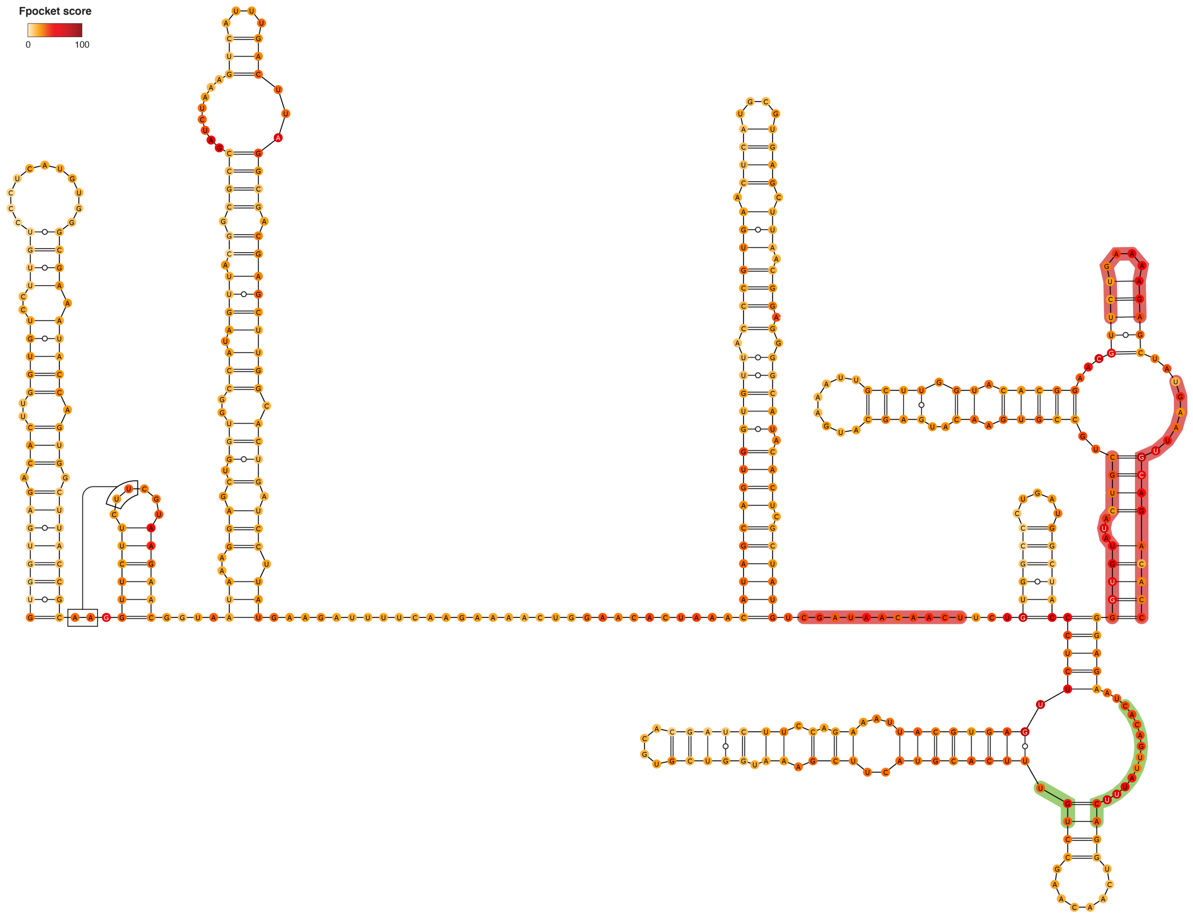


**Figure S11.** Consensus structure for segment 564-1026, as derived from the 1000 lowest energy 3D structure structures modeled by SimRNA (using the MEA secondary structure inferred from SHAPE-MaP as a restraint). Bases are color-coded according to the Fpocket normalized score (0-100; see Methods). Residues composing the identified druggable pockets are shaded. Different shading colors mark distinct pockets.

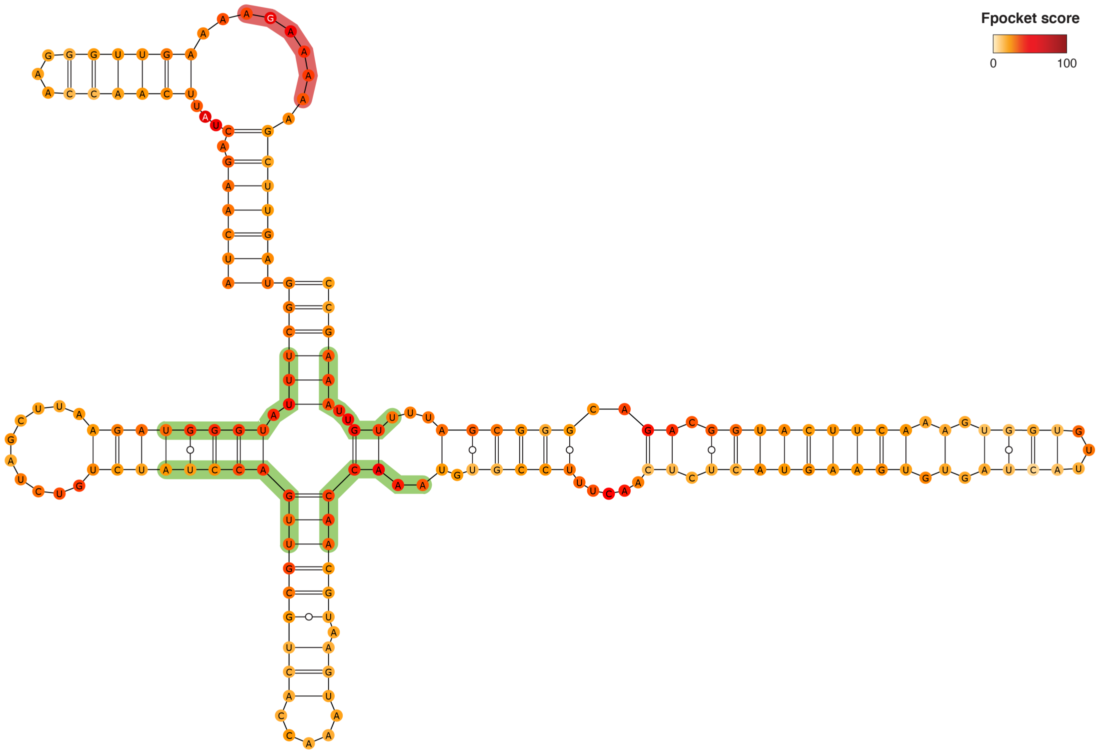


**Figure S12.** Consensus structure for segment 1106-1282, as derived from the 1000 lowest energy 3D structure structures modeled by SimRNA (using the MEA secondary structure inferred from SHAPE-MaP as a restraint). Bases are color-coded according to the Fpocket normalized score (0-100; see Methods). Residues composing the identified druggable pockets are shaded. Different shading colors mark distinct pockets.

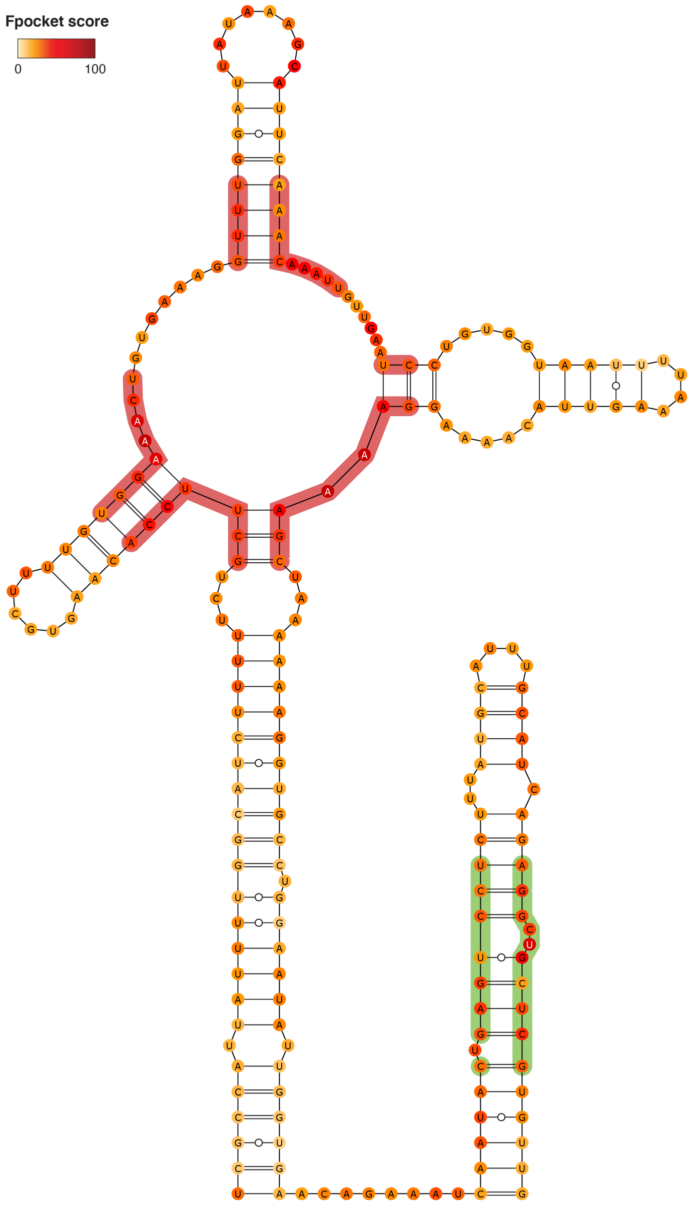


**Figure S13.** Consensus structure for segment 1683-1895, as derived from the 1000 lowest energy 3D structure structures modeled by SimRNA (using the MEA secondary structure inferred from SHAPE-MaP as a restraint). Bases are color-coded according to the Fpocket normalized score (0-100; see Methods). Residues composing the identified druggable pockets are shaded. Different shading colors mark distinct pockets.

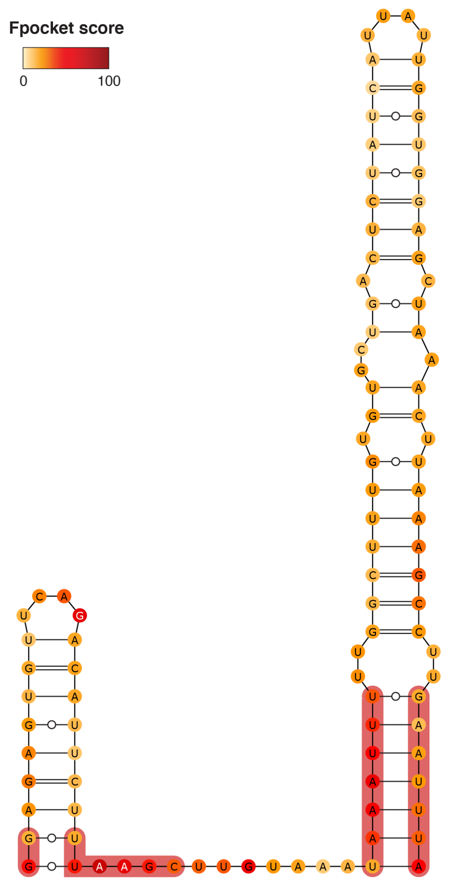


**Figure S14.** Consensus structure for segment 2281-2380, as derived from the 1000 lowest energy 3D structure structures modeled by SimRNA (using the MEA secondary structure inferred from SHAPE-MaP as a restraint). Bases are color-coded according to the Fpocket normalized score (0-100; see Methods). Residues composing the identified druggable pocket are shaded.

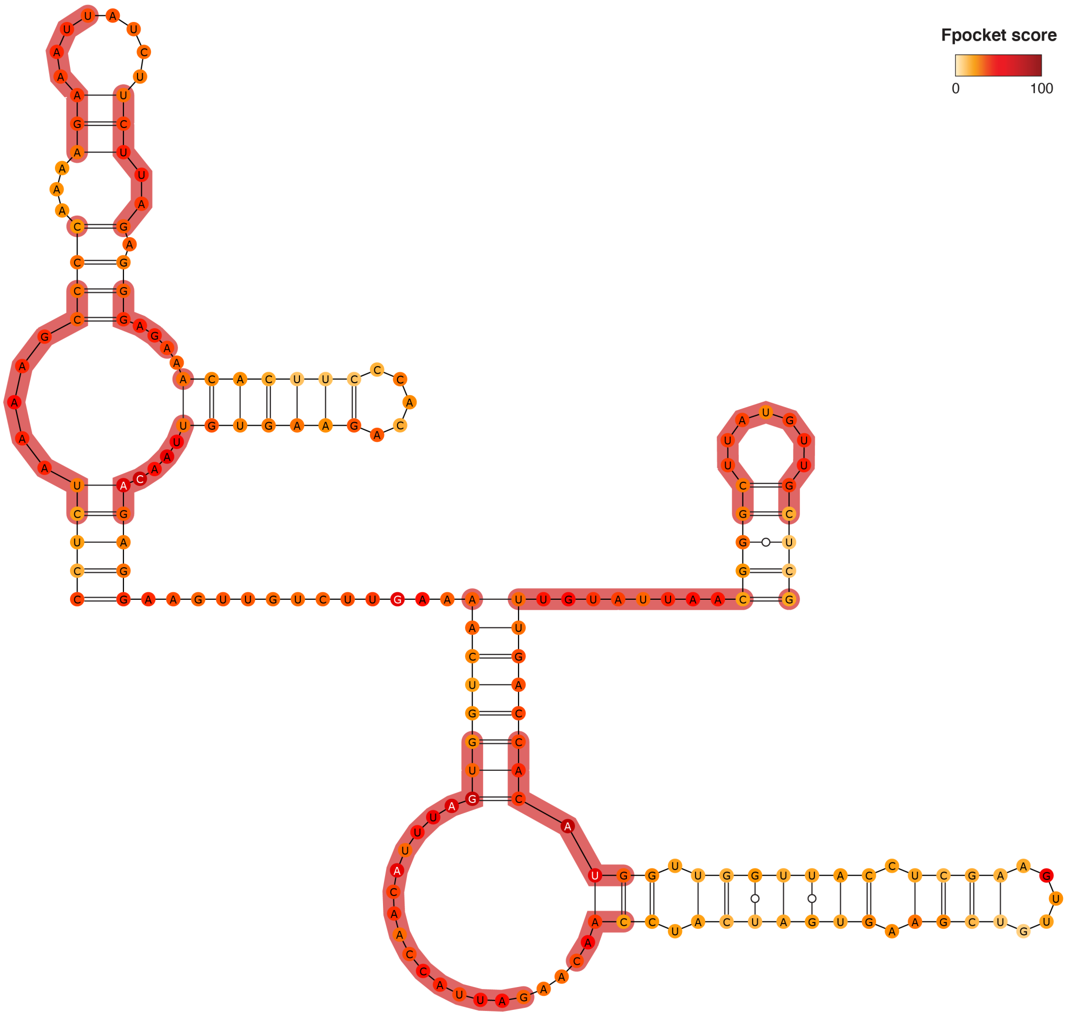


**Figure S15.** Consensus structure for segment 2459-2639, as derived from the 1000 lowest energy 3D structure structures modeled by SimRNA (using the MEA secondary structure inferred from SHAPE-MaP as a restraint). Bases are color-coded according to the Fpocket normalized score (0-100; see Methods). Residues composing the identified druggable pocket are shaded.

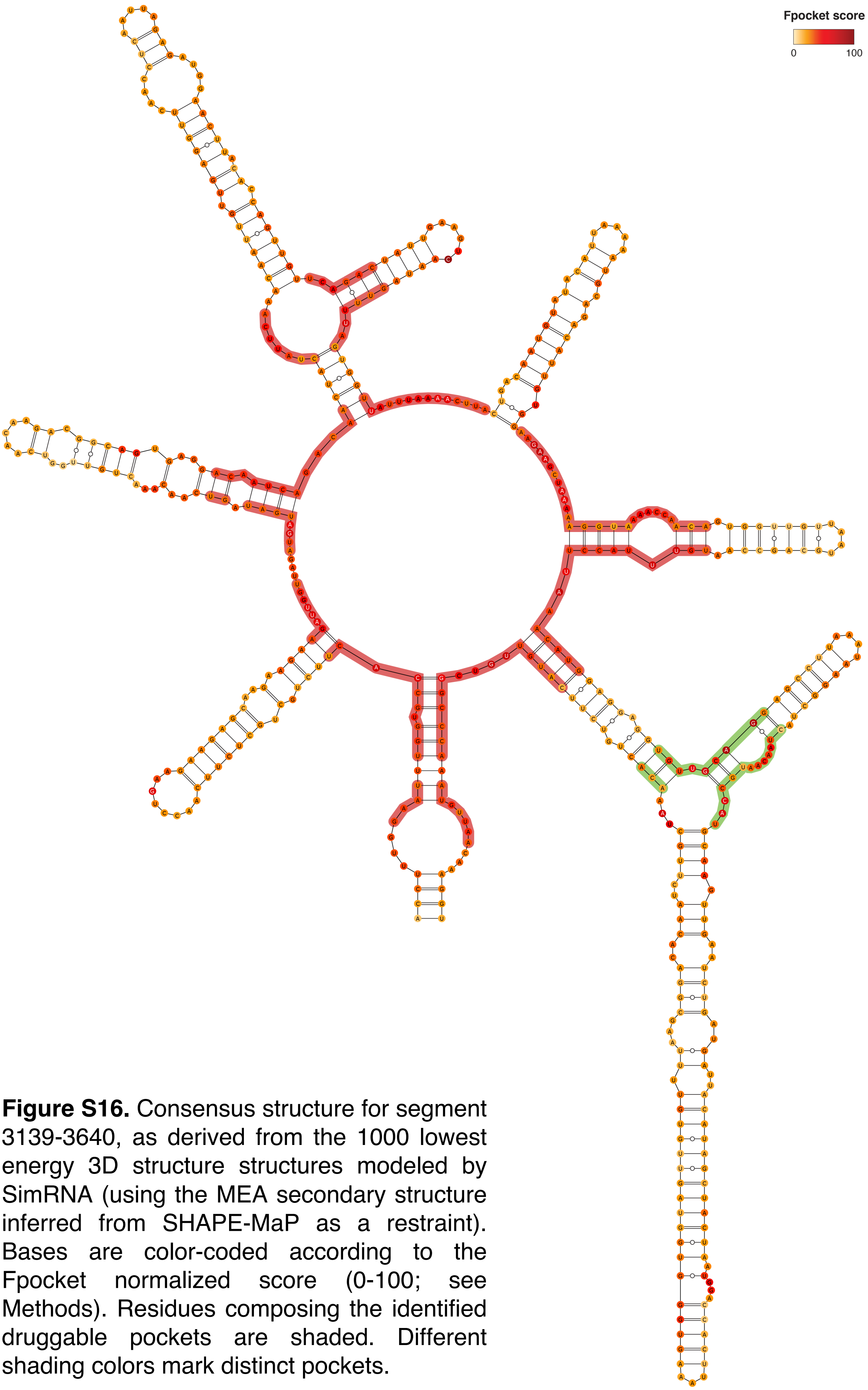


**Figure S16.** Consensus structure for segment 3139-3640, as derived from the 1000 lowest energy 3D structure structures modeled by SimRNA (using the MEA secondary structure inferred from SHAPE-MaP as a restraint). Bases are color-coded according to the Fpocket normalized score (0-100; see Methods). Residues composing the identified druggable pockets are shaded. Different shading colors mark distinct pockets.

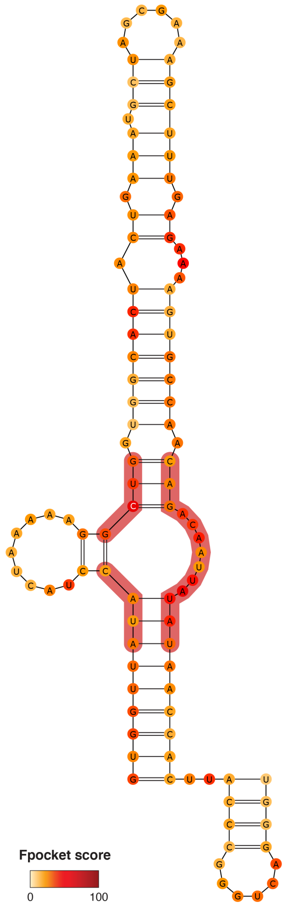


**Figure S17.** Consensus structure for segment 4160-4264, as derived from the 1000 lowest energy 3D structure structures modeled by SimRNA (using the MEA secondary structure inferred from SHAPE-MaP as a restraint). Bases are color-coded according to the Fpocket normalized score (0-100; see Methods). Residues composing the identified druggable pocket are shaded.

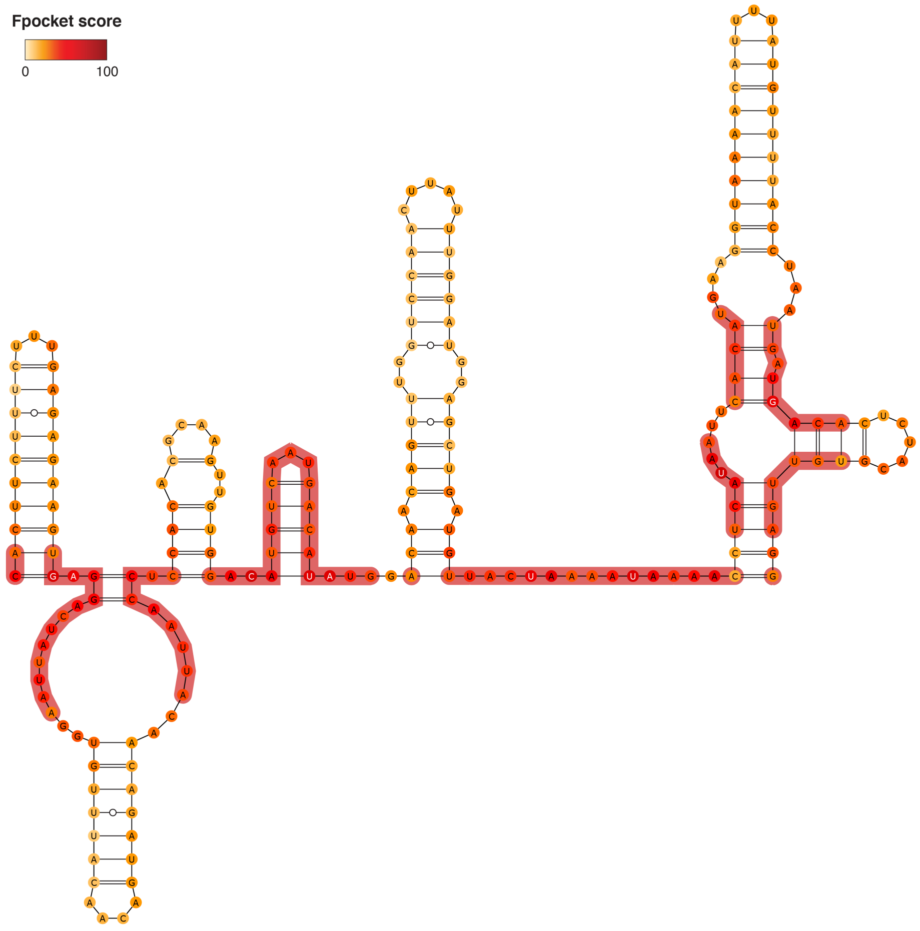


**Figure S18.** Consensus structure for segment 4938-5156, as derived from the 1000 lowest energy 3D structure structures modeled by SimRNA (using the MEA secondary structure inferred from SHAPE-MaP as a restraint). Bases are color-coded according to the Fpocket normalized score (0-100; see Methods). Residues composing the identified druggable pocket are shaded.

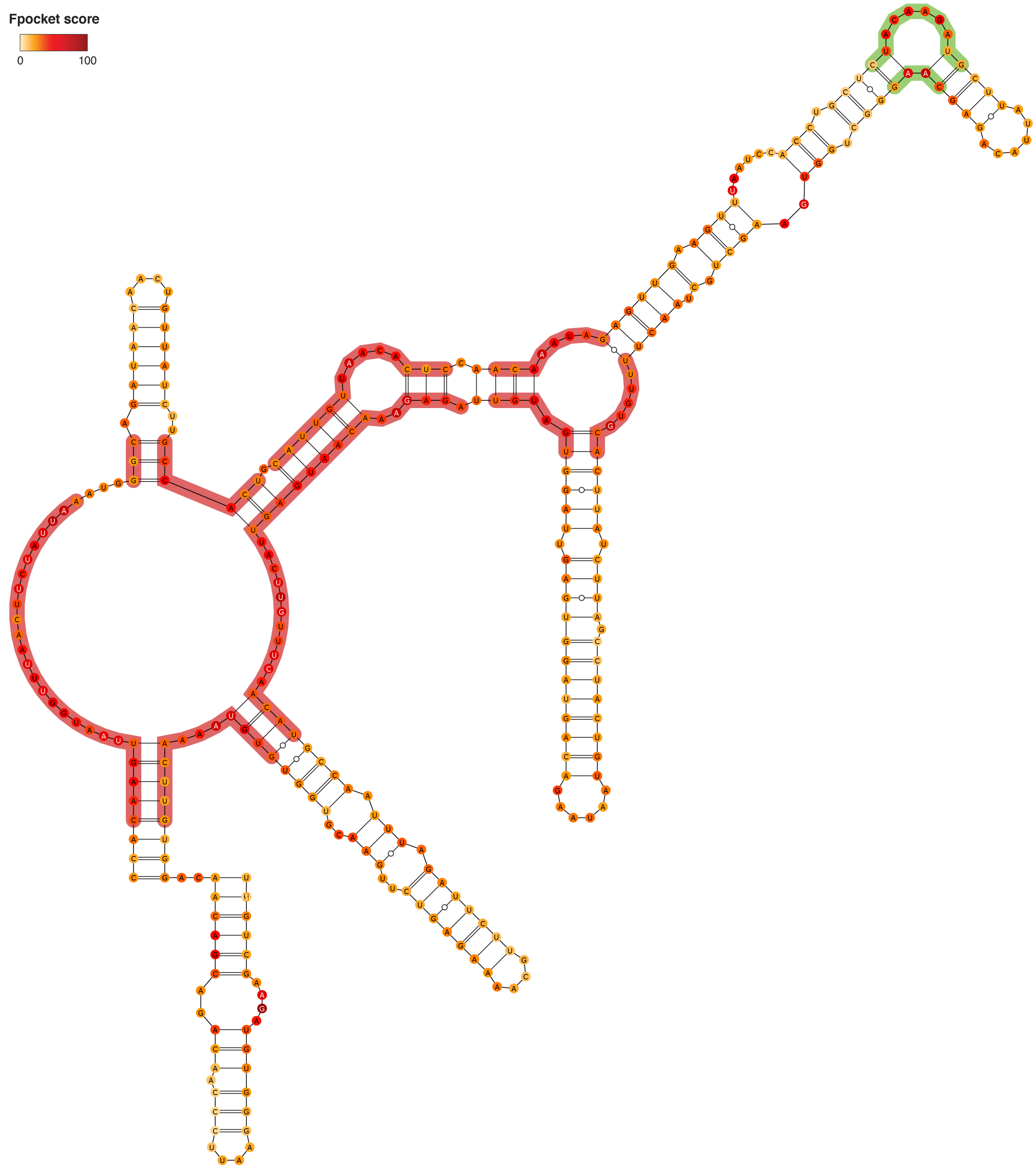


**Figure S19.** Consensus structure for segment 5240-5569, as derived from the 1000 lowest energy 3D structure structures modeled by SimRNA (using the MEA secondary structure inferred from SHAPE-MaP as a restraint). Bases are color-coded according to the Fpocket normalized score (0-100; see Methods). Residues composing the identified druggable pockets are shaded. Different shading colors mark distinct pockets.

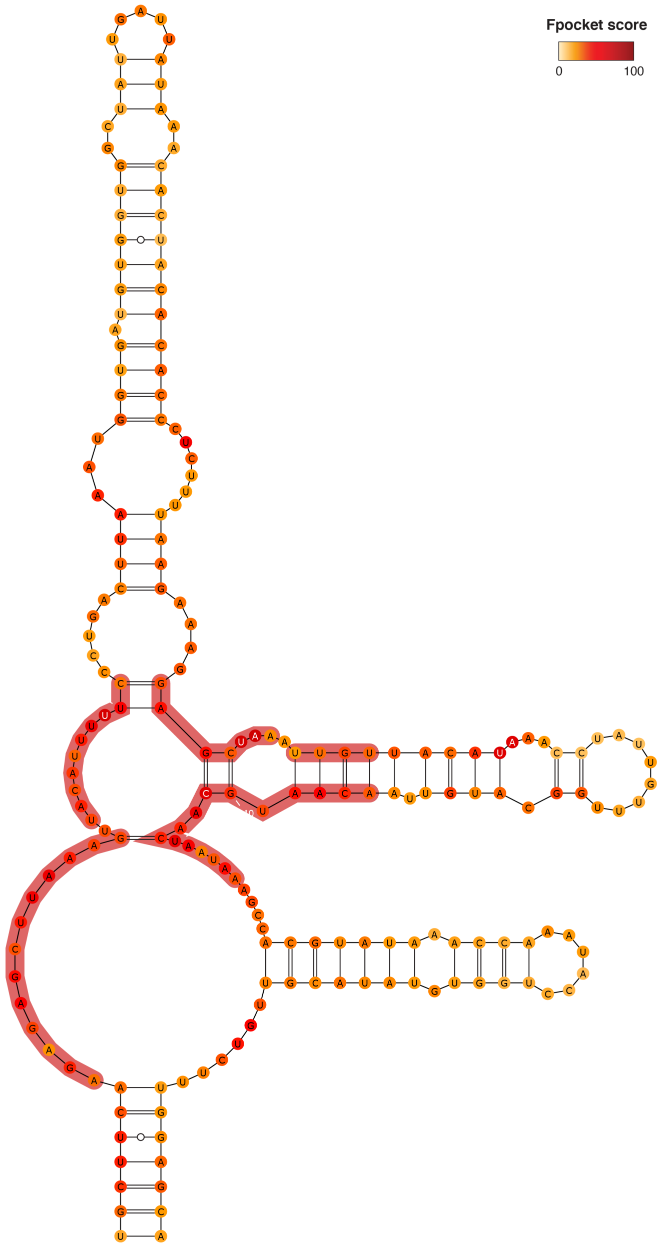


**Figure S20.** Consensus structure for segment 6115-6311, as derived from the 1000 lowest energy 3D structure structures modeled by SimRNA (using the MEA secondary structure inferred from SHAPE-MaP as a restraint). Bases are color-coded according to the Fpocket normalized score (0-100; see Methods). Residues composing the identified druggable pocket are shaded.

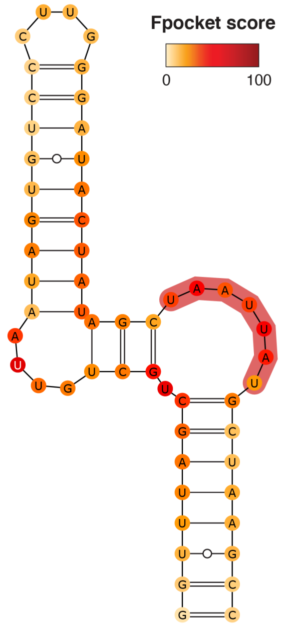


**Figure S21.** Consensus structure for segment 6641-6696, as derived from the 1000 lowest energy 3D structure structures modeled by SimRNA (using the MEA secondary structure inferred from SHAPE-MaP as a restraint). Bases are color-coded according to the Fpocket normalized score (0-100; see Methods). Residues composing the identified druggable pocket are shaded.

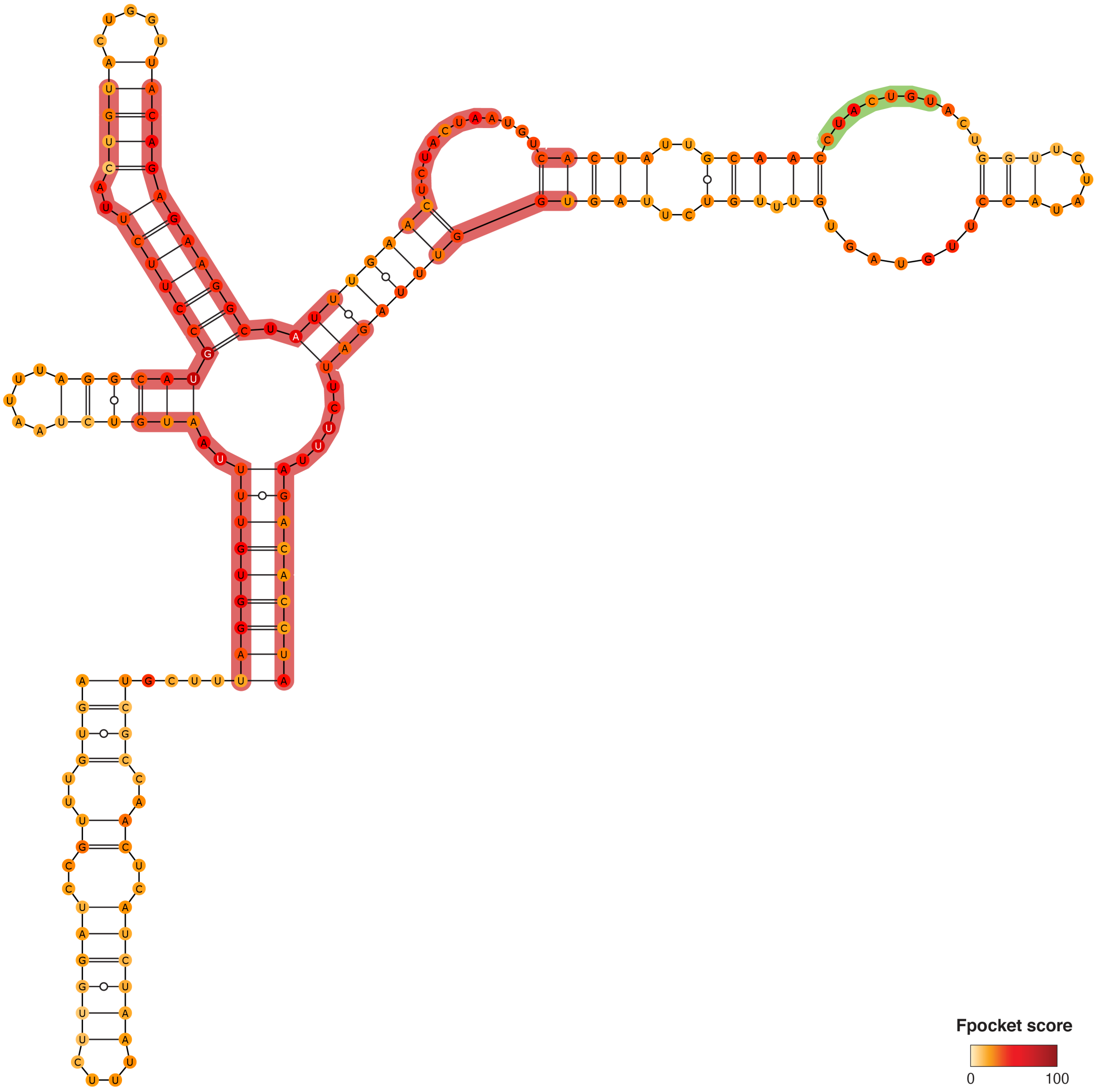


**Figure S22.** Consensus structure for segment 6974-7167, as derived from the 1000 lowest energy 3D structure structures modeled by SimRNA (using the MEA secondary structure inferred from SHAPE-MaP as a restraint). Bases are color-coded according to the Fpocket normalized score (0-100; see Methods). Residues composing the identified druggable pockets are shaded. Different shading colors mark distinct pockets.

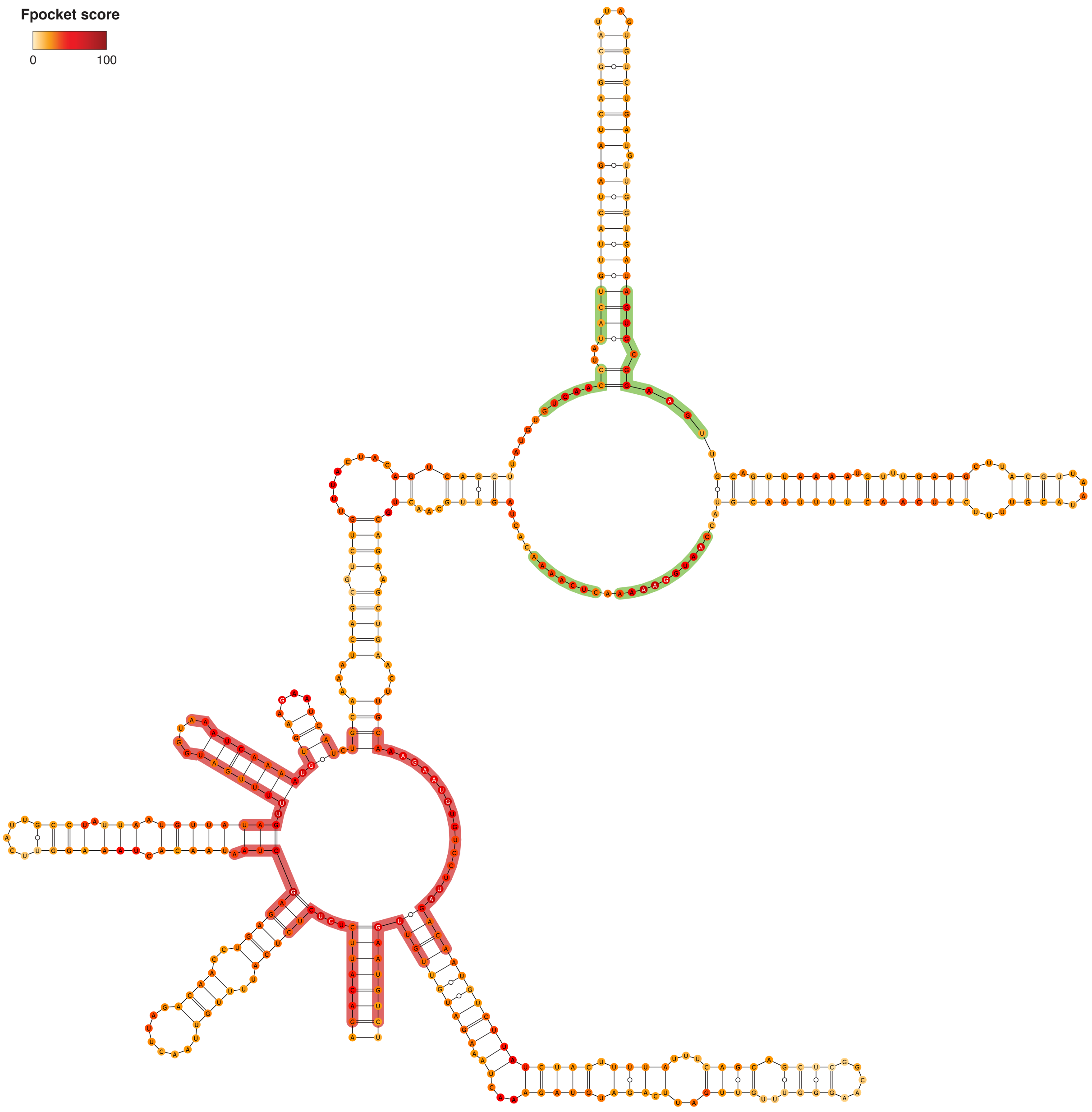


**Figure S23.** Consensus structure for segment 7808-8229, as derived from the 1000 lowest energy 3D structure structures modeled by SimRNA (using the MEA secondary structure inferred from SHAPE-MaP as a restraint). Bases are color-coded according to the Fpocket normalized score (0-100; see Methods). Residues composing the identified druggable pockets are shaded. Different shading colors mark distinct pockets.

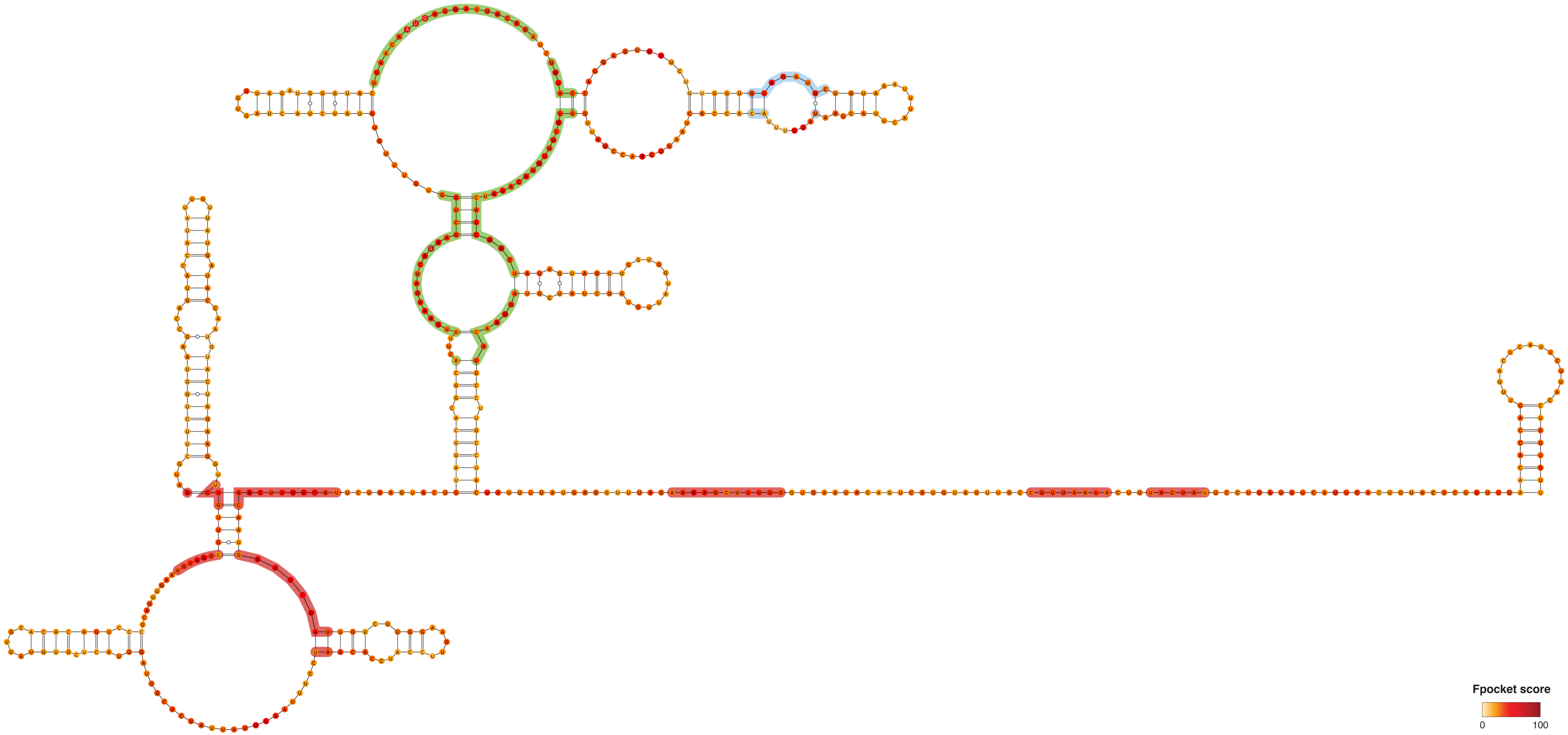


**Figure S24.** Consensus structure for segment 9035-9577, as derived from the 1000 lowest energy 3D structure structures modeled by SimRNA (using the MEA secondary structure inferred from SHAPE-MaP as a restraint). Bases are color-coded according to the Fpocket normalized score (0-100; see Methods). Residues composing the identified druggable pockets are shaded. Different shading colors mark distinct pockets.

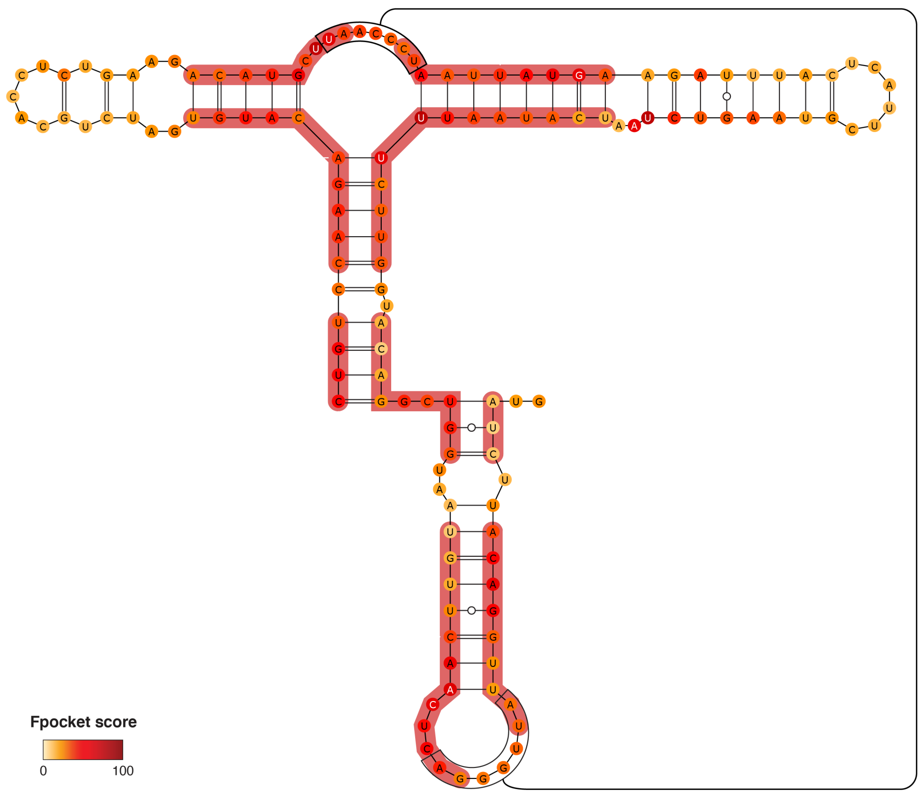


**Figure S25.** Consensus structure for segment 10165-10300, as derived from the 1000 lowest energy 3D structure structures modeled by SimRNA (using the MEA secondary structure inferred from SHAPE-MaP as a restraint). Bases are color-coded according to the Fpocket normalized score (0-100; see Methods). Residues composing the identified druggable pocket are shaded.

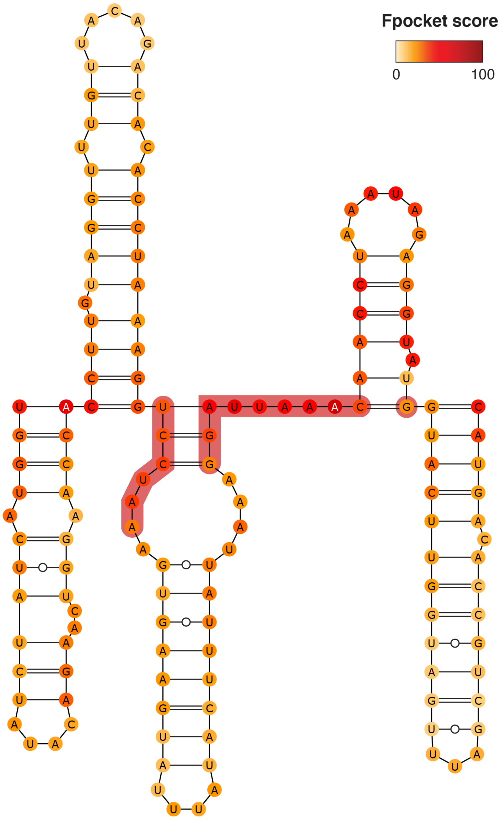


**Figure S26.** Consensus structure for segment 12871-13016, as derived from the 1000 lowest energy 3D structure structures modeled by SimRNA (using the MEA secondary structure inferred from SHAPE-MaP as a restraint). Bases are color-coded according to the Fpocket normalized score (0-100; see Methods). Residues composing the identified druggable pocket are shaded.

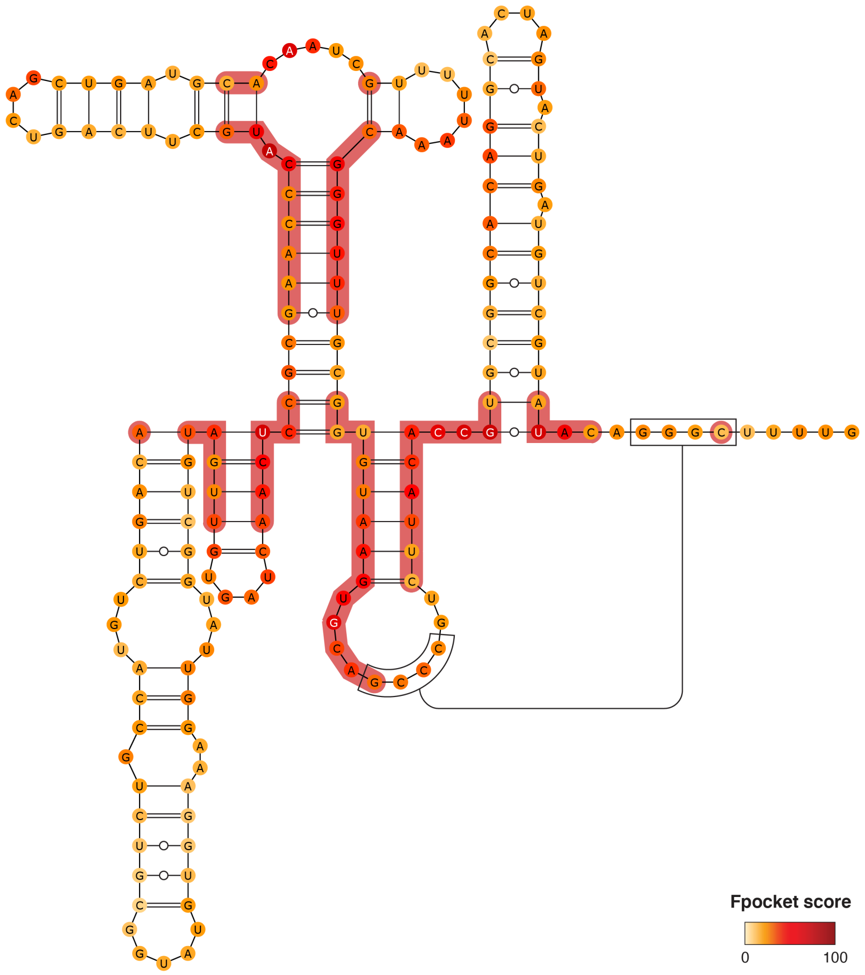


**Figure S27.** Consensus structure for segment 13367-13546, as derived from the 1000 lowest energy 3D structure structures modeled by SimRNA (using the MEA secondary structure inferred from SHAPE-MaP as a restraint). Bases are color-coded according to the Fpocket normalized score (0-100; see Methods). Residues composing the identified druggable pocket are shaded.

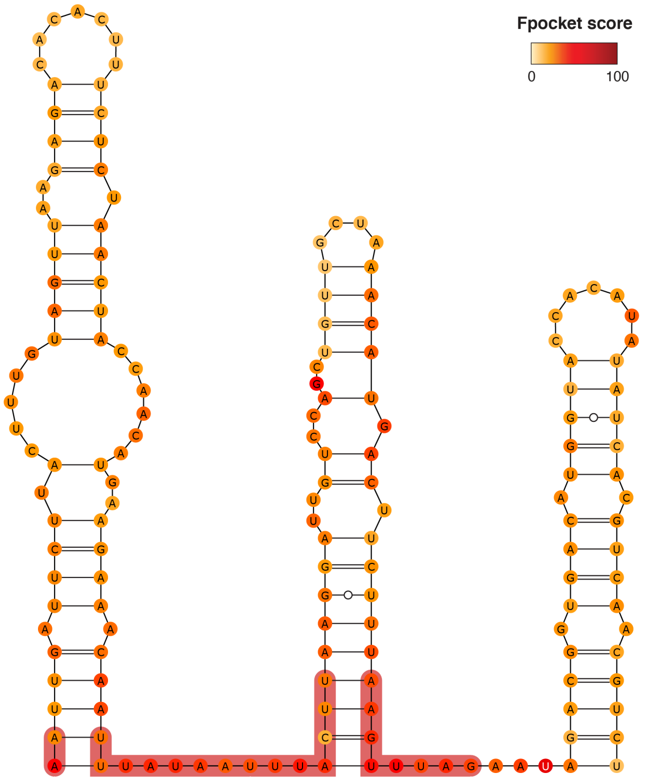


**Figure S28.** Consensus structure for segment 13635-13796, as derived from the 1000 lowest energy 3D structure structures modeled by SimRNA (using the MEA secondary structure inferred from SHAPE-MaP as a restraint). Bases are color-coded according to the Fpocket normalized score (0-100; see Methods). Residues composing the identified druggable pocket are shaded.

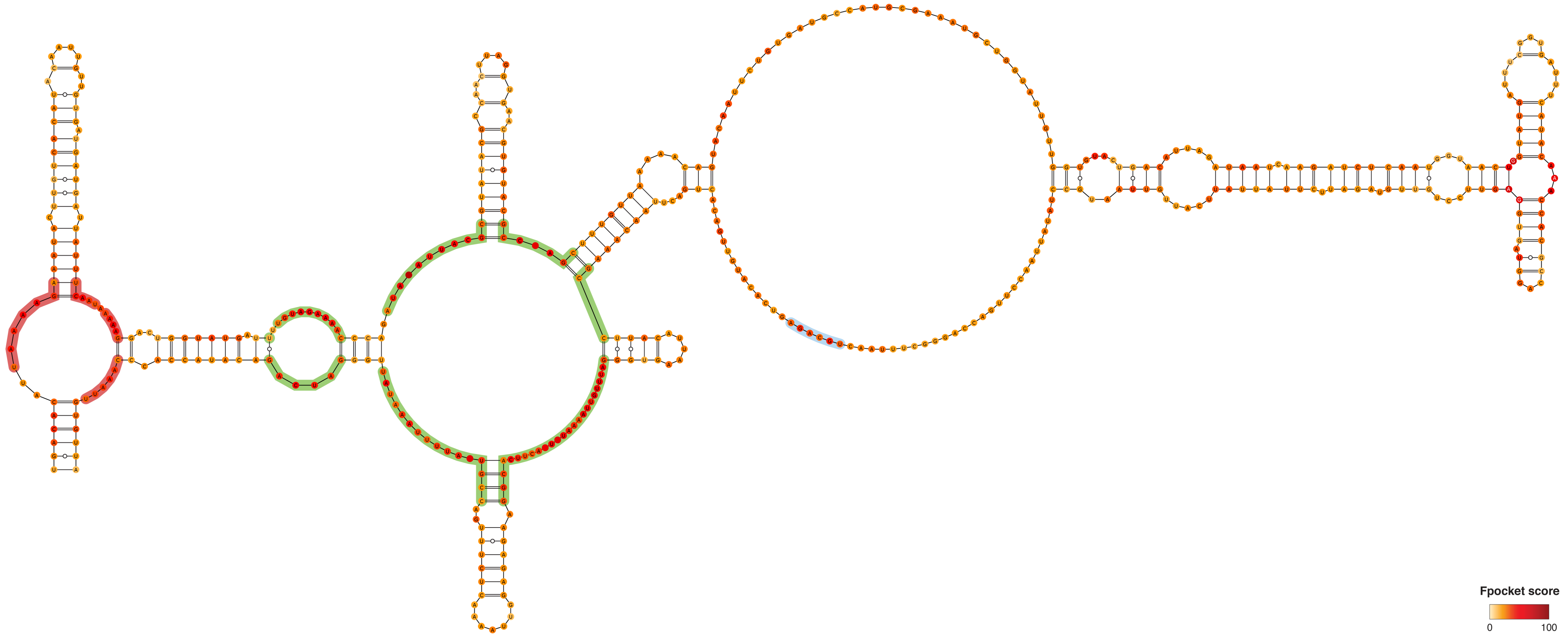


**Figure S29.** Consensus structure for segment 13857-14338, as derived from the 1000 lowest energy 3D structure structures modeled by SimRNA (using the MEA secondary structure inferred from SHAPE-MaP as a restraint). Bases are color-coded according to the Fpocket normalized score (0-100; see Methods). Residues composing the identified druggable pockets are shaded. Different shading colors mark distinct pockets.

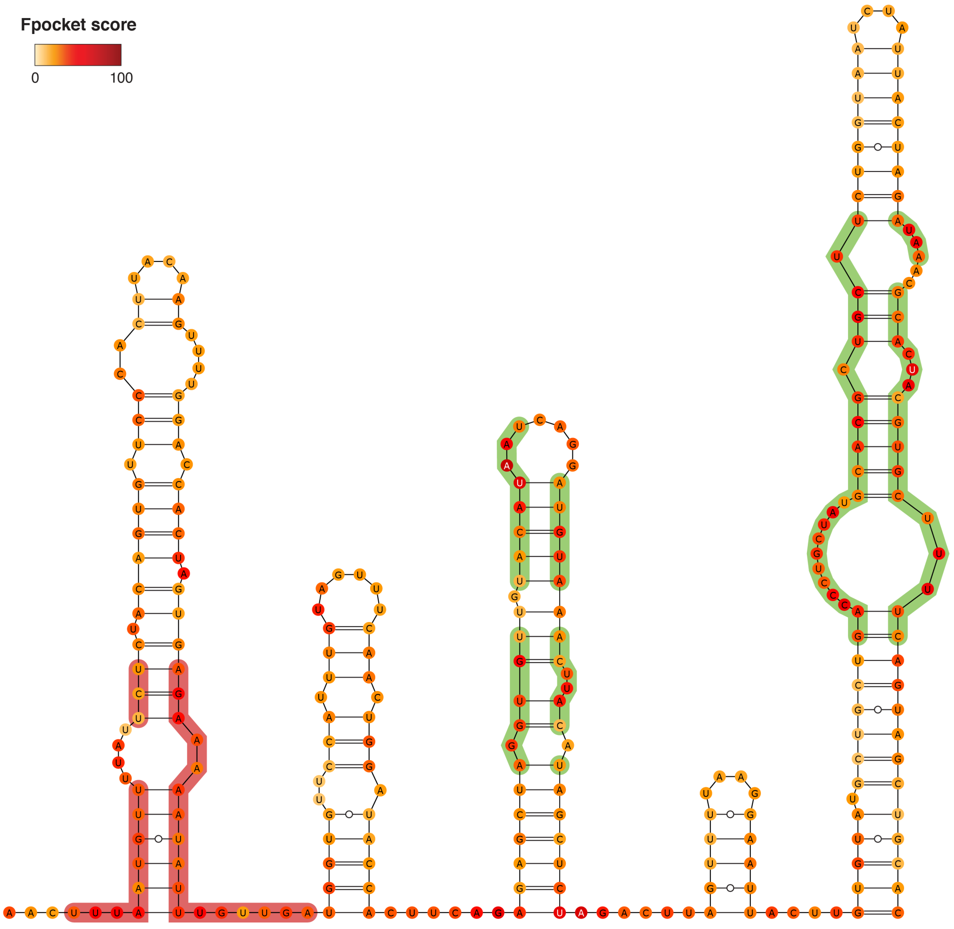


**Figure S30.** Consensus structure for segment 14374-14641, as derived from the 1000 lowest energy 3D structure structures modeled by SimRNA (using the MEA secondary structure inferred from SHAPE-MaP as a restraint). Bases are color-coded according to the Fpocket normalized score (0-100; see Methods). Residues composing the identified druggable pockets are shaded. Different shading colors mark distinct pockets.

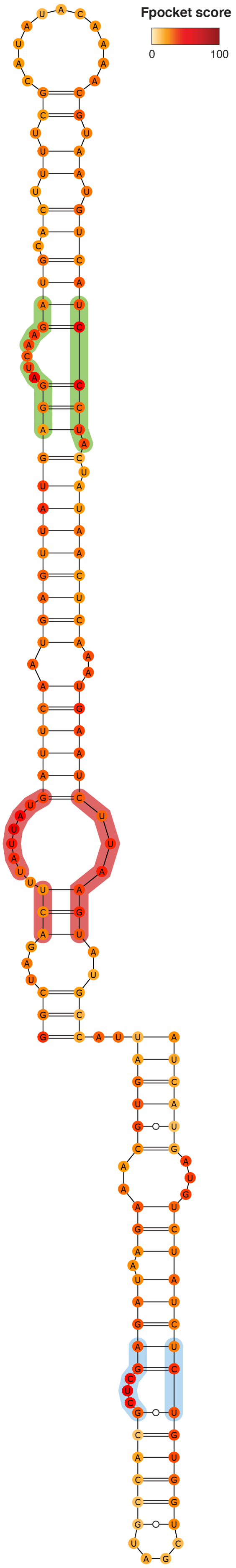


**Figure S31.** Consensus structure for segment 14973-15136, as derived from the 1000 lowest energy 3D structure structures modeled by SimRNA (using the MEA secondary structure inferred from SHAPE-MaP as a restraint). Bases are color-coded according to the Fpocket normalized score (0-100; see Methods). Residues composing the identified druggable pockets are shaded. Different shading colors mark distinct pockets.

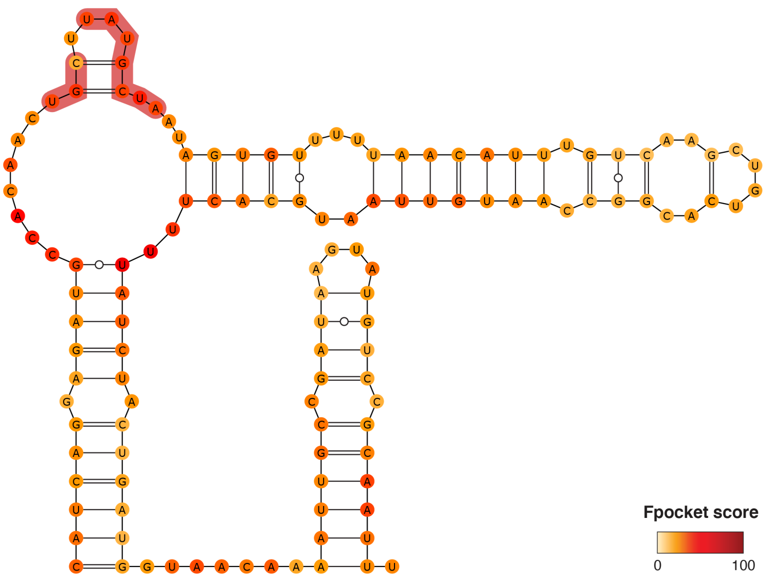


**Figure S32.** Consensus structure for segment 15482-15608, as derived from the 1000 lowest energy 3D structure structures modeled by SimRNA (using the MEA secondary structure inferred from SHAPE-MaP as a restraint). Bases are color-coded according to the Fpocket normalized score (0-100; see Methods). Residues composing the identified druggable pocket are shaded.

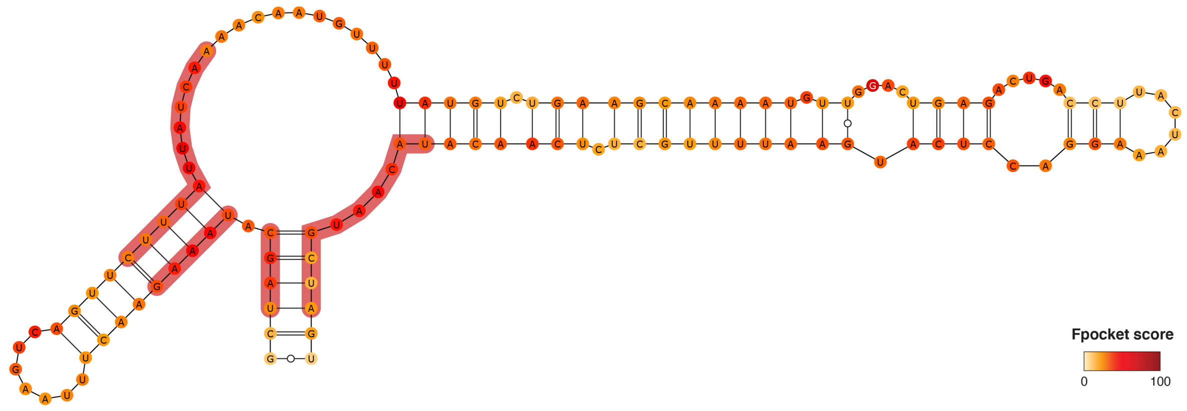


**Figure S33.** Consensus structure for segment 15769-15899, as derived from the 1000 lowest energy 3D structure structures modeled by SimRNA (using the MEA secondary structure inferred from SHAPE-MaP as a restraint). Bases are color-coded according to the Fpocket normalized score (0-100; see Methods). Residues composing the identified druggable pocket are shaded.

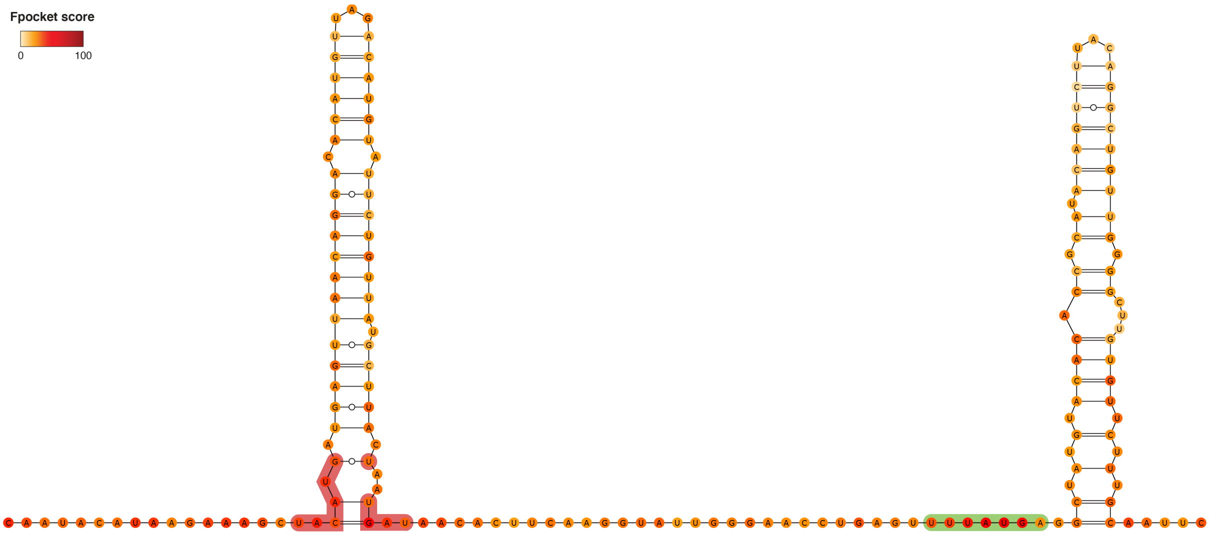


**Figure S34.** Consensus structure for segment 16096-16265, as derived from the 1000 lowest energy 3D structure structures modeled by SimRNA (using the MEA secondary structure inferred from SHAPE-MaP as a restraint). Bases are color-coded according to the Fpocket normalized score (0-100; see Methods). Residues composing the identified druggable pockets are shaded. Different shading colors mark distinct pockets.

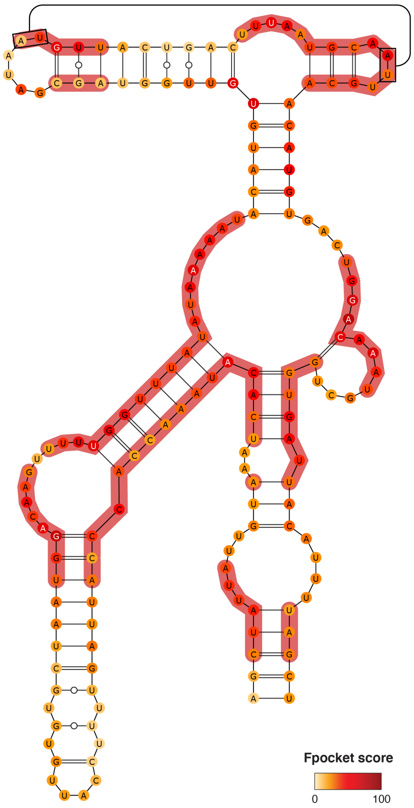


**Figure S35.** Consensus structure for segment 16441-16605, as derived from the 1000 lowest energy 3D structure structures modeled by SimRNA (using the MEA secondary structure inferred from SHAPE-MaP as a restraint). Bases are color-coded according to the Fpocket normalized score (0-100; see Methods). Residues composing the identified druggable pocket are shaded.

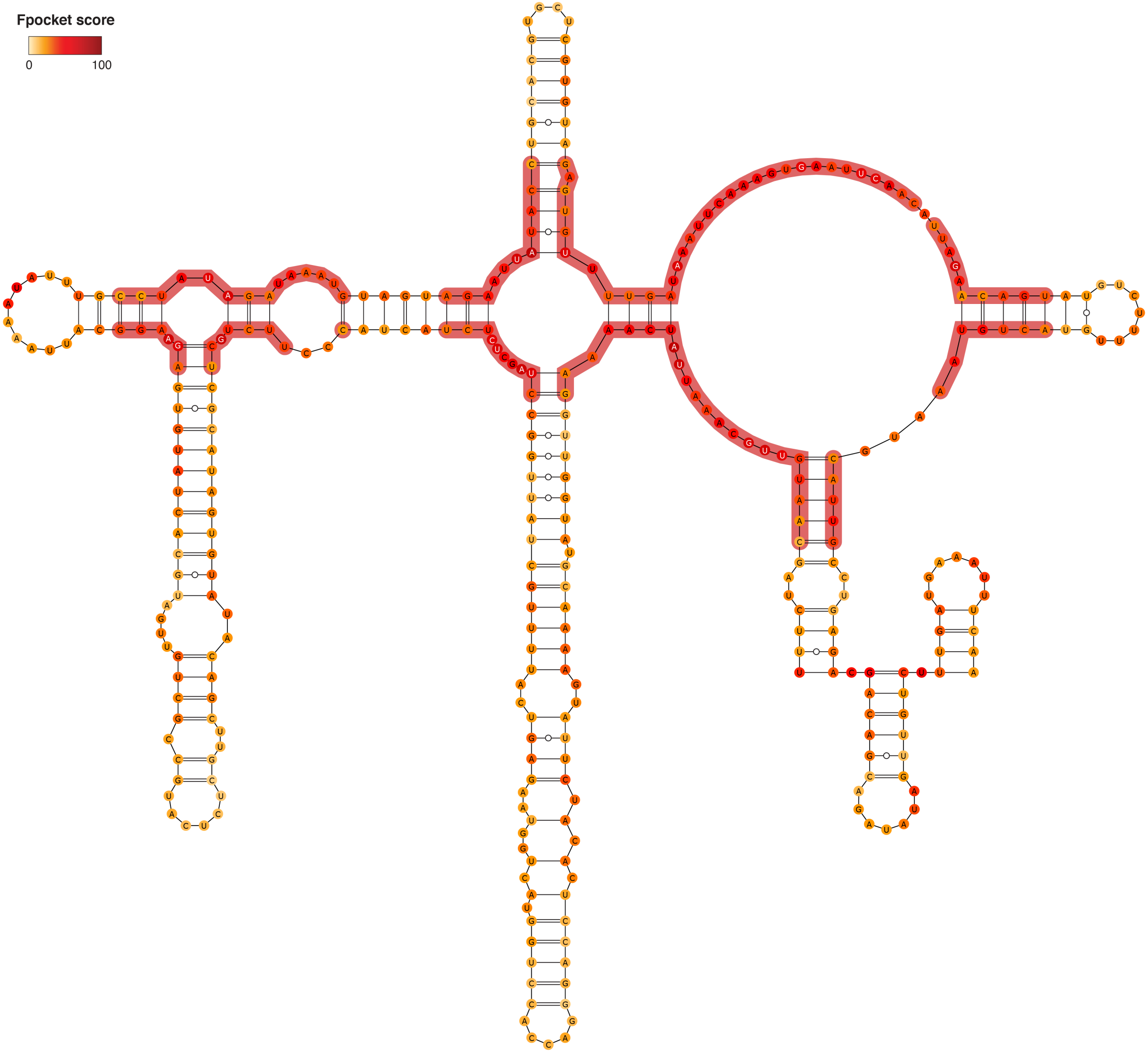


**Figure S36.** Consensus structure for segment 17021-17368, as derived from the 1000 lowest energy 3D structure structures modeled by SimRNA (using the MEA secondary structure inferred from SHAPE-MaP as a restraint). Bases are color-coded according to the Fpocket normalized score (0-100; see Methods). Residues composing the identified druggable pocket are shaded.

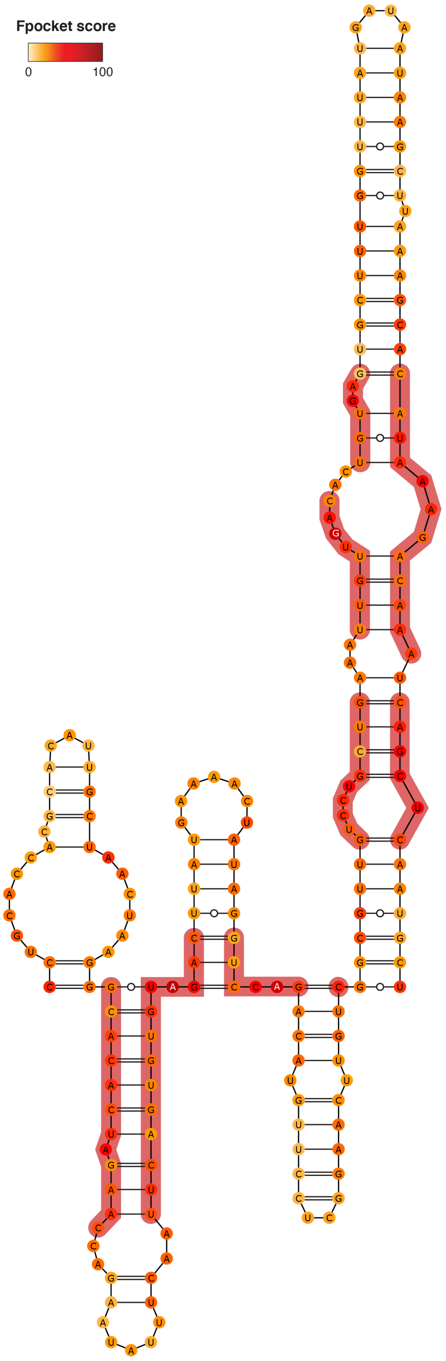


**Figure S37.** Consensus structure for segment 17452-17650, as derived from the 1000 lowest energy 3D structure structures modeled by SimRNA (using the MEA secondary structure inferred from SHAPE-MaP as a restraint). Bases are color-coded according to the Fpocket normalized score (0-100; see Methods). Residues composing the identified druggable pocket are shaded.

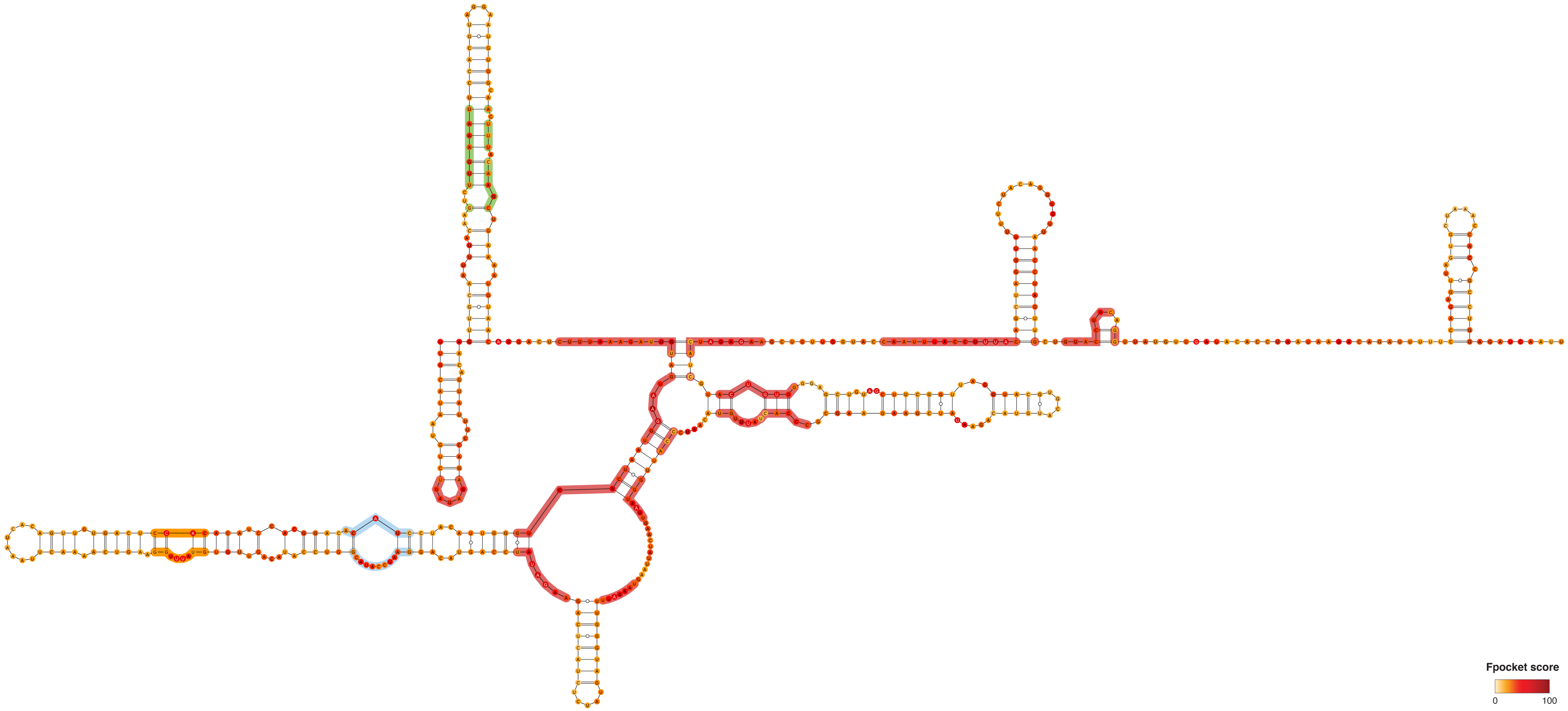


**Figure S38.** Consensus structure for segment 17955-18476, as derived from the 1000 lowest energy 3D structure structures modeled by SimRNA (using the MEA secondary structure inferred from SHAPE-MaP as a restraint). Bases are color-coded according to the Fpocket normalized score (0-100; see Methods). Residues composing the identified druggable pockets are shaded. Different shading colors mark distinct pockets.

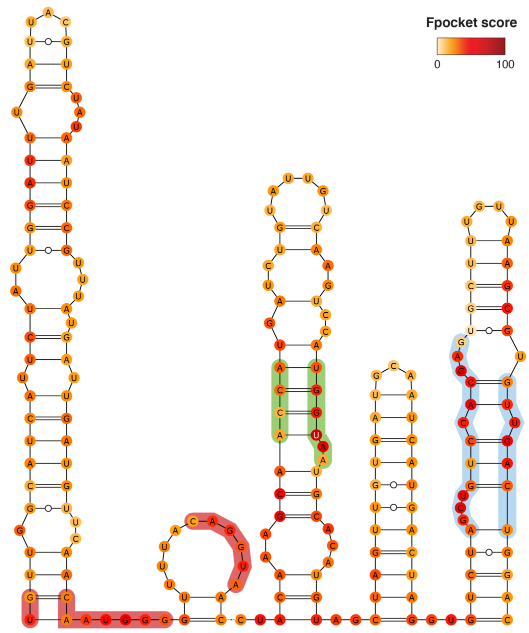


**Figure S39.** Consensus structure for segment 18715-18917, as derived from the 1000 lowest energy 3D structure structures modeled by SimRNA (using the MEA secondary structure inferred from SHAPE-MaP as a restraint). Bases are color-coded according to the Fpocket normalized score (0-100; see Methods). Residues composing the identified druggable pockets are shaded. Different shading colors mark distinct pockets.

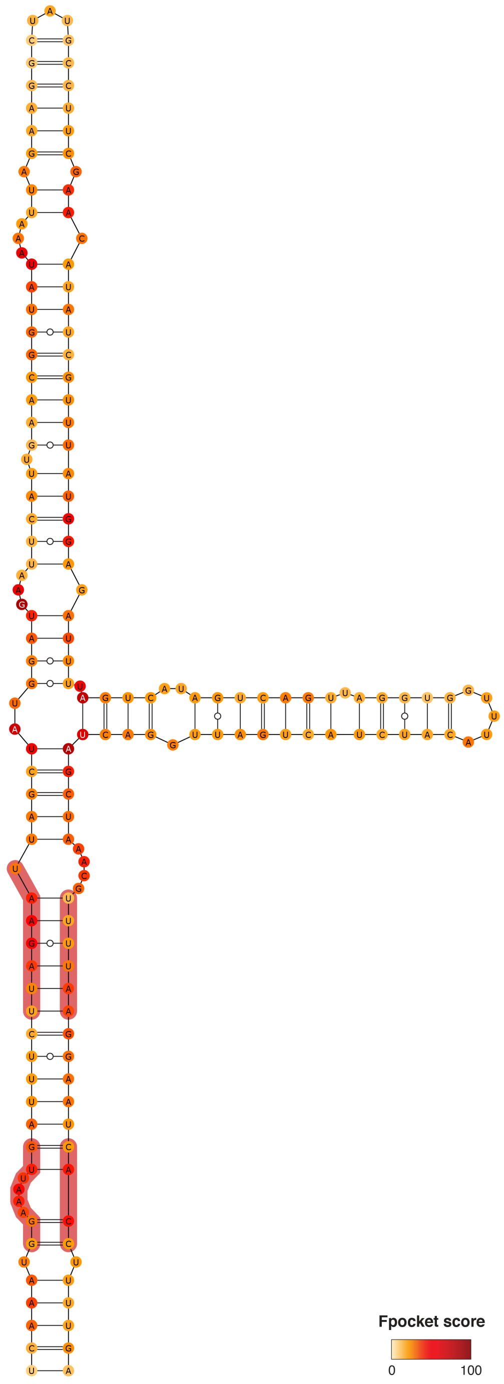


**Figure S40.** Consensus structure for segment 20241-20411, as derived from the 1000 lowest energy 3D structure structures modeled by SimRNA (using the MEA secondary structure inferred from SHAPE-MaP as a restraint). Bases are color-coded according to the Fpocket normalized score (0-100; see Methods). Residues composing the identified druggable pocket are shaded.

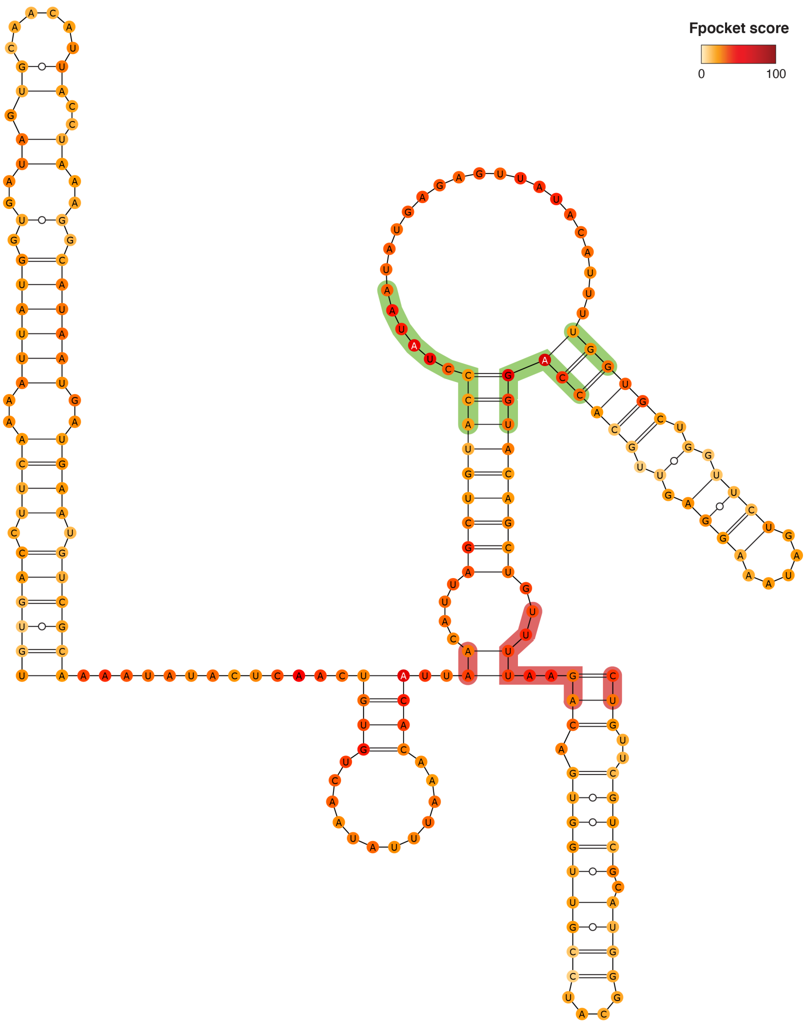


**Figure S41.** Consensus structure for segment 20731-20946, as derived from the 1000 lowest energy 3D structure structures modeled by SimRNA (using the MEA secondary structure inferred from SHAPE-MaP as a restraint). Bases are color-coded according to the Fpocket normalized score (0-100; see Methods). Residues composing the identified druggable pockets are shaded. Different shading colors mark distinct pockets.

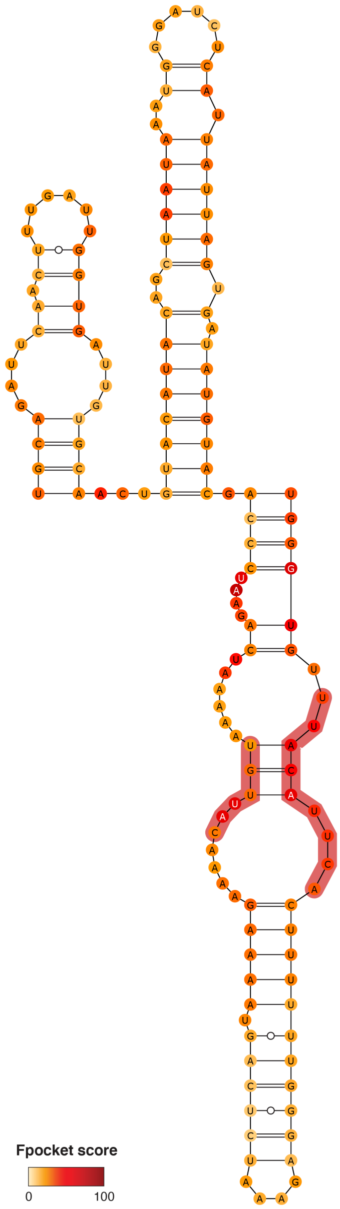


**Figure S42.** Consensus structure for segment 20976-21124, as derived from the 1000 lowest energy 3D structure structures modeled by SimRNA (using the MEA secondary structure inferred from SHAPE-MaP as a restraint). Bases are color-coded according to the Fpocket normalized score (0-100; see Methods). Residues composing the identified druggable pocket are shaded.

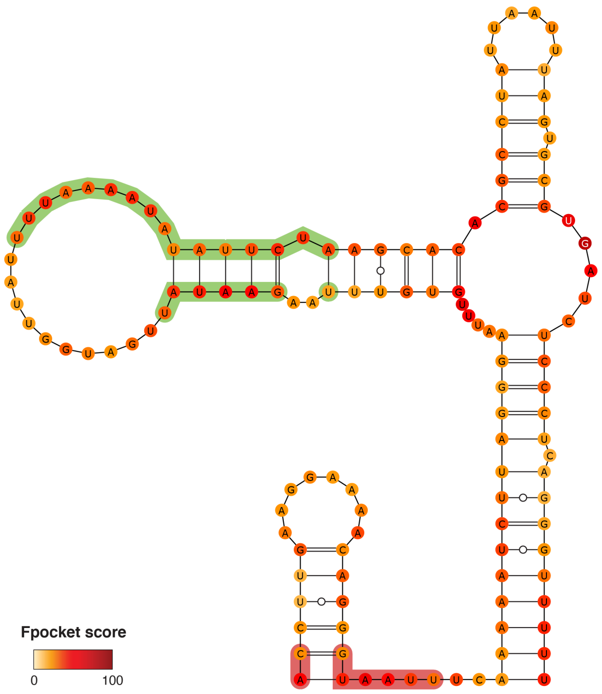


**Figure S43.** Consensus structure for segment 22095-22223, as derived from the 1000 lowest energy 3D structure structures modeled by SimRNA (using the MEA secondary structure inferred from SHAPE-MaP as a restraint). Bases are color-coded according to the Fpocket normalized score (0-100; see Methods). Residues composing the identified druggable pockets are shaded. Different shading colors mark distinct pockets.

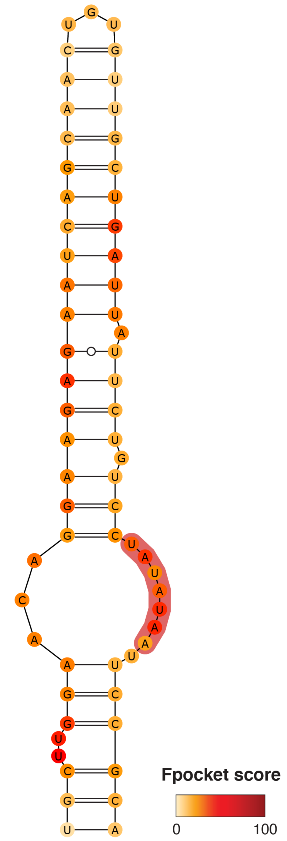


**Figure S44.** Consensus structure for segment 22615-22678, as derived from the 1000 lowest energy 3D structure structures modeled by SimRNA (using the MEA secondary structure inferred from SHAPE-MaP as a restraint). Bases are color-coded according to the Fpocket normalized score (0-100; see Methods). Residues composing the identified druggable pocket are shaded.

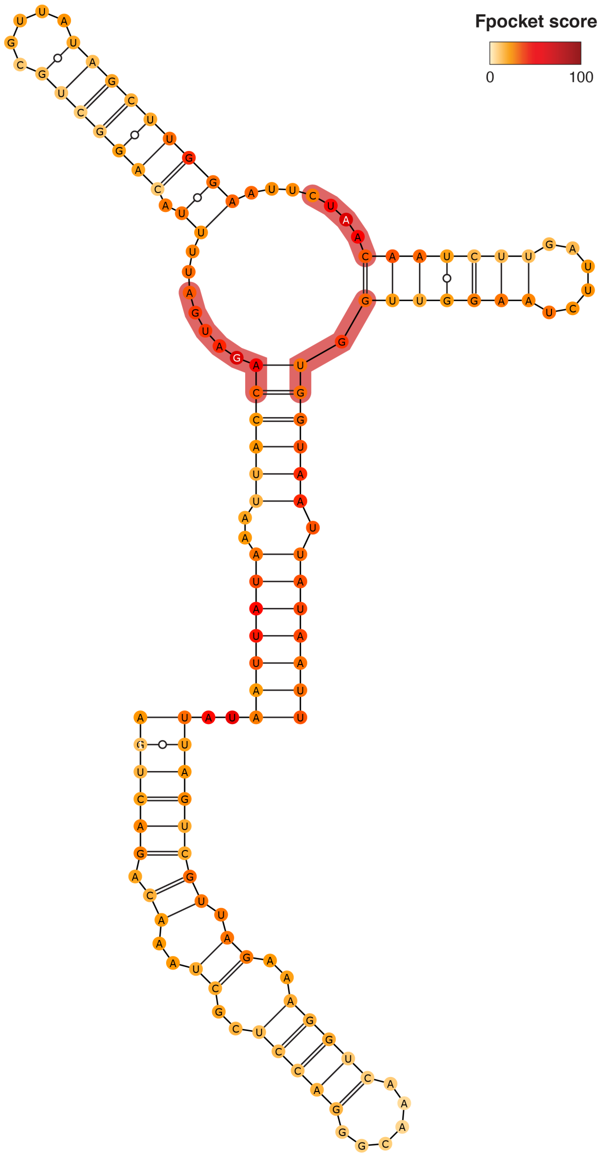


**Figure S45.** Consensus structure for segment 22780-22913, as derived from the 1000 lowest energy 3D structure structures modeled by SimRNA (using the MEA secondary structure inferred from SHAPE-MaP as a restraint). Bases are color-coded according to the Fpocket normalized score (0-100; see Methods). Residues composing the identified druggable pocket are shaded.

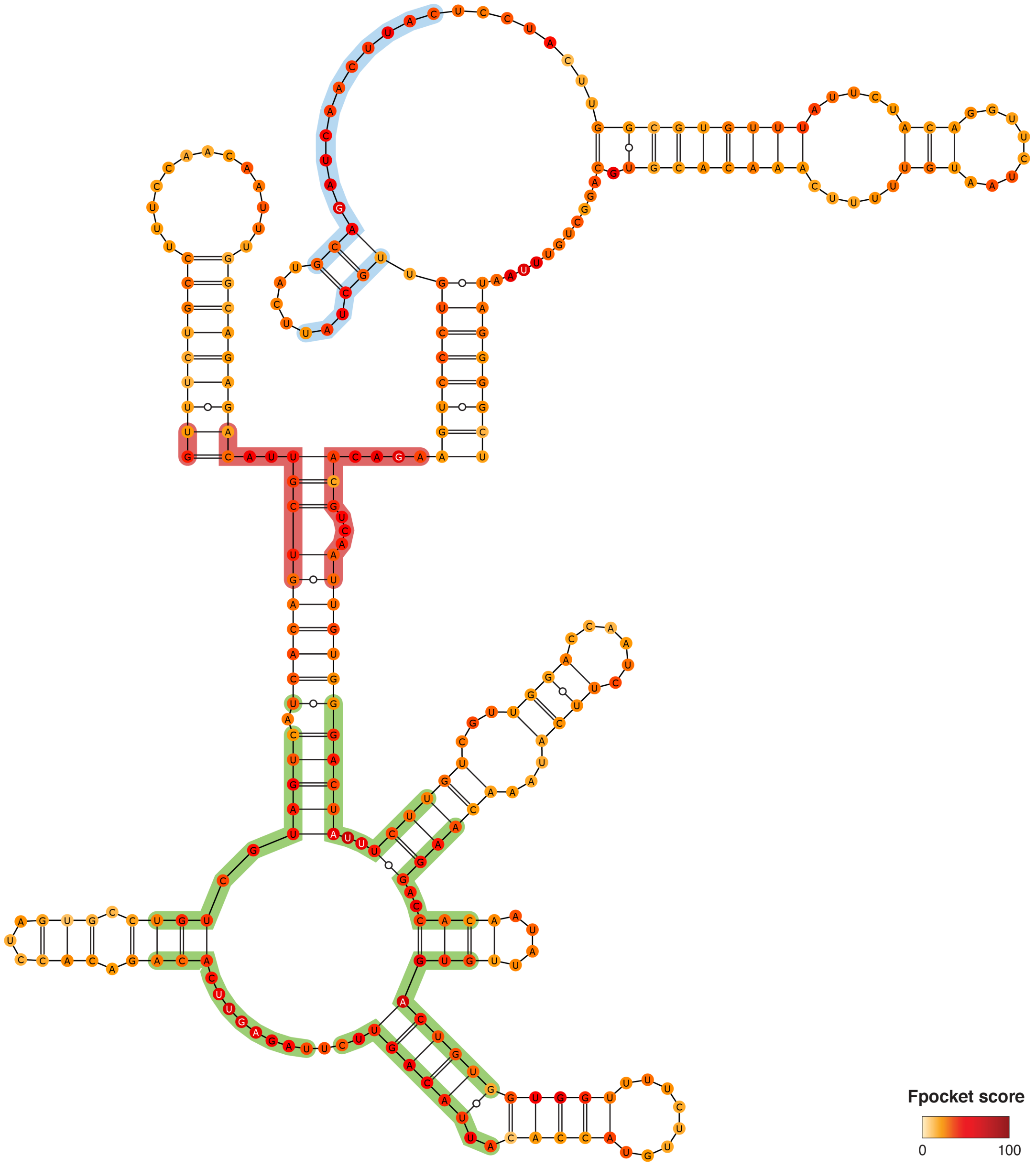


**Figure S46.** Consensus structure for segment 23236-23521, as derived from the 1000 lowest energy 3D structure structures modeled by SimRNA (using the MEA secondary structure inferred from SHAPE-MaP as a restraint). Bases are color-coded according to the Fpocket normalized score (0-100; see Methods). Residues composing the identified druggable pockets are shaded. Different shading colors mark distinct pockets.

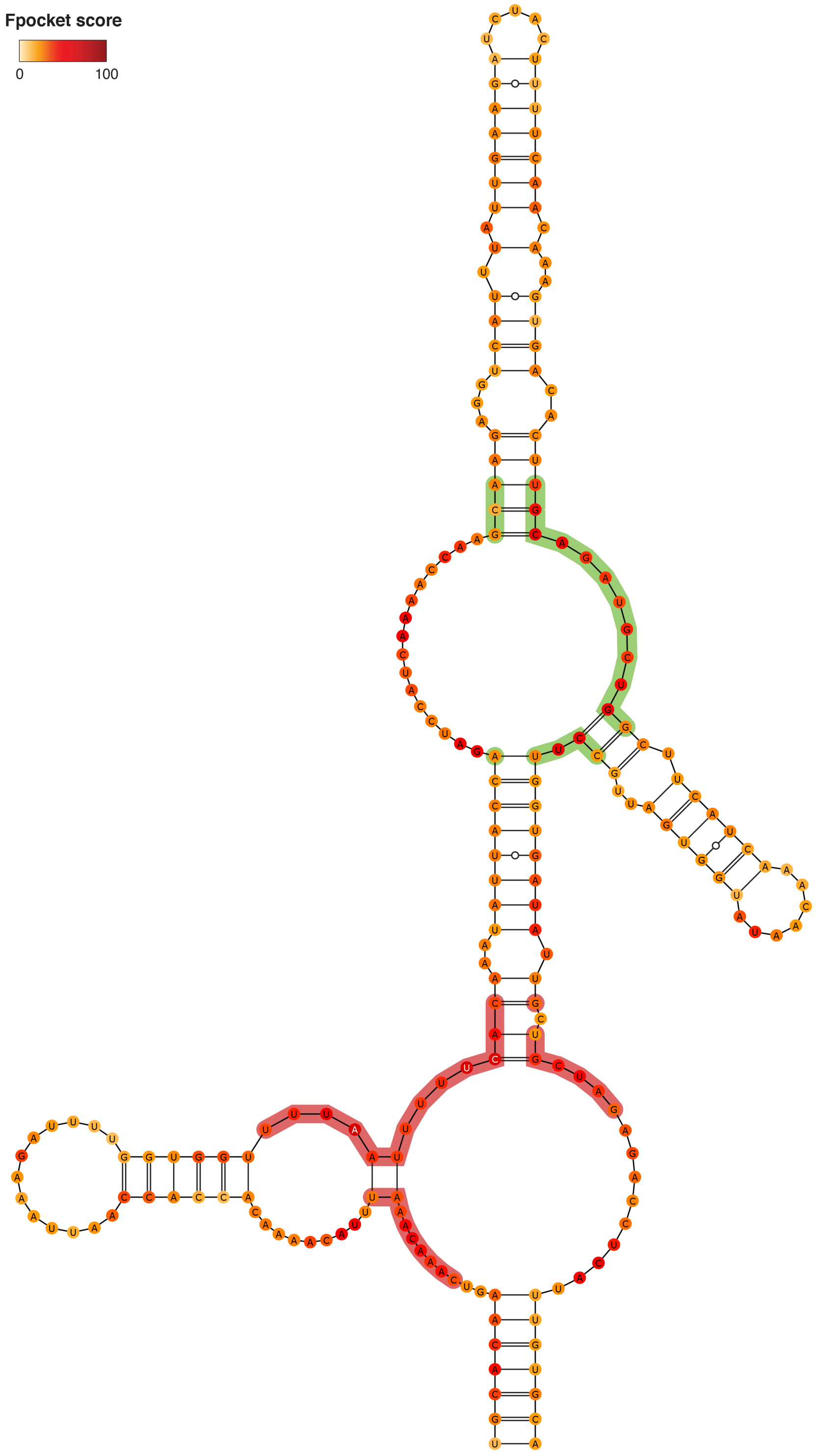


**Figure S47.** Consensus structure for segment 23908-24118, as derived from the 1000 lowest energy 3D structure structures modeled by SimRNA (using the MEA secondary structure inferred from SHAPE-MaP as a restraint). Bases are color-coded according to the Fpocket normalized score (0-100; see Methods). Residues composing the identified druggable pockets are shaded. Different shading colors mark distinct pockets.

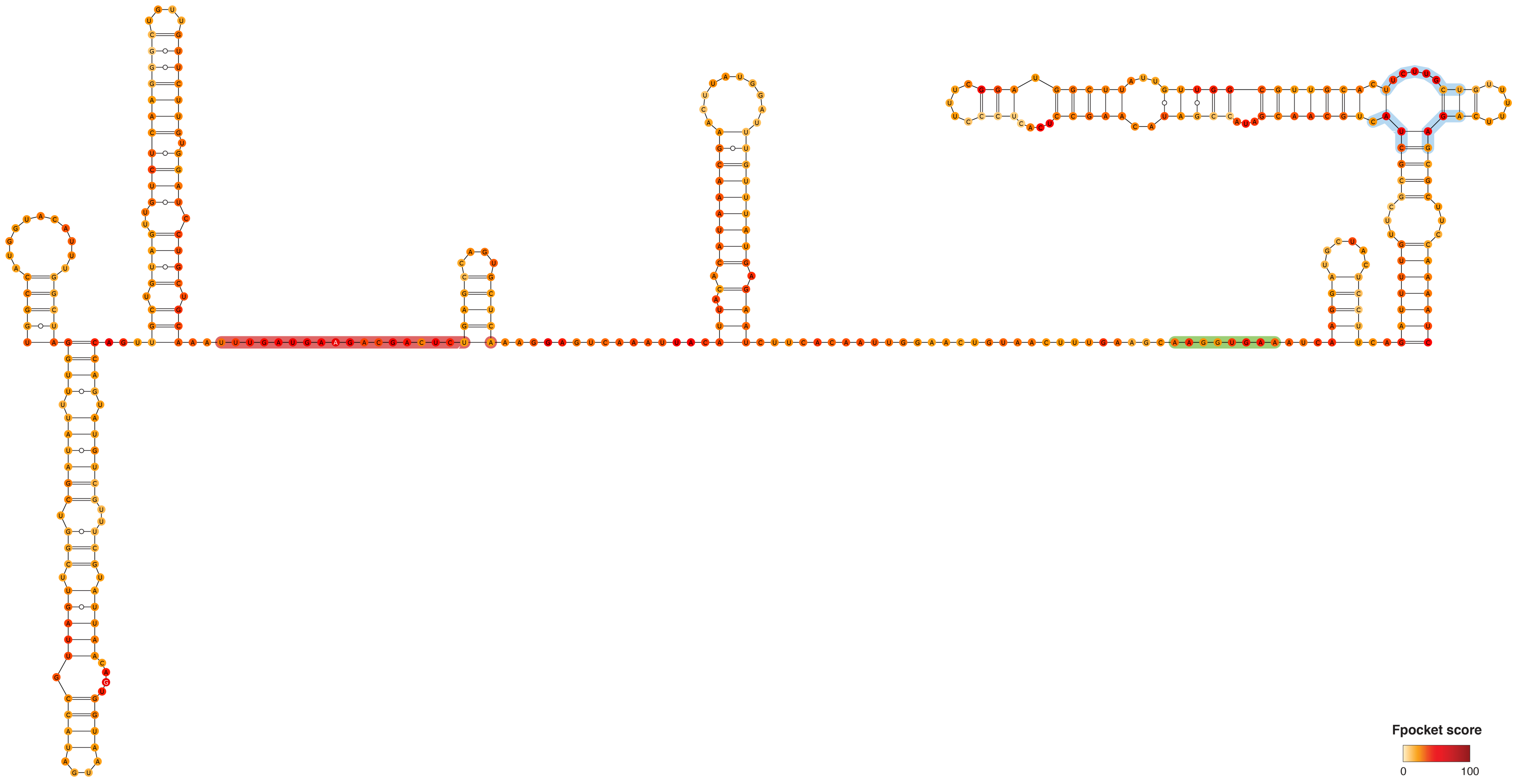


**Figure S48.** Consensus structure for segment 25196-25578, as derived from the 1000 lowest energy 3D structure structures modeled by SimRNA (using the MEA secondary structure inferred from SHAPE-MaP as a restraint). Bases are color-coded according to the Fpocket normalized score (0-100; see Methods). Residues composing the identified druggable pockets are shaded. Different shading colors mark distinct pockets.

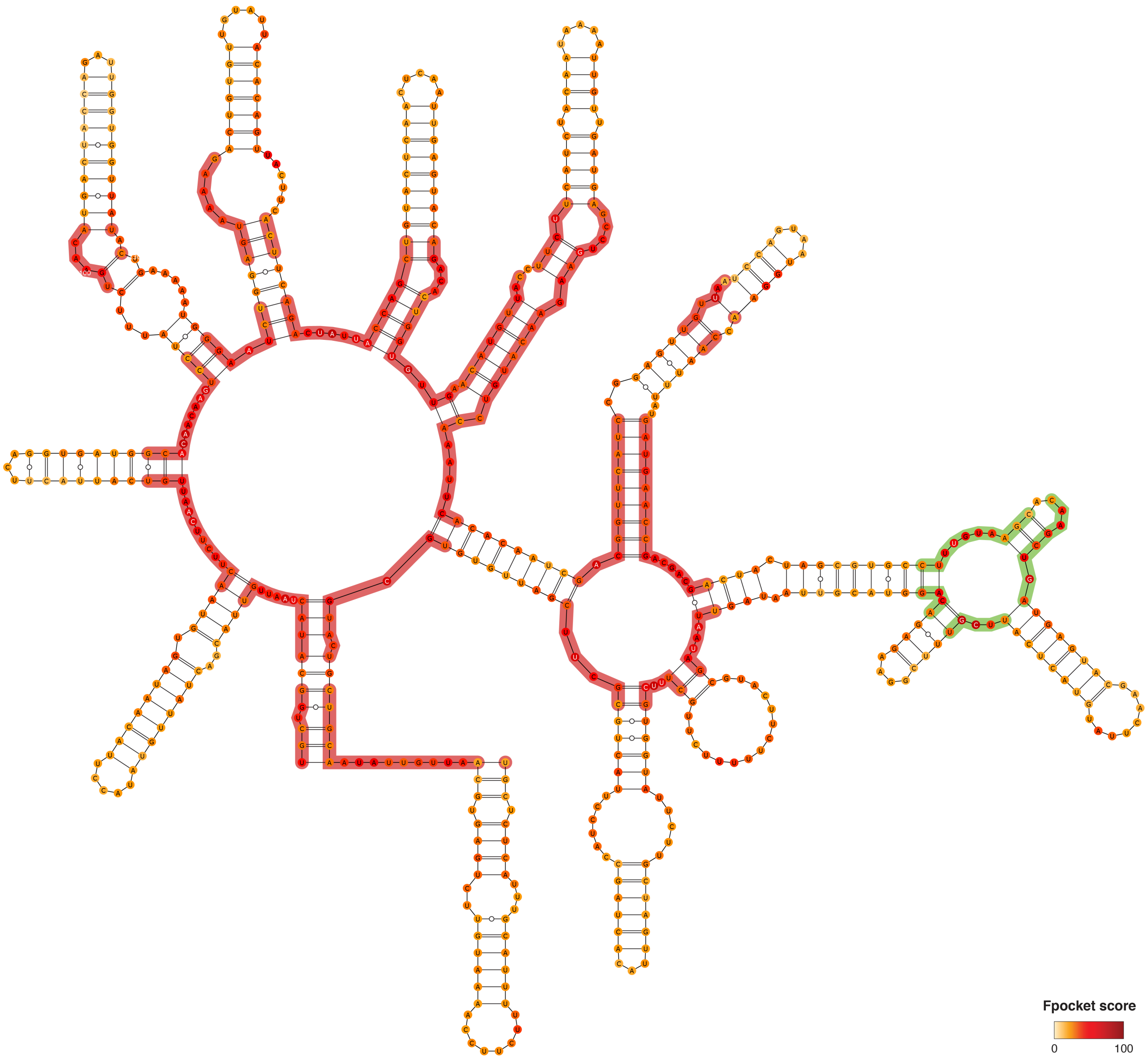


**Figure S49.** Consensus structure for segment 25834-26427, as derived from the 1000 lowest energy 3D structure structures modeled by SimRNA (using the MEA secondary structure inferred from SHAPE-MaP as a restraint). Bases are color-coded according to the Fpocket normalized score (0-100; see Methods). Residues composing the identified druggable pockets are shaded. Different shading colors mark distinct pockets.

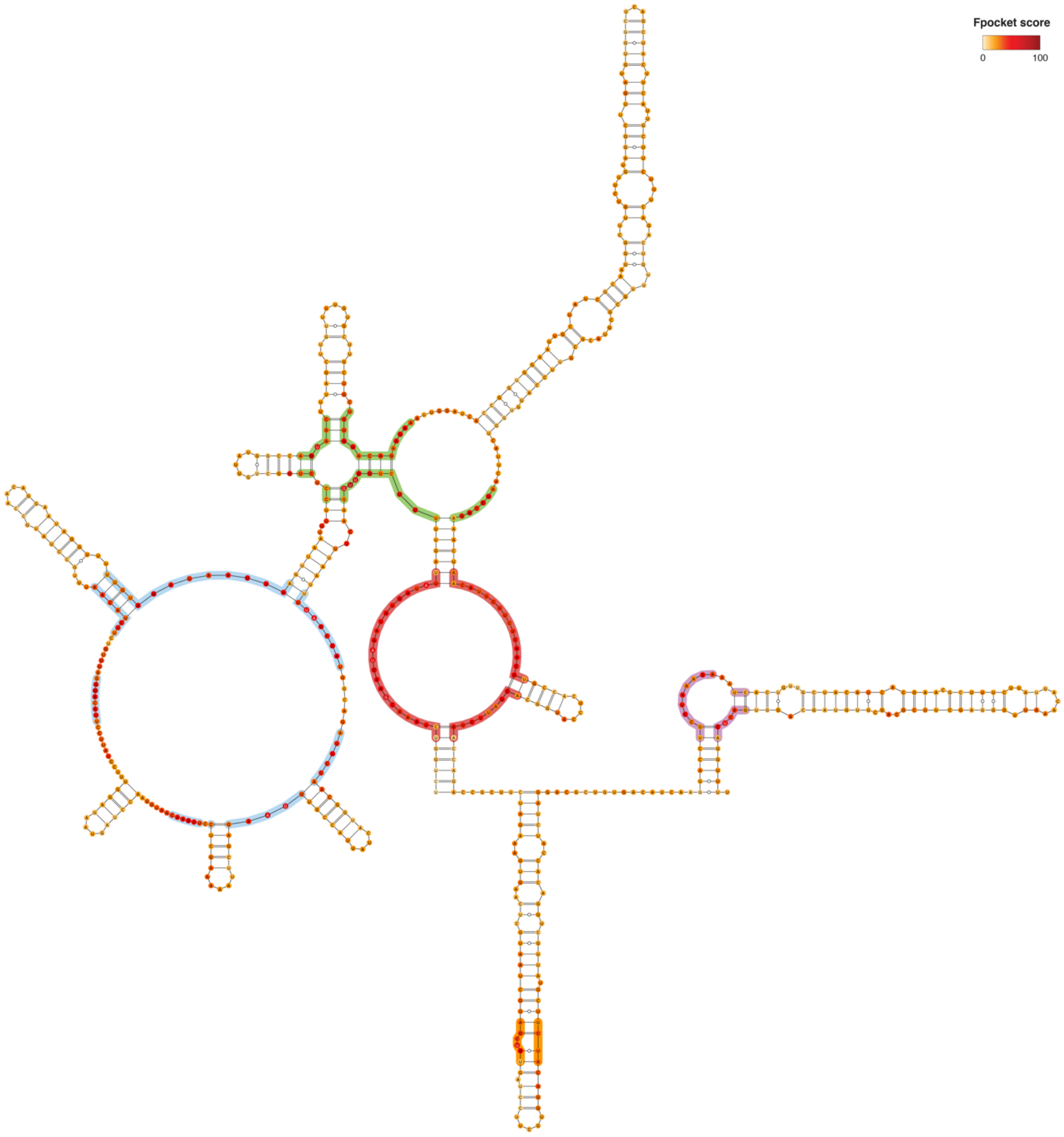


**Figure S50.** Consensus structure for segment 26463-27101, as derived from the 1000 lowest energy 3D structure structures modeled by SimRNA (using the MEA secondary structure inferred from SHAPE-MaP as a restraint). Bases are color-coded according to the Fpocket normalized score (0-100; see Methods). Residues composing the identified druggable pockets are shaded. Different shading colors mark distinct pockets.

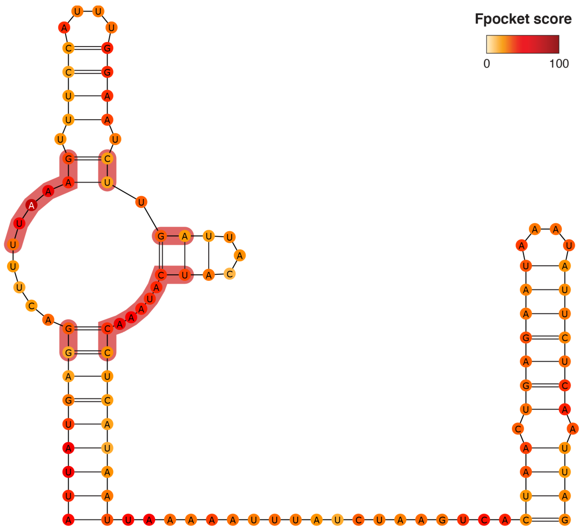


**Figure S51.** Consensus structure for segment 27253-27358, as derived from the 1000 lowest energy 3D structure structures modeled by SimRNA (using the MEA secondary structure inferred from SHAPE-MaP as a restraint). Bases are color-coded according to the Fpocket normalized score (0-100; see Methods). Residues composing the identified druggable pocket are shaded.

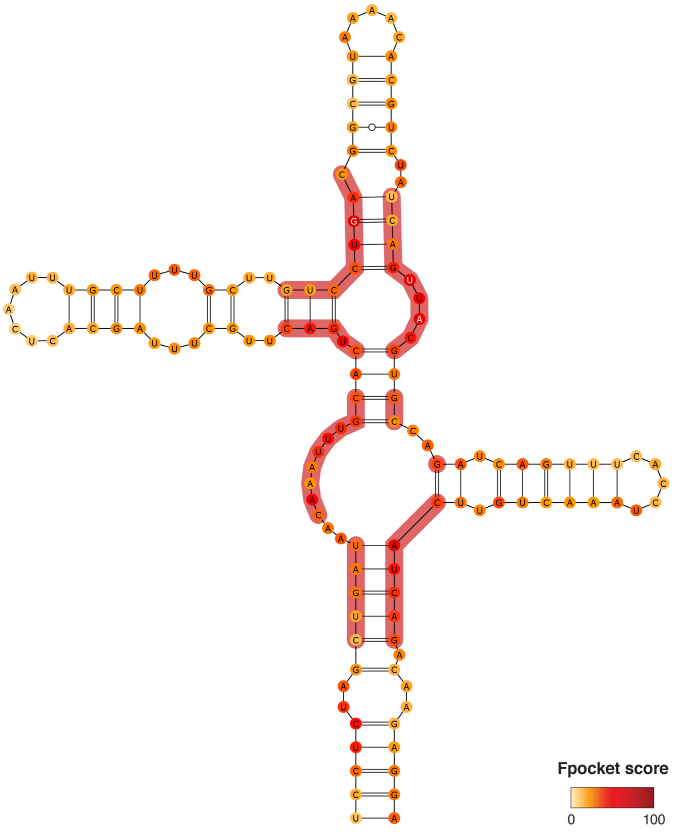


**Figure S52.** Consensus structure for segment 27534-27668, as derived from the 1000 lowest energy 3D structure structures modeled by SimRNA (using the MEA secondary structure inferred from SHAPE-MaP as a restraint). Bases are color-coded according to the Fpocket normalized score (0-100; see Methods). Residues composing the identified druggable pocket are shaded.

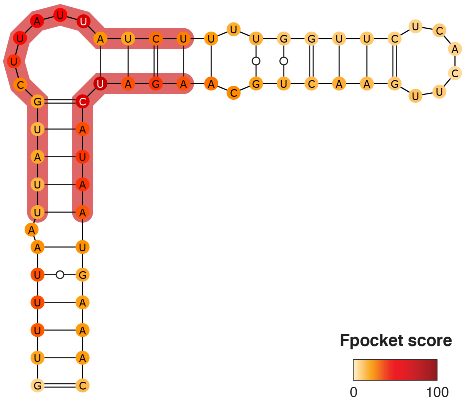


**Figure S53.** Consensus structure for segment 27816-27874, as derived from the 1000 lowest energy 3D structure structures modeled by SimRNA (using the MEA secondary structure inferred from SHAPE-MaP as a restraint). Bases are color-coded according to the Fpocket normalized score (0-100; see Methods). Residues composing the identified druggable pocket are shaded.

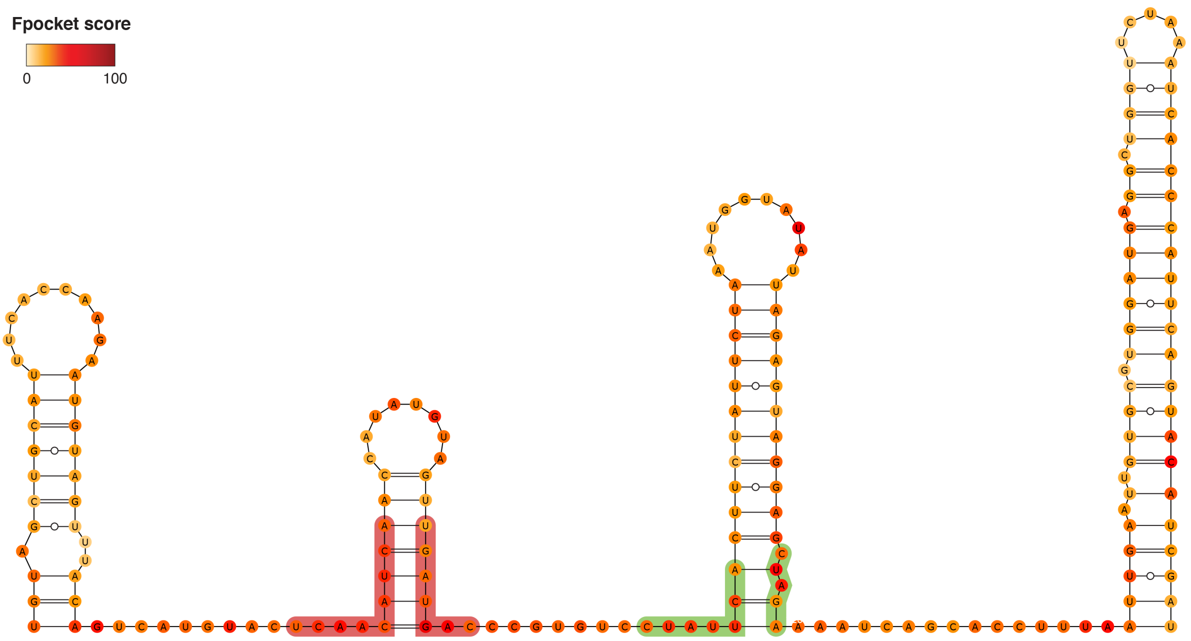


**Figure S54.** Consensus structure for segment 27929-28118, as derived from the 1000 lowest energy 3D structure structures modeled by SimRNA (using the MEA secondary structure inferred from SHAPE-MaP as a restraint). Bases are color-coded according to the Fpocket normalized score (0-100; see Methods). Residues composing the identified druggable pockets are shaded. Different shading colors mark distinct pockets.

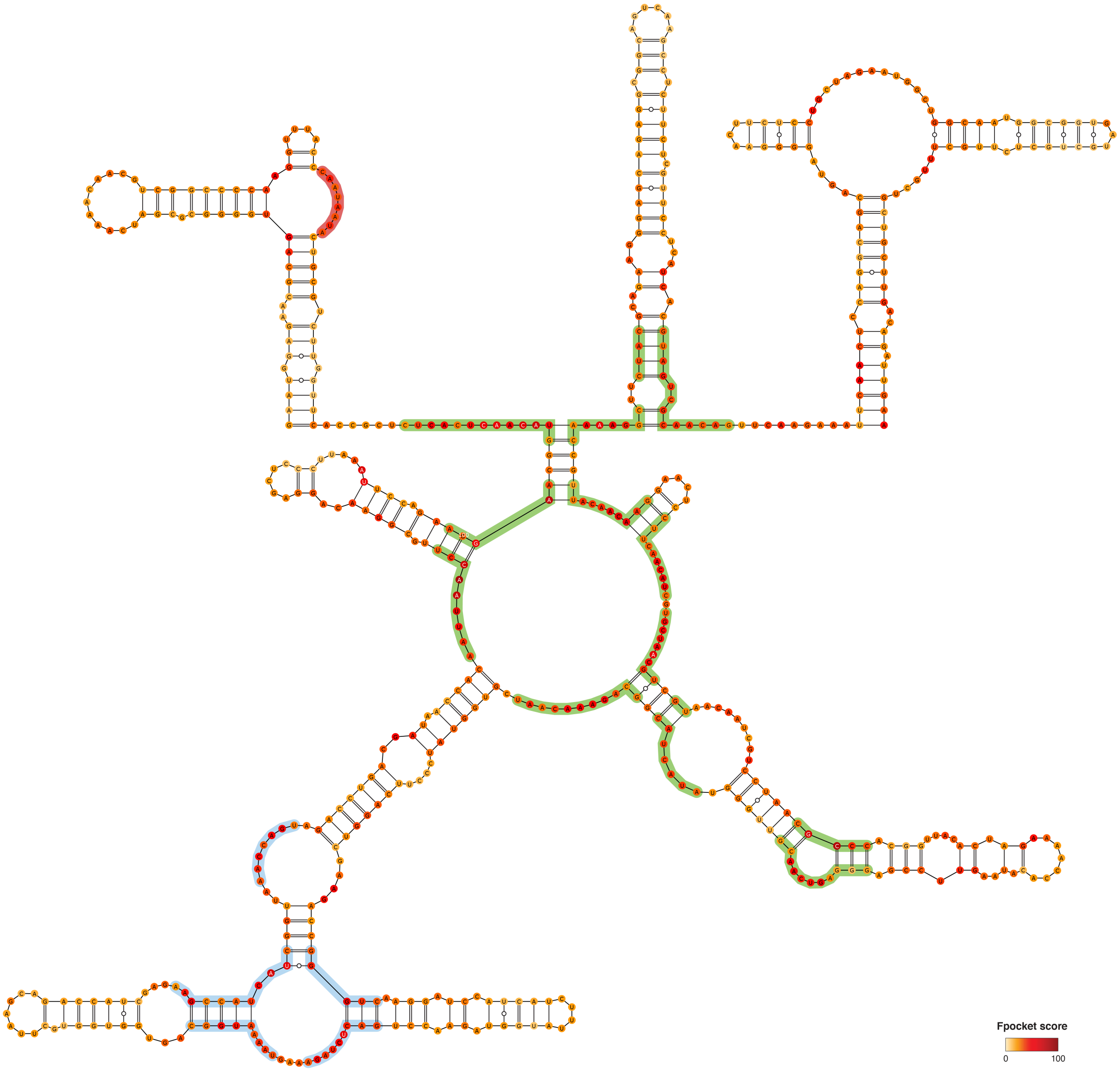


**Figure S55.** Consensus structure for segment 28357-28956, as derived from the 1000 lowest energy 3D structure structures modeled by SimRNA (using the MEA secondary structure inferred from SHAPE-MaP as a restraint). Bases are color-coded according to the Fpocket normalized score (0-100; see Methods). Residues composing the identified druggable pockets are shaded. Different shading colors mark distinct pockets.

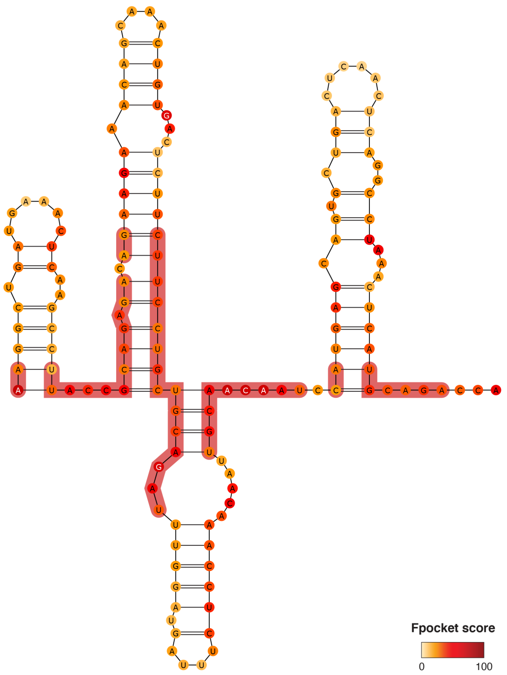


**Figure S56.** Consensus structure for segment 29396-29547, as derived from the 1000 lowest energy 3D structure structures modeled by SimRNA (using the MEA secondary structure inferred from SHAPE-MaP as a restraint). Bases are color-coded according to the Fpocket normalized score (0-100; see Methods). Residues composing the identified druggable pocket are shaded.

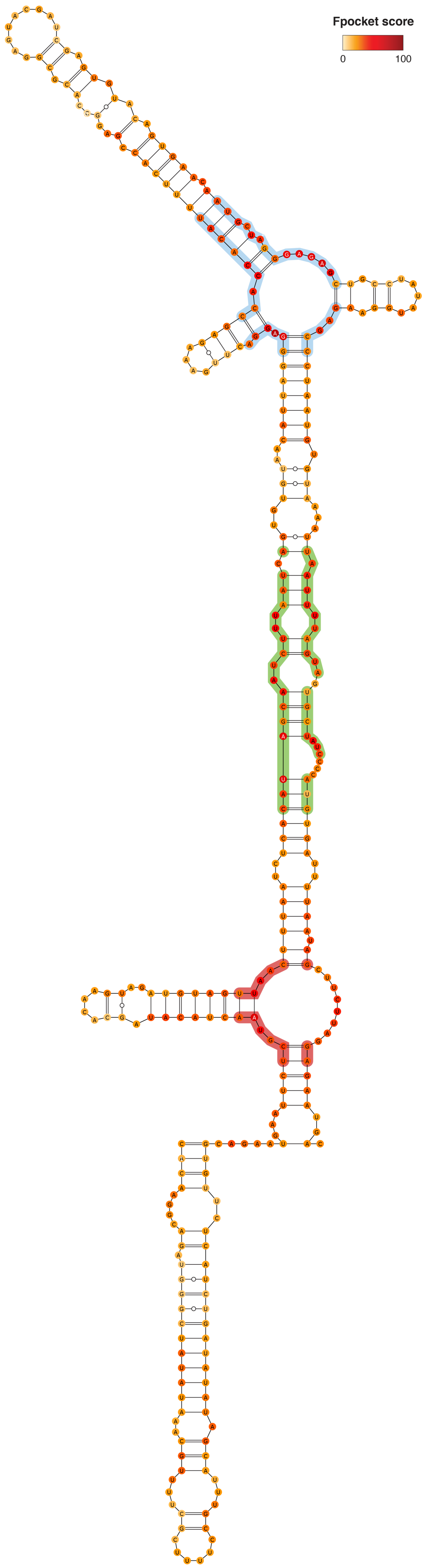


**Figure S57.** Consensus structure for segment 29548-29870, as derived from the 1000 lowest energy 3D structure structures modeled by SimRNA (using the MEA secondary structure inferred from SHAPE-MaP as a restraint). Bases are color-coded according to the Fpocket normalized score (0-100; see Methods). Residues composing the identified druggable pockets are shaded. Different shading colors mark distinct pockets.

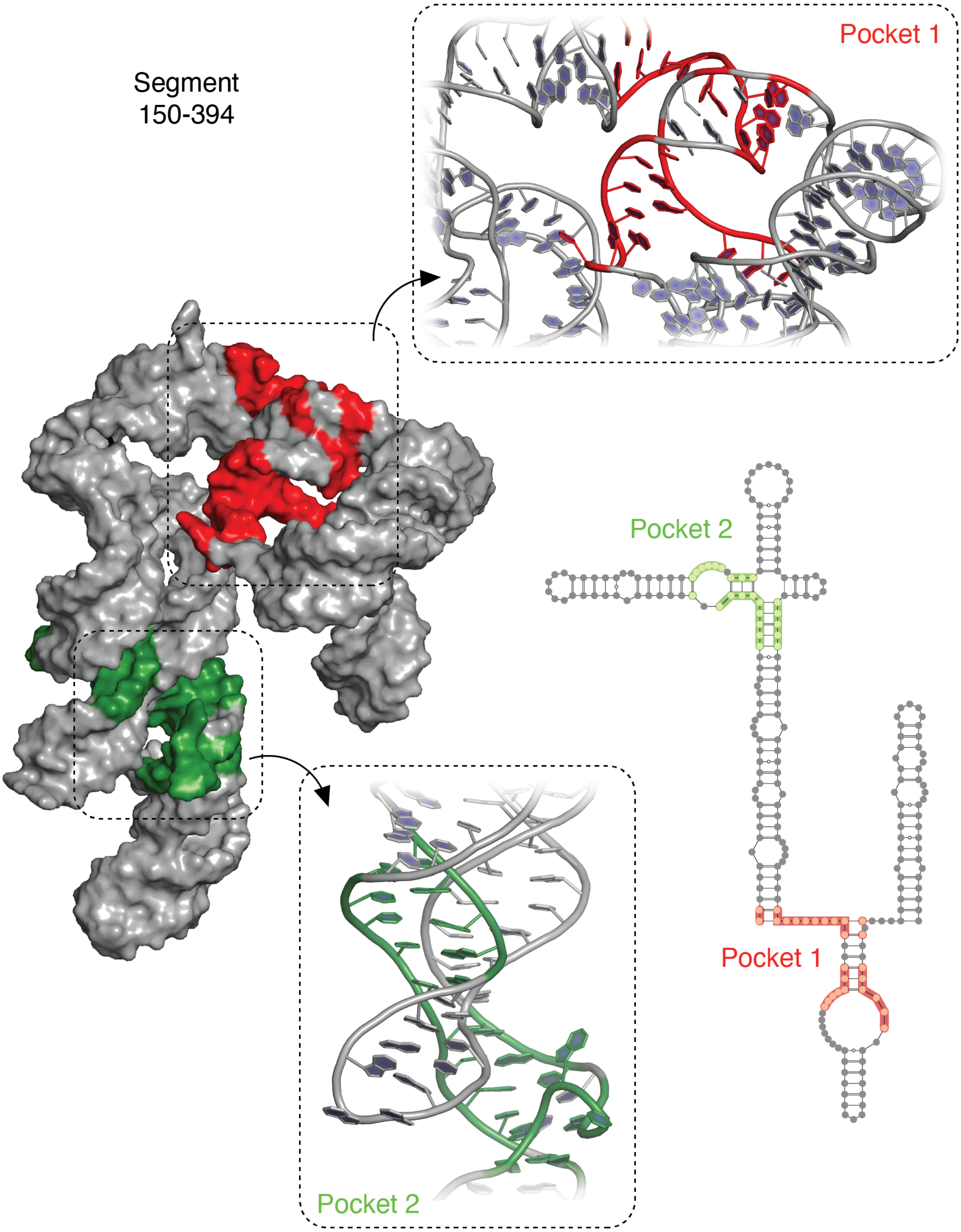


**Figure S58.** Consensus secondary structure for segment 150-394, as derived from the 1000 lowest energy 3D structure structures modeled by SimRNA (using the MEA secondary structure inferred from SHAPE-MaP as a restraint). Residues composing the identified druggable pockets are shaded. Different shading colors mark distinct pockets. The 3D model of the medoid structure from the most abundant cluster is also shown, with pockets colored as for the secondary structure. The insets show the atomic representation of the identified pockets.

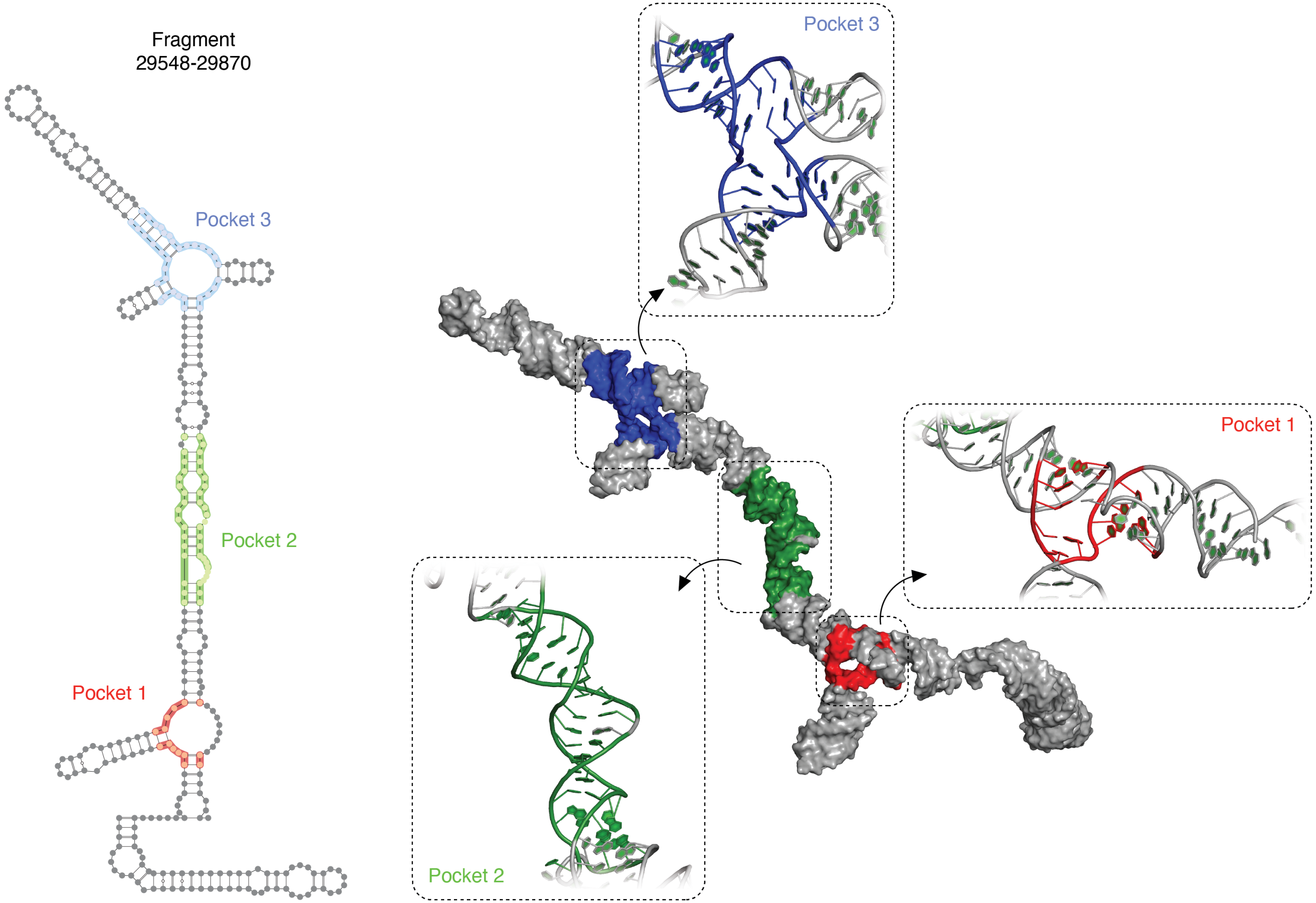


**Figure S59.** Consensus secondary structure for segment 29548-29870, as derived from the 1000 lowest energy 3D structure structures modeled by SimRNA (using the MEA secondary structure inferred from SHAPE-MaP as a restraint). Residues composing the identified druggable pockets are shaded. Different shading colors mark distinct pockets. The 3D model of the medoid structure from the most abundant cluster is also shown, with pockets colored as for the secondary structure. The insets show the atomic representation of the identified pockets.

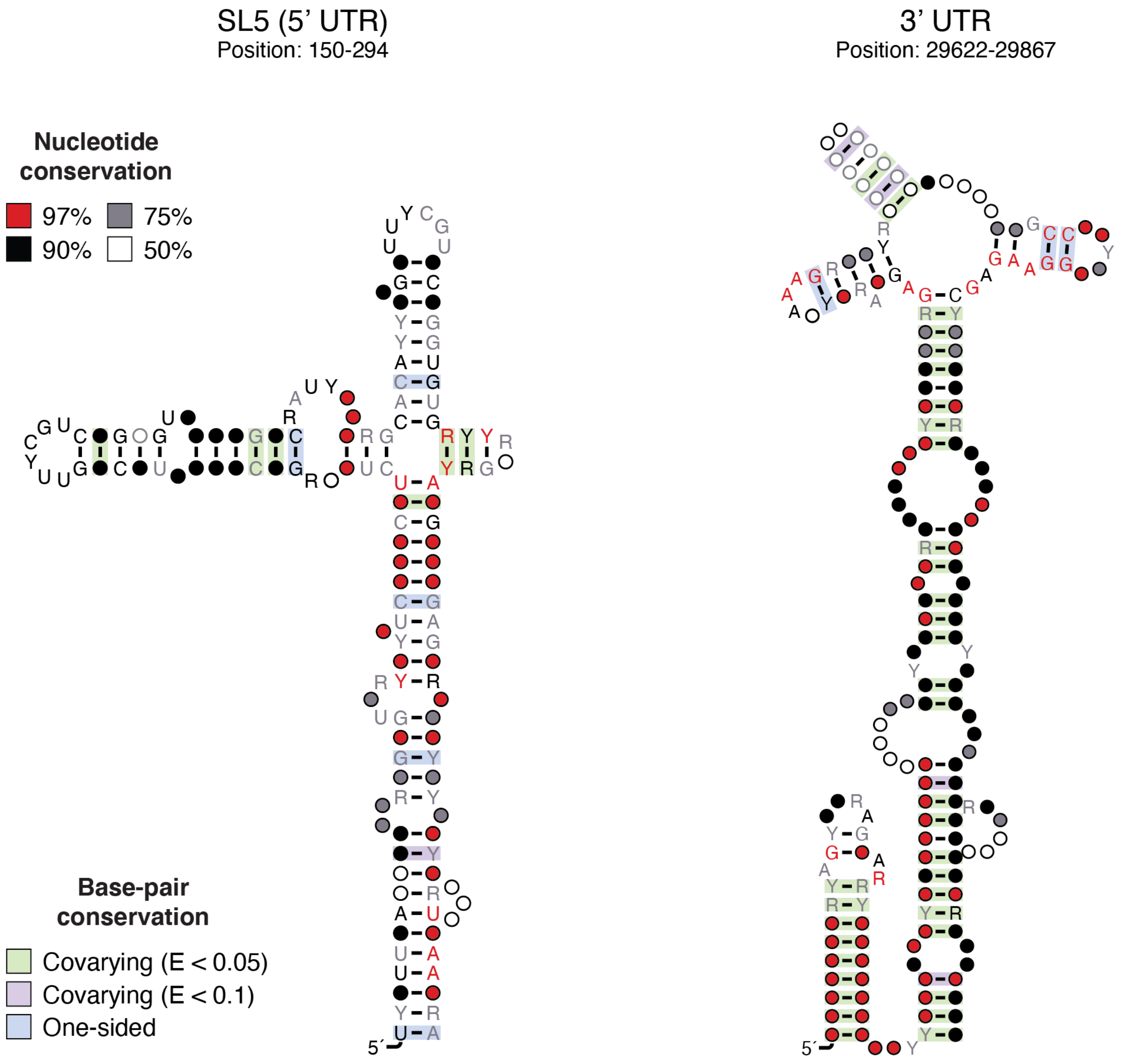


**Figure S60.** Structure model for the highly-conserved Sarbecovirus 5’ UTR’s SL5 four-way junction (*left*) and 3’ UTR (*right*). Models have been generated using the R2R software. Base-pairs showing significant covariation (as determined by R-scape) are boxed in green (E-value < 0.05) and violet (E-value < 0.1) respectively.

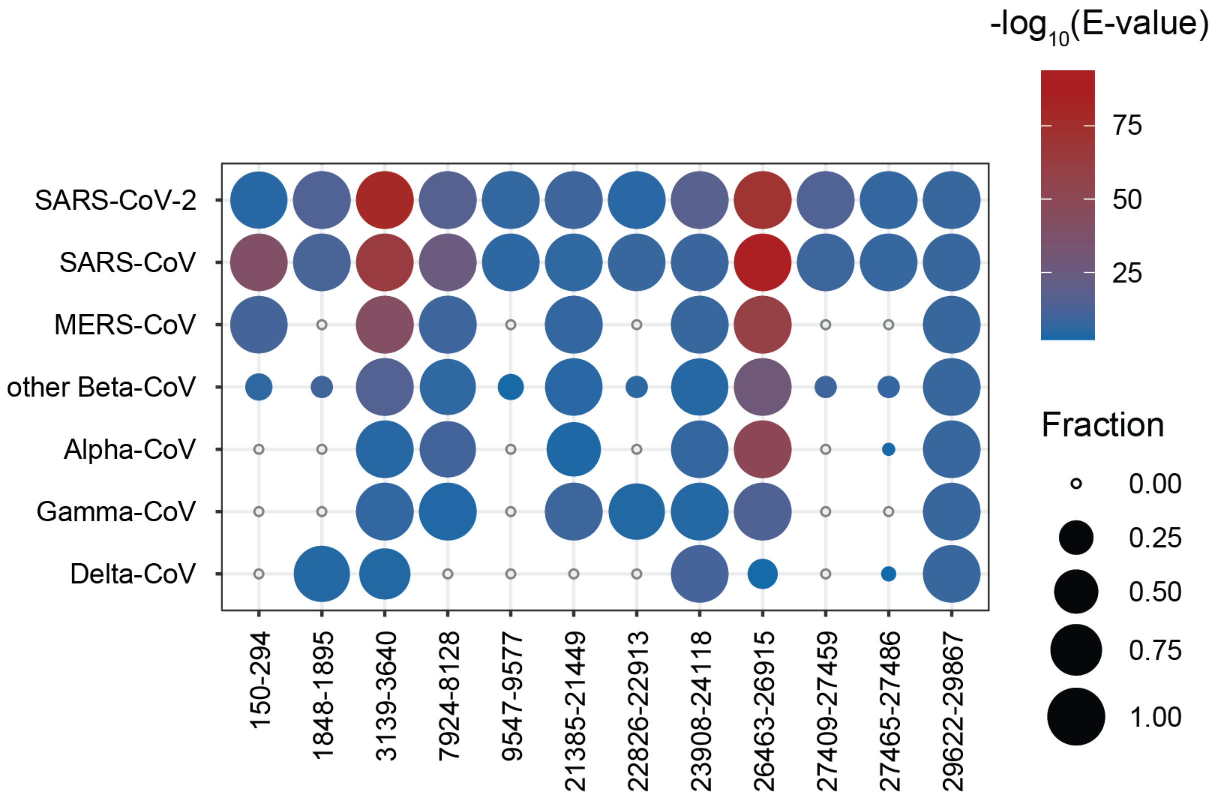


**Figure S61.** Heatmap showing the results for a search performed on a non-redundant database of 3,776 full-length coronavirus genomes using Infernal (E-value < 0.01) and the covariance models for the identified conserved SARS-CoV-2 structures. Circle diameter represents the fraction of sequences of a given coronavirus class, having a significant match for a given structure. Circles are colored according to the mean E-value.
